# Decellularized Normal and Tumor Extracellular Matrix as Scaffold for Cancer Organoid Cultures of Colorectal Peritoneal Metastases

**DOI:** 10.1101/2021.07.15.452437

**Authors:** Luca Varinelli, Marcello Guaglio, Silvia Brich, Susanna Zanutto, Antonino Belfiore, Federica Zanardi, Fabio Iannelli, Amanda Oldani, Elisa Costa, Matteo Chighizola, Ewelina Lorenc, Simone P. Minardi, Stefano Fortuzzi, Martina Filugelli, Giovanna Garzone, Federica Pisati, Manuela Vecchi, Giancarlo Pruneri, Kusamura Shigeki, Dario Baratti, Laura Cattaneo, Dario Parazzoli, Alessandro Podestà, Massimo Milione, Marcello Deraco, Marco A. Pierotti, Manuela Gariboldi

## Abstract

Peritoneal metastases (PM) from colorectal cancer (CRC) are associated with poor survival. The extracellular matrix (ECM) plays a fundamental role in modulating the homing of CRC metastases to the peritoneum. The mechanisms underlying the interactions between metastatic cells and the ECM, however, remain poorly understood and the number of *in vitro* models available for the study of the peritoneal metastatic process is limited. Here, we show that decellularized ECM of the peritoneal cavity allows the growth of organoids obtained from PM, favoring the development of three-dimensional nodules that maintain the characteristics of *in vivo* PM. Organoids preferentially grow on scaffolds obtained from neoplastic peritoneum, which are characterized by greater stiffness than normal scaffolds. A gene expression analysis of organoids grown on different substrates reflected faithfully the clinical and biological characteristics of the organoids. An impact of the ECM on the response to standard chemotherapy treatment for PM was also observed.

**Significance:** Evidence of the value of ex vivo 3D models obtained by combining patient-derived extracellular matrices depleted of cellular components and organoids to mimic the metastatic niche, to be used as a tool to develop new therapeutic strategies in a biologically relevant context, to personalize treatments and increase their efficacy.

## Introduction

The peritoneum is the second most common site of metastasis for colorectal cancer (CRC) after the liver [1]. In the past, peritoneal metastases (PM) were considered a terminal condition, amenable only to palliative treatments. The advent of cytoreductive surgery and hyperthermic intraperitoneal chemotherapy (HIPEC) in the 1990s have allowed some patients with PM to achieve longterm survival, pushing the median overall survival from 16 to 51 months [2]. However, about 70% of treated patients still experience peritoneal relapse [3]. The development of preclinical cellular models that faithfully recapitulate PM pathology, therefore, is crucial for the identification of more effective therapeutic strategies.

The peritoneal metastatic cascade consists of a series of steps that begin with cell detachment from the primary tumor [4]. Fine-tuned interactions between biochemical factors and biomechanical events, such as remodeling of the extracellular matrix (ECM), govern the cascade and allow the formation of the metastatic niche [5–8]. The metastatic niche facilitates organotrophic metastasis through the direct promotion of cancer stem cell survival, exploiting a tissue-specific microenvironment that is more suitable for the attachment of exfoliated neoplastic cells [5]. The biology behind these processes, however, is poorly understood due to the lack of organ-specific experimental models.

Most of the current data on metastatic spread have been obtained using cancer cell lines or patient-derived xenograft models, which do not fully reflect the physiopathology of their tumor of origin [9]. Tumor-derived organoids (TDO) are an intermediate model between cell lines and xenografts; they grow three-dimensionally and retain cell–cell and cell–matrix interactions, which more closely reflect the characteristics of the original tumor. Importantly, organoids can be established in a short time and are easy to manipulate [10]; they retain the genetic status of the original tissues and can be used to identify new therapeutic targets [11].

The possibility to isolate natural decellularized ECM and at the same time preserve both the 3D tissue architecture and the biochemical properties, enabling the development of more physiological cancer models [12, 13], prompted us to develop a tissue-engineered PM model for *in vitro* studies. Our model is based on the seeding of PM-derived organoids onto decellularized peritoneum-derived ECMs. Thanks to the possibility of characterizing the biochemical and biophysical properties of both organoids and ECM, the developed PM model allowed us to study the complex interactions between the ECM and neoplastic cells, by which we gained new insights into PM biology and the mechanisms underlying the cell-microenvironment interaction in this system.

## Materials and Methods

### Human tissues

Peritoneal tissue was collected from six patients with peritoneal metastatic colorectal carcinoma, who underwent surgical resection at the Peritoneal Malignancies Unit of our Institution. The patients were staged according to the WHO classification [14]. The study was approved by the Institutional review board (134/13; I249/19) and was conducted in accordance with the Declaration of Helsinki, 2009. Written informed consent was acquired. Metastatic lesions and apparently normal tissue (> 10 cm from the metastatic lesions) were harvested and one part of the metastatic tissue (1 cm in diameter) was placed in ice-cold PBS (ThermoFisher Scientific, Waltham, MA) containing gentamicin (50 ng/ml, ThermoFisher Scientific) and amphotericin B (50 ng/ml, ThermoFisher Scientific) for the generation of PM-derived organoids, while a second specimen was frozen in liquid nitrogen for molecular and histopathological analyses. The remaining tissue was used to develop 3D-decellularized matrices (3D-dECMs).

Normal tissue was partly used to develop 3D-decellularized matrices and partly frozen for further studies. FFPE blocks were prepared for IHC analyses of normal and metastatic tissue.

### Development of PM-derived organoids

Tumor-derived tissue was cut into small pieces, washed with ice-cold PBS at least ten times, and subsequently digested with 500 U/ml collagenase type II (Sigma Aldrich, St.Louis, Missouri, USA) in DMEM-F12 for 1 hour at 37 °C with vigorous pipetting every 15 minutes. The remaining fragments were digested with 1 mg/ml trypsin, 5 mM EDTA (ThermoFisher Scientific) at 37 °C for 20 minutes. The supernatant was collected and centrifuged at 300 g for 5 minutes at 4 °C. The cell pellet was resuspended with Matrigel, growth factor reduced (Corning, NY, USA), and dispensed into 24-well cell culture plates (50 μl/well). After Matrigel polymerization, the cells were overlaid with 500 μl of basal cell culture medium consisting of Advanced DMEM-F12 (ThermoFisher Scientific) and supplemented with different growth factors (Supplementary Table S1) to mimic different niche factor conditions (Fig. 1A), as in Fujii *et al* [10]. Incubation was performed at 20% O_2_ and 5% CO_2_. After expansion, the TDO were cultured in cell culture medium lacking growth factors, which was refreshed every three days. Optimal cell culture medium conditions were determined separately for each organoid culture (Supplementary Table S2).

**Fig. 1.**
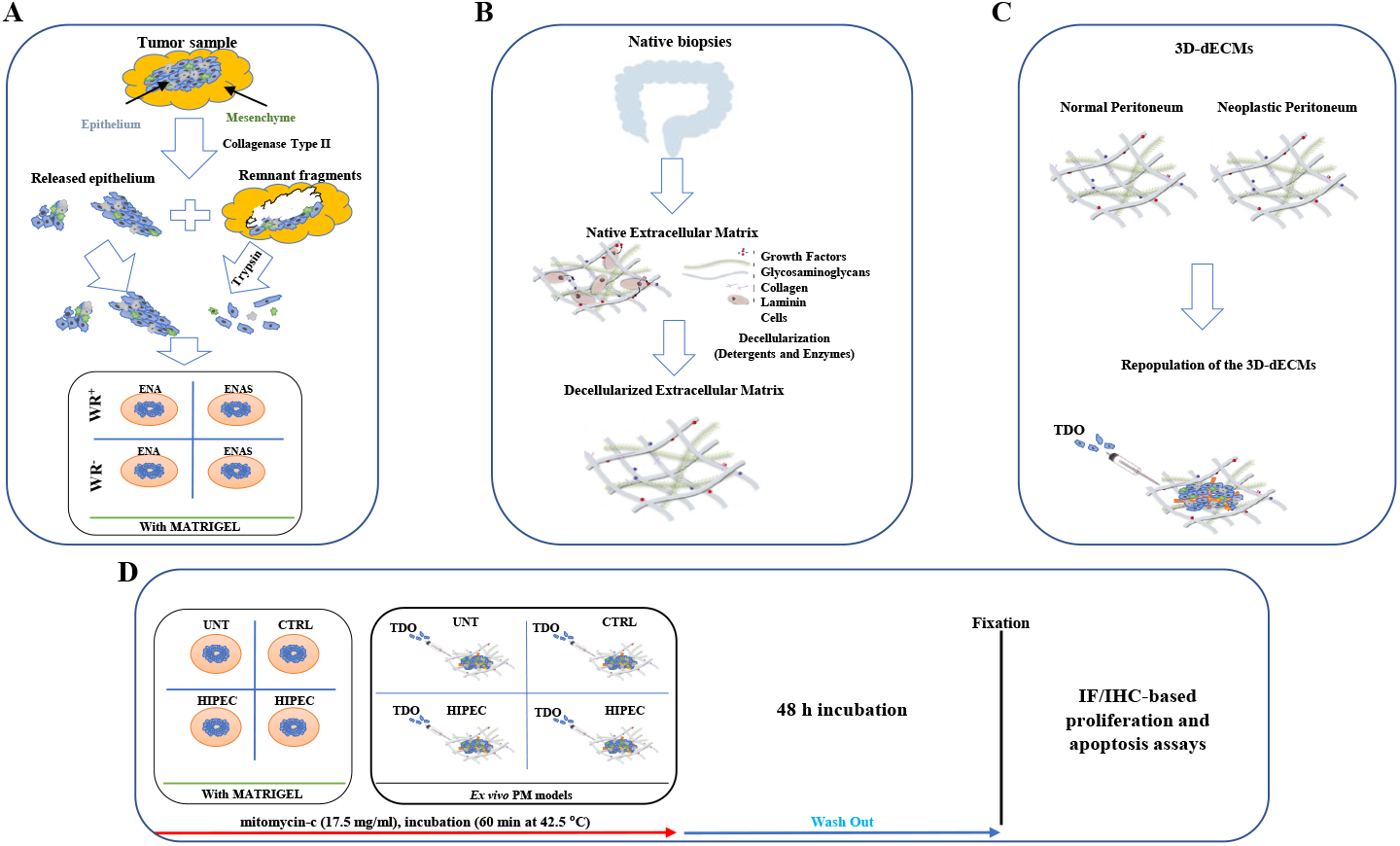
Diagram of established method. **(A)** Schematic representation of the protocol used to develop CRC PM-derived organoids. Digested cells were cultured in presence of different growth factors, to mimic different niche factors conditions. (**ENA:** human EGF recombinant protein (**E**); human Noggin recombinant protein (**N**); A83-01, anti-p38 inhibitor (**A**); **ENAS:** ENA supplemented with SB201290 anti-Rock inhibitor (**S**); **WR+/-:** cell culture media supplemented with or without human Wnt3A recombinant protein (**W**) and human R-Spondin-1 recombinant protein (**R**)). Organoids developed under a specific culture medium condition, depending on the growth factors they needed. **(B)** Schematic representation of the protocol used to obtain peritoneal-derived 3D-dECMs. **(C)** Experimental design used to develop the *ex vivo* 3D-engineered PM lesions. **(D)** Workflow chart representing the experimental design used to reproduce *in vitro* HIPEC treatments (**UNT:** untreated group; **CTRL:** control group (mitomycin-c vehicle: physiological solution); **HIPEC:** Hyperthermic Intraperitoneal Chemotherapy; **TDO:** tumor-derived organoid; **IF:** Immunofluorescence; **IHC:** Immunohistochemistry).

Organoids were split every 1-2 weeks as follows: they were mechanically removed from the Matrigel by pipetting, incubated in Cell Recovery Solution (Corning) for 1 hour at 4 °C, washed twice with ice-cold PBS and seeded as described above.

Aliquots of each organoid culture were frozen or prepared for IHC analyses as follows: samples were fixed in 10% formalin at room temperature (RT) for 10 minutes and embedded into 200 μl Bio-Agar (Bio-Optica, Milan, Italy). The samples were then cooled at −20 °C until solidification. For each sample, sections of 3 μm were obtained.

### Preparation of 3D-dECMs

3D-dECMs were derived from both PM and the corresponding normal peritoneum. Each experiment was conducted using three to ten different surgical specimens deriving from different patients.

The decellularization was performed as shown in Fig. 1B, according to the protocol described by Genovese et al. [13]. Briefly, both PM and normal peritoneum samples (60-100 mg wet weight) were kept in ice-cold PBS for 1 hour before processing. The specimens were split into two fragments, one of which was kept untreated for later comparison of the characteristics of the 3D-dECMs. For decellularization, the fragments were washed with ice-cold PBS supplemented with 50 ng/ml gentamicin and 50 ng/ml amphotericin B, followed by treatment with solutions containing detergents and enzymatic agents.

The success of the decellularization procedure was evaluated by analyzing the DNA content of the 3D-dECMs (see below). The 3D-dECMs were then washed with ice-cold PBS and either transferred into chilled freezing solution (90% DMEM-F12, 10% DMSO) and frozen for storage or fixed for IHC and immunofluorescence (IF) analyses. All the decellularization experiments were performed in triplicate, using at least three different samples, each derived from a different donor.

### *ex vivo* engineered PM lesion

Engineered PM lesions were obtained from three organoid cultures (C1, C2 and C3). TDO were removed from the Matrigel as described above and dissociated into single-cell suspensions with Trypsin-EDTA by vigorous pipetting for 10 minutes. About one million dissociated cells were counted with an automatic cell counter (ThermoFisher Scientific). 3D-dECMs derived from normal peritoneal tissue and PM were incubated overnight at 37 °C in DMEM-F12 supplemented with 10% FBS (Euroclone, Milan, Italy) and 50 ng/ml gentamicin and amphotericin B. To reduce intra-sample variability, the 3D-dECMs were cut into fragments of comparable sizes.

TDO were resuspended in 1 ml cell culture medium (Supplementary Table S1) and seeded on the top of 50 mg of 3D-dECMs. Repopulated matrices were placed in a 24-well cell culture plate (Corning) containing DMEM-F12 supplemented with 10% FBS and 50 ng/ml gentamicin and amphotericin B, followed by incubation for 2 hours at 37 °C. Each well was filled with 2 ml cell culture medium, which was changed every two days (Fig. 1C). Repopulated matrices were either frozen for RNA extraction or fixed for IHC and IF analyses. Representative 3 μm FFPE sections were cut at different depths to verify the presence of TDO in the inner part of the 3D-dECM scaffold. The repopulation experiments were performed in triplicate, using three different neoplastic and normal-derived matrices obtained from three different donors.

### Nucleic acids extraction

DNA from FFPE sections of the PM-derived organoids and their tissue of origin was used for mutational analysis. DNA was extracted using the Masterpure Complete DNA Purification Kit (Lucigen-Biosearch Technologies, Middleton, WI, USA) and quantified on the QIAxpert^®^ spectrophotometer (QIAGEN, Hilden, Germany).

DNA from 20 mg of normal peritoneum and PMs, both decellularized and untreated, was used to evaluate the success of the decellularization procedure. DNA was extracted using the DNeasy Blood&Tissue kit (QIAGEN) according to the manufacturer’s instructions, and quantified using Nanodrop 1000 (ThermoFisher Scientific) at 260/280 nm ratio. DNA from decellularized ECM, normal peritoneum, PM, and their corresponding non-decellularized samples was loaded onto a 1% agarose gel. The separated bands were visualized by exposing the gel to UV light and images were acquired using Gel Doc (Bio-Rad, Hercules, CA, USA). All of the experiments were performed in triplicate.

RNA from the three organoid cultures (C1, C2 and C3) grown both in Matrigel and on normal or neoplastic peritoneal 3D-dECMs was used for RNA-seq analyses. For the PM organoids, the Matrigel was digested with Cell Recovery Solution (Corning) as described above. The pellet was washed three times with ice-cold PBS and suspended in 1 ml TRIzol™ reagent (QIAGEN). Instead, for the repopulated 3D-dECMs, the matrices were washed three times with ice-cold PBS and homogenized using the TISSUE Tearor Homogenizer (QIAGEN) in 500 μl TRIzol™ reagent (QIAGEN). Then, RNA was extracted following the manufacturer’s instructions, quantified on a ND-1000 spectrophotometer (ThermoFisher Scientific) and stored at −80 °C.

RNA from FFPE sections (10 μm) of the PMs from which the six TDO were derived was used to validate the results from the RNA-seq analysis. RNA was extracted using the miRNeasy FFPE kit (QIAGEN) and quantified on a NanoDrop™ 1000 (ThermoFisher Scientific, Waltham, MA).

### Histochemistry (HC), IHC and IF

Before HC and IHC staining, FFPE sections were cut into slices and dewaxed in xylene, rehydrated through decreasing concentrations of ethanol and washed with water. Slices were stained with H&E for quality control. For HC analysis, sections were stained with Masson’s trichrome (Aniline blue kit; Bio-Optica), Alcian blue stain (pH 2.5 kit, Bio-Optica), van Gieson trichrome (Bio-Optica), and Periodic Acid Schiff (PAS, Bio-Optica) following the manufacturers’ instructions. IHC was performed using the following mouse anti-human antibodies: Ki-67, CK19, CK20, CK AE1/AE3, CDX2, LGR5, vimentin, YAP and TAZ. Images were acquired with a DM6000B microscope (Leica). Staining for Ki-67, CK19, CK20, CK AE1/AE3, CDX2, LGR5 and vimentin antibodies was performed automatically using theAutostainer Link 48 (Dako, Agilent, Santa Clara, CA. US). Antigen retrieval for YAP and TAZ antibodies was carried out using preheated target retrieval solution (pH 6.0) for 30 minutes. Tissue sections were blocked with FBS serum in PBS for 60 min and incubated overnight with primary antibodies. The antibody binding was detected using a polymer detection kit (GAM/GAR-HRP, Microtech) followed by a diaminobenzidine chromogen reaction (Peroxidase substrate kit, DAB, SK-4100; Vector Lab). All sections were counterstained with Mayer’s hematoxylin. Dilutions and experimental conditions are listed in Supplementary Table S3. For IF analyses, FFPE sections were stained with Alexa680-conjugated Wheat Germ Agglutinin (WGA) marker (ThermoFisher Scientific) and DAPI (VECTASHIELD Mounting Medium with DAPI, Maravai LifeScencies, San Diego, CA, USA), with anti-human Ki-67 and LGR5 monoclonal antibodies, with anti-human Collagen-IV and anti-human monoclonal cCASPASE3 antibodies and DAPI, followed by Alexa488-conjugate goat anti-mouse or Alexa546-conjugated goat anti-rabbit IgG polyclonal secondary antibodies for 1 hour at RT in dark (ThermoFisher Scientific). Images were acquired with a DM6000B microscope (Wetzlar, Germany Leica,) equipped with a 100 W mercury lamp, and analyzed using Cytovision software (Leica). Dilutions and experimental conditions are listed in Supplementary Table S3.

### DNA sequencing

About 150–200 ng genomic DNA (measured with Qubit dsDNA HS assay kit, ThermoFisher Scientific), were sheared by the Sure Select Enzymatic Fragmentation kit (Agilent Technologies Inc., Santa Clara, CA, USA). NGS libraries were created using Sure Select XT2 Low input Custom library probes (Agilent Technologies Inc.). The probe set was custom designed by Cogentech (OncoPan panel) and includes the exonic regions of the following genes: APC, ATM, BARD1, BMPR1A, BRCA1, BRCA2, BRIP1, CDH1, CDKN2A (α and β isoform), CDK4 (exon 2), CHEK2, CTNNA1, EPCAM, FANCM, MLH1, MSH2, MSH3, MSH6, MUTYH, NBN, NHTL1, PALB2, PMS2, POLD1, POLE, PTEN, RAD51C, RAD51D, SMAD4, STK11, TP53, KRAS, NRAS, BRAF, EGFR, HER2 (ERBB2), and PIK3CA. Sequencing was performed on Illumina MiSeq platform, in PE mode (2 x 150 bp). Raw reads were demultiplexed and aligned to a reference genome (Human GRCh37) using a pipeline developed in-house in collaboration with enGenome Software Company and annotated with the eVai tool. Results were compared to find the percentage of common SNVs (Single Nucletode Variants). Five PM-derived organoids (C1, C2, C3, C4 and C6) and their corresponding surgical samples were analyzed. FFPE tissue for C5, unfortunately, was not available.

### Morphological evaluation of the decellularized matrices

3D-dECMs from normal peritoneum and PM lesions were washed twice with 1X PBS and placed in a 60 mm petri dish. Samples were illuminated with a widefield lamp laser to visualize the architecture of the collagen fibers. An image format of 1024×1024 pixels was used and all images were acquired with Leica Application Suite X, ver. 3 software. 3D-dECMs FFPE sections deriving from normal and PM peritoneum were used to perform polarized light microscopy (PLM). FFPEs were analyzed with an Olympus BX63 upright widefield microscope equipped with a motorized stage and a Hamamatsu OrcaAG camera, using Metamorph software. UplanSApo 4X/0.16 N.A objective was used to acquire the mosaics of the sections. Insets were acquired with UplanSApo 10X/0.4 N.A. and UplanSApo 20X/0.75 N.A. objectives. All experiments were perfomed at least in duplicate. Confocal reflection microscopy images were acquired with a Leica TCS SP8 laser confocal scanner mounted on a Leica DMi8 microscope through a HC PL FLUOTAR 20×/0.5 NA.

### Nanoscale topographical analysis of 3D-dECMs

The topographical evaluation of the 3D-dECMs was performed by atomic force microscopy (AFM) analysis on samples deriving from normal peritoneum and PM of three different patients. Before the AFM analysis, the 3D-dECM slides were left for 30 minutes at RT to dissolve the optimal cutting temperature (OCT) compound. Then, the samples were carefully washed with ultrapure water and covered with 1X PBS buffer. AFM topographic measurements were carried out at RT using a NanoWizard3 AFM (JPK, Germany) coupled to an Olympus BX61 inverted microscope and equipped with tapping mode silicon ACTG AFM probes (APPNANO). The 50 μm thick tissue slices, instead, were mounted on polarized glass slides (ThermoFisher Scientific), left for 30 minutes at RT and carefully washed with ultrapure water. The topography of each tissue was characterized by collecting at least 10 areas (5×5 μm^2^) of the sample surface with 512×512 points (scan speed 3,5 μm s^-1^).

### ECM component quantification

Total collagen and sulphated glycosaminoglycan (sGAG) content in fresh and decellularized normal and PM peritoneum were quantified using the SIRCOL collagen assay (Biocolor, Carrickfergus, UK) and the Blyscan GAG assay kit (Biocolor), respectively. The experiments were performed in triplicate following the manufacturer’s instruction. Data are the mean of three different neoplastic and normal-derived samples obtained from three different donors.

### Nanoindentation measurements by AFM

AFM mechanical analysis was carried out on 3D-dECMs deriving from normal peritoneum and PM of five patients (see Fig. 2). 3D-dECMs were embedded in OCT and frozen with nitrogen-cooled 2-propanol for 10 seconds. Slices of 100 μm thickness were cut with a microtome (Leica) and attached to positively charged poly-lysine coated glass coverslips (ThermoFisher Scientific), exploiting the electrostatic interaction. Nanomechanical tests were performed in liquid on samples covered by a PBS droplet confined by a circular ridge of hydrophobic two-component silicone paste (Leica). A Bioscope Catalyst AFM (Bruker) was used, which was resting on an active anti-vibration base (DVIA-T45, Daeil Systems) and put into an acoustic enclosure (Schaefer). The measurements were performed at RT.

**Fig. 2.**
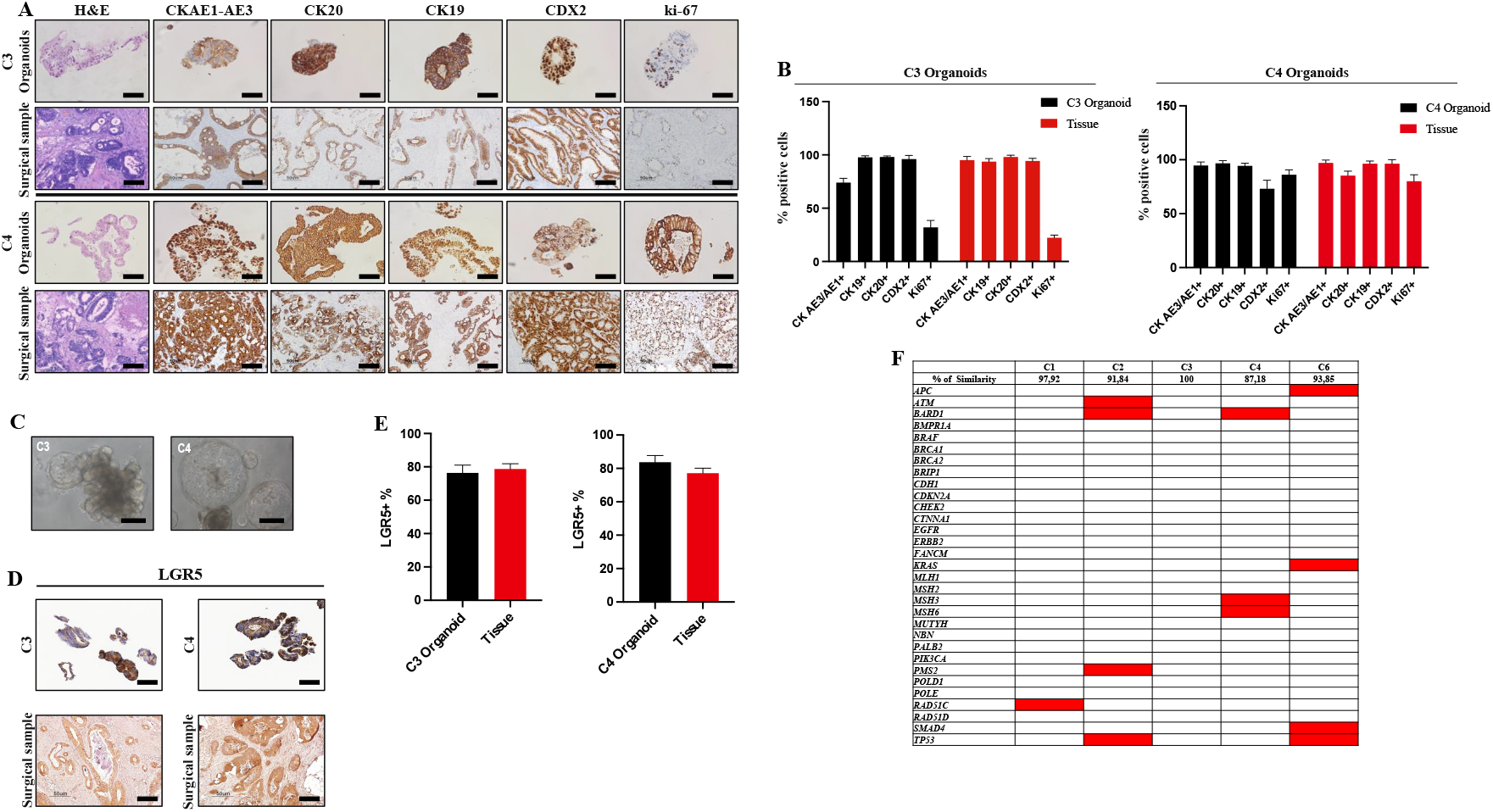
Establishment and characterization of human peritoneal metastases (PM)-derived organoids. **(A)** Comparative histological and IHC analysis of PM-derived organoids and their tissue of origin, using H&E staining and CK AE1/AE3, CK20, CK19, CDX2, and Ki-67 immunostaining. Organoids retain the main features of their tissue of origin. PM tissues generally present a tumor epithelium surrounded by stromal-derived cells, while organoids consist exclusively of epithelial cells. Scale bar: 100 μm. **(B)** Quantitative counts of the percentage of CK AE/AE3, CK19, CK20, CDX2 and Ki-67 positive cells in C3 and C4 organoids Vs their corresponding tumor of origin. Data are presented as median and SD of three fields per experiments, counted using Qpath software. One-way ANOVA did not show differences between the two groups. **(C)** Bright-field images depicting organoid phenotypes. The left micrograph shows a glandular-like branched organoid, while the right one shows a spherical-like cohesive organoid. Scale bar: 100 μm. **(D)** IHC analysis of organoids (top) and their tissue of origin (bottom), using LGR5 immunostaining. LGR5-positive cells found in PM tissue are retained in the derived organoids. Scale bar: 100 μm. **(E)** Quantitative counts of the percentage of LGR5 positive cells in C3 and C4 organoids Vs their corresponding tumor of origin. Data are presented as median and SD of three fields per experiments, counted using Qpath software. One-way ANOVA did not show differences between the two groups. **(F)** Summary of cancer-related genes, analyzed by target DNA sequencing, with acquired mutations in TDO with respect to their tumor of origin (red boxes). The percentage of similarity was reported. Passage numbers of the organoid lines were: C1: P11; C2: P13; C3: P10; C4: P14; C6: P10.

Custom monolithic borosilicate glass probes consisting of spherical glass beads (SPI Supplies), with radii R in the range of 7.5–12.5 μm, were attached to tipless cantilevers (Nanosensor, TL-FM) with nominal spring constant k = 3-6 N/m. Probes were fabricated and calibrated, in terms of tip radius, according to an established custom protocol [15]. The spring constant was measured using the thermal noise calibration [16, 17] and corrected for the contribution of the added mass of the sphere [18, 19]. The deflection sensitivity was calibrated *in situ* and non-invasively before every experiment by using the previously characterized spring constant as a reference, according to the SNAP procedure described in [20].

The mechanical properties of the 3D-dECMs were obtained by fitting the Hertz model to sets of force versus indentation curves (simply force curves, FCs), as described elsewhere [20, 21, 22, 23], to exctract the value of the YM of elasticity, which measures ECM rigidity. FCs were collected in Point and Shoot (P&S) mode, selecting the regions of interest from optical images, exploiting the accurate alignment of the optical and AFM images obtained using the Miro software module integrated in the AFM software. Each set of FCs consisted of an array of typically 15×15=225 FCs spatially separated by 5-10 μm, each FC containing 8192 points, with ramp length L = 8-15 μm, maximum load Fmax = 150-1500 nN, and ramp frequency f = 1 Hz. The maximum load was chosen in order to achieve a maximum indentation in the range of 4-9 μm. Typical approaching speed of the probe during indentation was 16-30 μm/s.

Five samples were characterized for each condition. In each sample, 3-10 P&S were acquired in macroscopically separated locations, for a total of 10-25 independent P&S per patient and condition (up to 2250-5500 FCs per patient and condition).

### Stem cell maintenance, proliferation and apoptosis assays

Growing cells, stem cells and apoptotic cells were detected on FFPE sections. Growing cells, deriving from disaggregated TDO, were stained with anti-human Ki-67 monoclonal antibody (clone MIB-1) and DAPI, and their growth rate was expressed as the percentage of Ki-67-positive cells present in fields devoid of dead cells. Stem cells were stained with anti-human LGR5 monoclonal antibody (clone OTI2A2) and DAPI, and their density was expressed as the percentage of LGR5-positive cells present in fields devoid of dead cells. Apoptotic cells were stained with anti-human cCASPASE3 monoclonal antibody (clone 9661) and DAPI, and the apoptotic rate was calculated as the percentage of cCASPASE3-positive cells present in the field. The percentage of Ki-67-positive, LGR5-positive and cCASPASE3-positive cells was obtained by dividing the number of positive cell present in one field by the total number of cells in one field, multiplied by 100. Cells in three independent fields (40X magnification) were counted using ImageJ software. The experiments were performed in triplicate using three different neoplastic and normal-derived matrices obtained from three different donors.

### Qpath analyses

Percentage estimation and cell counting were performed using Qupath software (https://qupath.github.io, version 0.2.3). The images used for Qpath analyses were acquired using Aperio Leica ScanScope XT (Leica Biosystems, Wetzlar, Germany). The slides were evaluated by an expert pathologist. The percentage of CK AE1/AE3, CK20, CK19, CDX2, Ki-67, and LGR5 positive cells was calculated by dividing the number of positive cells present in each field by the total number of cells in the same field. TDO-derived infiltrating cells were evaluated by calculating the total number of H&E stained cells. Three fields were counted per experiments.

### RNA-seq analysis

Gene expression profiles were conducted on C1, C2 and C3 organoid cultures grown in Matrigel and on 3D-dECMs. Total RNA was extracted using TRIzol™ reagent (QIAGEN). Qubit fluorimeter (ThermoFisher Scientific) and Agilent Bioanalyzer 2100 (RIN > 8) were used to measure and assess RNA abundance and integrity, respectively. Indexed library preparation was performed starting with 500 ng total RNA with the TruSeq stranded mRNA (Illumina) according to the manufacturer’s instructions. RNA-seq was performed in PE mode (2×75nt) on an Illumina NextSeq550 platform, generating an average of 55 million PE reads per sample. For every condition (Matrigel, normal 3D-dCM and neoplastic 3D-dECM), two replicates per organoid were sequenced, for a total of 18 data points. Raw reads were aligned to the human transcriptome (hg38) with STAR [24] using the quantMode option to generate transcripts counts. Differentially expressed genes in the three growth conditions were identified with DESeq2 [25]. All *p*-values were adjusted for false discovery rate with the Benjamini-Hochberg method.

### Gene Set Enrichment Analysis

Gene Set Enrichment Analysis was performed with the enrichR R package [26] on deregulated genes (absolute fold change > 2 and adjusted *p*-value <0.05). In particular, the enrichment for the Matrisome database was assessed. This database provides live cross-referencing to gene and protein databases for every ECM and ECM-associated gene, also integrating experimental proteomic data on ECM and ECM-associated proteins and genes from the ECM Atlas [27]. Gene sets with adjusted *p*-value <0.05 were considered significantly enriched.

### Quantitative real-time polymerase chain reaction (qRT-PCR)

For gene expression analysis, cDNA was synthesized from 100 ng of total RNA using a High-Capacity cDNA Reverse Transcription Kit (ThermoFisher Scientific, Waltham, MA) and qPCR was carried out with gene-specific assays for MT1A (Hs00831826_s1), LOX (Hs00942480_m1), THY1 (Hs00174816_m1), FZD9 (Hs00268954_s1), SPP1 (Hs00959010_m1), and performed using the TaqMan FAST Universal PCR Master Mix, no AmpErase® UNG in a PRISM 7900HT Real-Time PCR system (Thermo Fisher Scientific). The expression values of the genes were normalized to GAPDH (Hs99999905_m1).

### Treatment with cytotoxic drugs

Mitomycin-c (MMC) (Kyowa Kirin Co., Ltd., Tokyo, Japan) and oxaliplatin (OXA) (Fresenius Kabi, Bad Homburg, Germany) were used for the *in vitro* simulation of HIPEC treatment. MMC was dissolved in DMSO to obtain a 60 mM stock solution. OXA was diluted in physiological solution (0.45 % sodium chloride and 2.5 % glucose) to obtain a 15 mM stock solution. Both drugs were diluited to the working concentration in the cell culture medium, where the final solvent concentration was <0.1% for all samples, including controls. The experiments were performed in triplicate, using three different neoplastic and normal-derived matrices obtained from three different donors.

### Dose-response curves for HIPEC treatment

To determine the IC_50_ value of MMC and OXA, 5×10^3^ C1, C2 and C3 TDO were suspended in 100 μl of culture medium and seeded on 96-well plates (Costar 3904; Corning, New York, USA) coated with 40 μl of Matrigel. TDo were dispensed on the top of the matrigel. After two days, TDO were incubated with 100 μl preheated drug at concentrations ranging from 2.5 to 200 μM for MMC and between 10 and 700 μM for OXA, for 60 min (MMC) or 90 min (OXA) at 42.5 °C. The values were chosen by scaling up and down the concentrations used for patients (35 mg/m^2^ for MMC and 200 mg/m for OXA, which correspond to 41.9μM for MMC and 252 μM for OXA for *in vitro* treatments [28]). TDO viability, was assessed using a CellTiterGlo® 2.0 kit (Promega, Fitchburg, Wisconsin, USA) on a TECAN spark microplate reader (Tecan Trading AG, Switzerland). Viability was normalized to the mean of three control samples/plate (TDO treated with 0.5 % DMSO in MMC and physiological solution in OXA experiments). All the experiments were performed in triplicate.

### *ex vivo* PM lesion to test HIPEC efficacy *in vitro*

An *ex vivo* engineered micrometastasis that reproduces the binding of CRC circulating metastatic cells to the peritoneum was obtained by growing PM-derived organoids on 3D-dECMs from neoplastic peritoneum. In the model, PM-derived organoids were in contact with the drug, as during HIPEC. Briefly, PM-derived organoids were grown on the top of neoplastic 3D-dECMs in a 24-well cell culture plate for 12 days in order to allow a complete colonization of the matrix [12, 13, 29]. TDO grown in Matrigel were used as control to evaluate the impact of native 3D-dECMs on the HIPEC treatment. The engineered PM lesions were treated with preheated MMC and OXA at a concentration of 41.9 μM for 60 min at 42.5 °C and 252 μM for 90 min at 42.5 °C respectively, corresponding to the calculated clinical concentrations, which is in line with the current standard protocols used for HIPEC at Fondazione IRCCS Istituto Nazionale dei Tumori - Milano Institution. Whereafter the PM models were subjected to three washes with 1X PBS and incubated for 48 hours with appropriate cell growth medium (Fig. 1D).

Samples were fixed in formalin and FFPE sections were obtained as described above. The impact of HIPEC treatment on TDO proliferation and the activation of an apoptotic program were determined by Ki-67 and cCASPASE3 immunostaining, respectively. All the experiments were performed in triplicate using three different neoplastic and normal-derived matrices obtained from three different donors.

### Immunoblotting

After HIPEC simulation using drugs at a concentration corresponding to the IC_50_ value of each TDO, C1, C2, and C3 TDO were lysed in Ripa buffer (50 mM Tris, pH 8.0, 50 mM NaCl, 0.5% Triton X-100, 0.1% sodium deoxycholate, 0.25% sodium dodecyl sulphate [SDS]) supplemented with protease inhibitors (Merck Millipore, Billerica, MA, USA) and a phosphatase inhibitor cocktail (Sigma-Aldrich, St. Louis, MO, USA), for 3 h at 4 °C on a rotation wheel. Samples were then sonicated and their protein content was quantified using the Bradford protein assay (Bio-Rad, Hercules, CA, USA). For each sample, 40 μg of protein extract were separated on 4–12% polyacrylamide gels, transferred onto nitrocellulose membranes (Sigma-Aldrich, St. Louis, MO, USA) and incubated with primary antibodies (Supplementary Table S4). The signals were detected using enhanced chemiluminescence, and protein levels were quantified using Imagelab software (Bio-Rad, Hercules, CA, USA). Each experiment was repeated at least three times.

### Statistical analyses

Statistical analyses were performed using GraphPad Prism software (version 8.4.1 (676), GraphPad Software, San Diego, USA). Data are expressed as mean and SEM. A two-tailed Student’s *t* test was used to compare paired groups. Differences among groups were evaluated using two-way ANOVA.

In the case of AFM mechanical experiments, for each patient and each condition tested, the median values of the YM were extracted from each measured location (P&S) using the procedure described in Cramer et al [23, 30]. The distributions of the measured YM values were obtained by grouping all P&S measured in all locations, for each patient and each condition tested. The mean and median values and the corresponding standard deviations of the mean (as SEM) were calculated by averaging between P&S. The statistical significance of differences between normal and neoplastic conditions was estimated by applying the two-tailed *t* test. A *p*-value <0.05 was considered statistically significant.

## Results

### Development of PM-derived organoids

Six organoid cultures (C1 to C6) were developed from PM, following established protocols [10, 31], as detailed in the Materials and Methods section. The main characteristics of the patients from which the organoids were derived are summarized in Supplementary Table S5. In line with previous works on organoids from advanced CRC [10, 31], the growth of the organoids required supplementation of minimal niche factors. Organoids carrying mutations in the KRAS gene required EGF or noggin-1 supplementation, while organoids carrying FGR1 amplification grew in medium with EGF. BRAF-mutant TDO grew in a medium with minimal requirements for niche factors. These results highlight how the different mutational profiles lead to specific requests for niche factors [10] (Supplementary Table S1).

### PM-derived organoid characterization

PM-derived organoids retained the main characteristics of their tissues of origin, expressing the specific colorectal markers CK AE1/AE3, CK19, CK20 and CDX2 in the same percentage of cells as the tissue from which they originated (Fig. 2A and 2B Supplementary Fig. S1A, S1C and S1E). The TDO exhibited the typical glandular features observed in the corresponding surgical sample, including signet-ring cells, nest-like growth pattern, nuclear atypia, cuboidal nuclear morphology and pleomorphism (Fig. 2A and 2C).

All organoids and corresponding clinical samples were positive for the Leucine-rich repeat-containing G-protein coupled receptor 5 (LGR5), the well-established stem cell marker for the colonic niche [11] (Fig. 2C, Supplementary Fig. S1B and S1D). The percentage of LGR5 positive cells was similar to that of their tissue of origin (Fig. 2E and Supplementary Fig S1F). The organoids were also Ki-67-positive, suggesting that they underwent active proliferation (Fig. 2A, Supplementary Fig. S1A and S1C). Moreover, the TDO mutational profile was very similar to that of the tumor of origin (Fig. 2F, Supplementary Fig. S1G).

### Development of 3D-dECMs

3D-dECMs from PM and normal peritoneum were generated following the protocol described by Genovese et al. [13]. DNA quantification showed almost total DNA depletion after the decellularization procedure (\*\*\**p*<0.001; Fig. 3A Supplementary Fig. S2A). Fluorescence analysis of formalin-fixed, paraffin-embedded (FFPE) sections highlighted the complete removal of the nuclei and lipids form the cellular membranes (Fig. 3B). Immunohystochemistry (IHC) analysis showed loss of cytokeratins and vimentin, indicating the absence of epithelial and mesenchymal cells, respectively, and the maintenance of collagen IV distribution, the main structural protein (Fig. 3C). Hematoxilin and Eosin (H&E), van Gienson, Masson trichrome and Alcian-PAS stainings revealed the maintainance of the architectural structure of the corresponding non-decellularized samples (Fig. 3C and 3D).

**Fig. 3.**
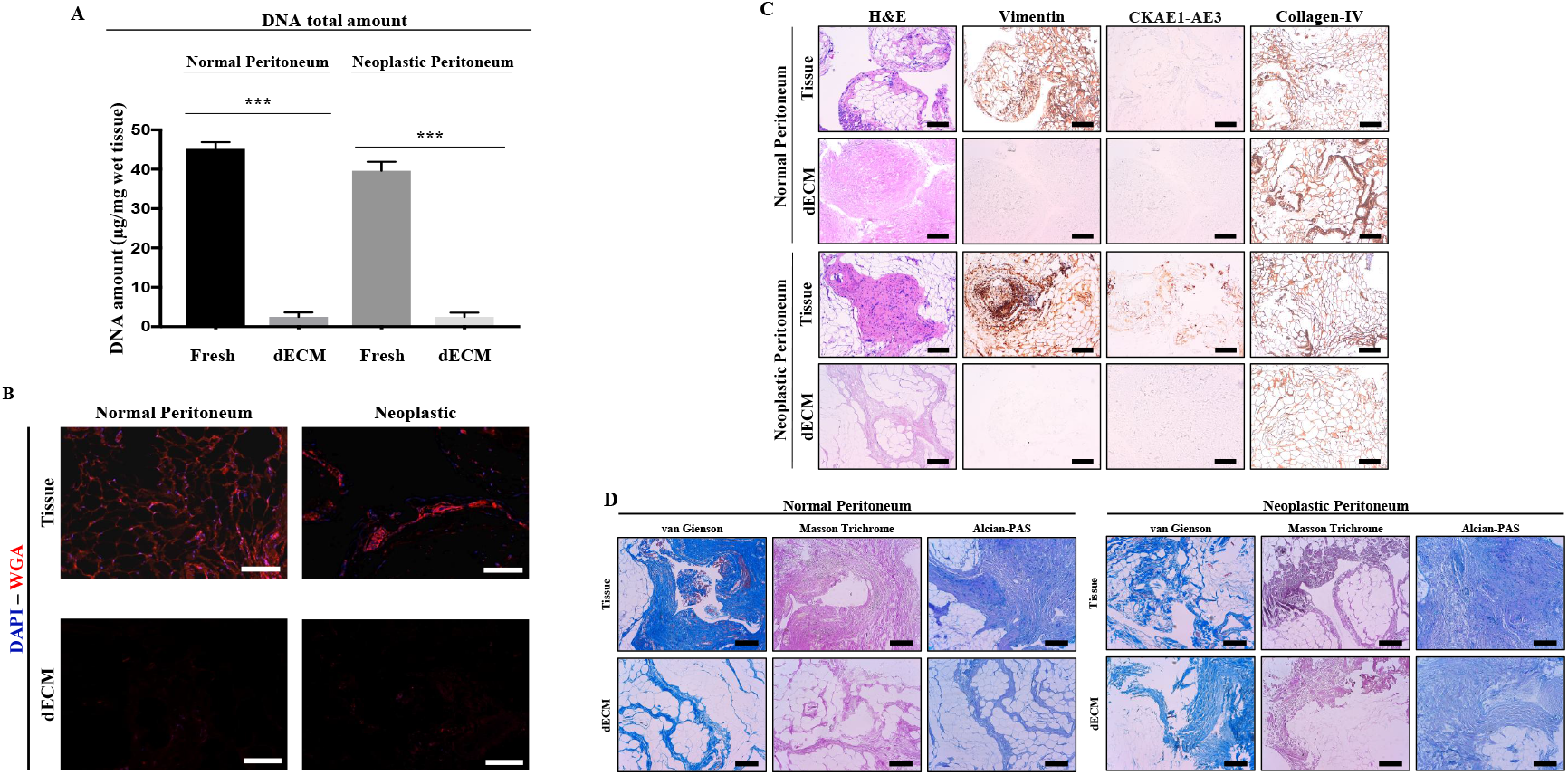
Establishment of 3D-dECM scaffolds from peritoneal cavity. **(A)** DNA quantification of normal and neoplastic peritoneal tissue samples before (fresh) and after the decellularization treatment (dECM), showing a high DNA depletion. Student’s *t-test* (\*\*\**p*<0.001). **(B)** IF analysis of normal and neoplastic peritoneal samples before and after the decellularization procedure using the WGA (Wheat Germ Agglutinin, a marker for glycoproteins in lipid membranes; red) antibody. The samples were counterstained with DAPI (blue). Scale bar: 100 μm. **(C)** IHC analysis of fresh peritoneal-derived tissues and the corresponding decellularized samples using H&E staining and vimentin, pan-Cytokeratin (CK AE1/AE3), and collagen-IV immunostaining. Scale bar: 200 μm. **(D)** Van Gienson, Masson’s Trichrome and Alcian-PAS stains for the detection of collagens, polysaccharides, glycoproteins, and structural tissue preservation on fresh and decellularized peritoneal-derived samples. Scale bar: 200 μm.

Finally, lyophilyzed 3D-dECMs were added to the culture media and no differences were observed in cell viability after 72 hours of growth, indicating that the decellularization procedure has no cytotoxic effects (Supplementary Fig. S2B).

### 3D-dEMCs: morphological features and mechanical properties

Confocal reflection and polarized light microscopy techniques showed that the differences between normal and neoplastic-derived tissues obtained from three different PM patients were related both to the tissue texture and to the distribution and integrity of the single collagen fiber (Fig. 4A). The 3D-dECMs of normal tissues showed higher density and clusters of collagen fibers, with random relative orientation; in the neoplastic 3D-dECMs, the collagen structure appears more irregular and porous, although a tendency for fiber alignment and bundling on a larger scale can be observed. Atomic Force Microscopy (AFM) topographical analysis revealed an asymmetric distribution of collagen at the micrometric scale: matrices derived from normal tissues are organized in groups of very thin, intersecting fibers (with diameters below 50nm), characterized by a variety of orientations, while matrices derived from neoplastic tissues have a more corrugated and disordered surface pattern (Fig. 4B).

**Fig. 4.**
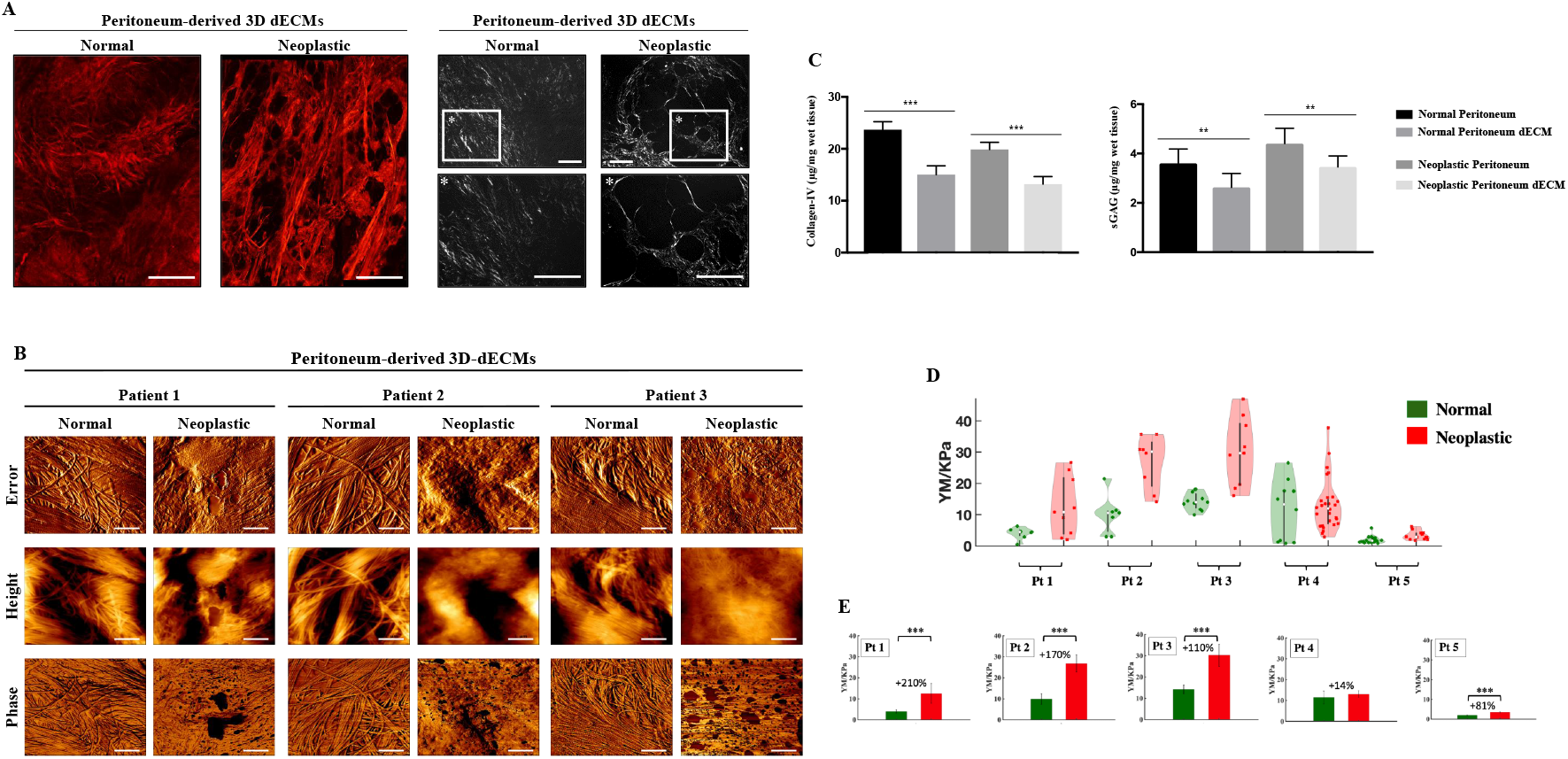
Morphological and mechanical properties of PM-derived 3D-dECMs. **(A)** Confocal and polarized light microscopy analysis of peritoneal-derived 3D-dECMs samples. Scale bar: 50 μm. **(B)** Topography analysis of peritoneal-derived 3D-dECMs. Phase, height and peak force error images of both normal and neoplastic decellularized matrices are shown. Scale bar: 1 μm. **(C)** Quantification of collagens I/IV and sulphated glycosaminoglycans (sGAG) on fresh and decellularized peritoneal tissues. Student’s *t-test* (\*\*\**p*<0.001 and \*\**p*<0.01). **(D)** Distribution of the YM values obtained for each patient (Pt) and condition (normal and neoplastic). Violin-plots: each dot represents the median YM value extracted from a single measurement (P&S) made approximately by 225 FCs; **(E)** Result of the statistical analysis of the YM value for each patient and condition tested. The bars and error bars represent mean of the median YM values and effective standard deviation of the mean, calculated as described in Materials and Methods. The percentage represent the relative stiffening of the neoplastic ECM.

At odd with the case of ECM derived from healthy and CRC-affected tissues [21], we do not observe a clear tendency for collagen fibers to form aligned bundles in the neoplastic state, although at this small scale the neoplastic matrix appears structurally more compact than the normal ECM. The increased anisotropy of the ECM structure reported in Nebuloni et al. [21] in the case of CRC is more evident in the optical confocal image of the neoplastic sample (Fig. 4A). Collagen was less abundant in neoplastic tissue than in normal tissue, however, the latter showed higher levels of glycosaminoglycans GAGs. Both collagen and GAG levels decreased in decellularized samples (about 20% of loss in 3D-dECMs \*\**p*<0.01; Fig. 4C).

The neoplastic 3D-dECMs showed a broader distribution of Young Modulus (YM) values and were markedly stiffer than the normal 3D-dECMs (Fig. 4D). Hovewer, the YM distributions were considerably scattered and a significant overlap was present between the two conditions. These results indicate that the ECM is a complex system that remains locally heterogeneous on the scale of several typical cellular lengths, i.e., 10-100 μm, because the transition from the normal to the neoplastic condition, in terms of change in stiffness and structural organization, is not uniform across the whole macroscopic tissue region.

Fig. 4E shows the results of the statistical analysis on median values of the YM measured in the different sites and conditions: the stiffening is statistically significant for four out of five patients. Note that the strongest relative stiffening in the tumor tissue, with an increase in the YM value of more than 200%, was measured in the tumor sample of the the youngest patient (Supplementary Fig. S3A-C). Overall, the YM and the relative increase in stiffness correlate with the age of the patients: the YM tends to increase with age, although its relative increase is smaller, suggesting a greater propensity for ECM remodeling in younger individuals.

### Decellularized scaffolds sustain PM-derived organoid growth

To develop an *in vitro* model of PM disease, 3D-dECMs were repopulated with C1, C2 and C3 TDO, with mutational profiles resembling the most common gene alterations in CRC (C1 was KRAS-mutant, C2 BRAF-mutant, while C3 had amplified FGFR1). In line with literature data, which show that colonization of decellularized matrices takes from eight to 12 days, TDO were grown on the matrices for five, 12 and 21 days [12, 13, 29]. Five days after seeding, C1 organoids were localized along the perimeter of the 3D-dECMs generated from normal peritoneum (Fig. 5A). A similar distribution was observed 12 and 21 days after seeding, with a slight increase in the number of cells in the stromal region. In 3D-dECMs generated from neoplastic peritoneum, instead, C1 TDO were distributed throughout the matrix five days after seeding, and colonization was evident with high stromal infiltration on day 21 (Fig. 5A and Supplementary Fig. S4A and S4B). After 21 days, TDO grown on neoplastic-derived 3D-dECMs were able to infiltrate and colonize the ECM much better than TDO grown on normal-derived 3D-dECMs (Fig 5B and Supplementary Fig. S4C and S4D). Cell density of C1, C2 and C3 TDO grown on neoplastic-derived 3D-dECM was 520, 960 and 980 cells/mm^2^, respectively, and decreased to 200, 470 and 500 cells/mm^2^ when TDO were cultured on normal-derived 3D-dECMs. Morphological characteristics and the efficiency of TDO infiltration were correlated with grade and differentiation state of their tumours of origin. Organoids derived from grade III/G2 metastatic lesions with moderate undifferentiation and low invasive capacity (C1), infiltrated to the 3D-dECM as single cells, while organoids from poorly undifferentiated grade IV/G3 tumors with BRAF mutation (C2) and FGFR1 amplification (C3), infiltrated into the 3D-dECMs maintaining the spheroid shape (Fig.5A and 5B, Supplementary Fig. S4A-D).

**Fig. 5.**
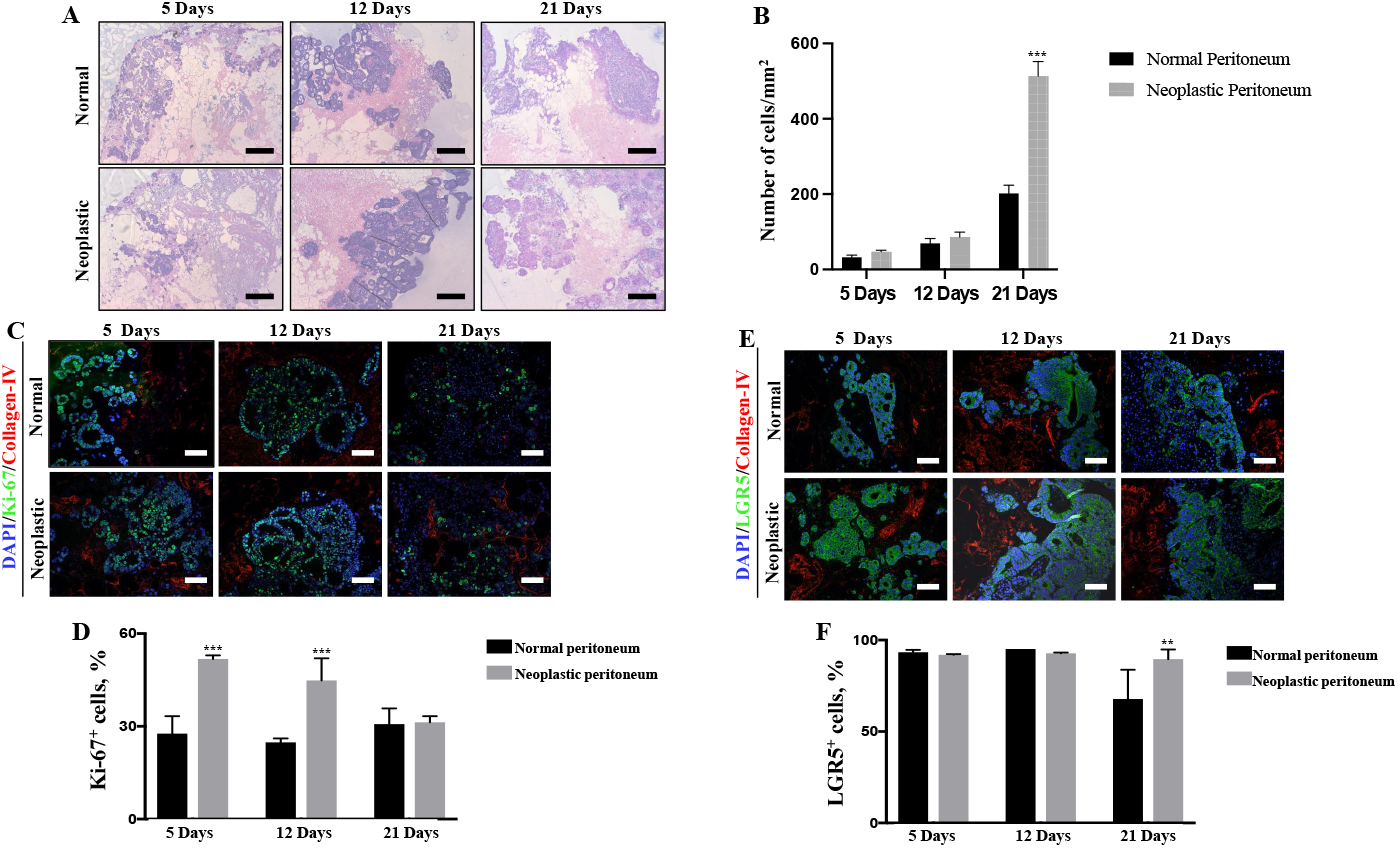
Peritoneum-derived 3D-dECM scaffolds support colonization, infiltration and proliferation of PM-derived organoids, maintaining the stem cell pool. **(A)** H&E staining of decellularized matrices derived from normal (top) or neoplastic (bottom) peritoneum repopulated with PM-derived organoids (C1) as indicated. Scale bar: 50 μm. The repopulation experiments were performed in triplicate. **(B)** Number of PM-organoid derived cells growth onto normal and neoplastic-derived 3D-dECM per mm^2^ ma after 5, 12 and 21 days. Data are presented as median and SD of three fields per experiments, counted using Qpath software. One-way ANOVA (\*\*\**p*<0.001). **(C)** IF analysis of 3D decellularized matrices derived from normal (top) and neoplastic (bottom) peritoneum repopulated with organoids (C1) using Ki-67^+^ (green) and collagen IV^+^ (red) antibodies as indicated. The samples were counterstained with DAPI (blue). Scale bar: 20 μm. **(D)** Proliferation rate of organoids measured as the percentage of Ki-67^+^ cells present in fields devoid of dead cells. Data are presented as median and SD of five fields per experiment (40X magnification), counted using Qpath software. One-way ANOVA (\*\*\**p*<0.001). **(E)** IF analysis of 3D decellularized matrices derived from normal (top) and neoplastic (bottom) peritoneum repopulated with organoids (C1) using LGR5^+^ (green) and collagen IV^+^ (red) antibodies as indicated. The samples were counterstained with DAPI (blue). Scale bar: 20 μm. **(F)** Amount of stem cells in organoids, measured as the percentage of LGR5^+^ cells present in fields devoid of dead cells. Five fields per experiment (40X magnification) were counted. Data are presented as median and SD of five fields per experiment (40X magnification), counted using Qpath software. One-way ANOVA (\*\**p*<0.01).

### 3D decellularized scaffolds support the proliferation of organoids

Results showed that 3D-dECMs generated from neoplastic peritoneum had a significantly higher percentage of Ki-67-positive cells five and 12 days (\*\*\**p*<0.001) after seeding, indicating that these organoids underwent a faster growth (Fig. 5C, D and Supplementary Fig. S4E-H). No differences were observed 21 days after seeding. After 12 days of growth, the fraction of Ki-67-positive cells was less that at five days; however, at both times the fraction of Ki-67-positive cells was greater on the neoplastic matrix than on the normal matrix, indicating that the neoplastic 3D-dECMs promote TDO proliferation.

LGR5-positive staining showed that the stem cell pool was maintained on both normal and neoplastic 3D-dECMs five and 12 days after seeding (Fig. 5E and 5F, Supplementary Fig. S4E-H). At day 21, the stem cell pool was significantly lower on 3D-dECMs generated from normal peritoneum compared with 3D-dECMs generated from neoplastic peritoneum. Most likely, after 21 days of growth, the cells are at confluence and the stem cells pool grown on the neoplastic matrix has an environment that favors the maintainance of its phenotype, as demonstrated by transcriptomics analyses (see below and supplementary Fig S6C).

### 3D-dECM stiffness does not activate YAP/TAZ proteins

Expression of the transcription factors Yes-associated protein (YAP) and transcriptional co-activator with PDZ binding motif (TAZ) was investigated on TDO grown on differente substrates. YAP and TAZ are sensors of the structural and mechanical characteristics of the cell microenvironment and can translate changes in ECM stiffening into genomic transcriptional alterations [18]. IHC analyses showed that YAP/TAZ were expressed in all TDO grown in Matrigel, with YAP mainly located in the nucleus and TAZ in the cytoplasm (Fig. 6A, Supplementary Fig S5A-B). YAP/TAZ positive cells were stable at all times (5, 12 and 21 days), ranging from 75 to 90% (Supplementary Fig. S6A-B). In contrast, YAP and TAZ were not expressed in TDO grown on normal and neoplastic-derived 3D-dECMs, indicating that the peritoneal-derived matrix is able to modulate and block their expression levels (Fig 6A, Supplementary Fig. S5A-B). Both proteins were expressed only in one (C1) of the three tumors from which TDO were derived (Fig. 6A, Supplementary Fig. S5A-B), with YAP mainly located in the nucleus and TAZ in the cytoplasm.

**Fig. 6.**
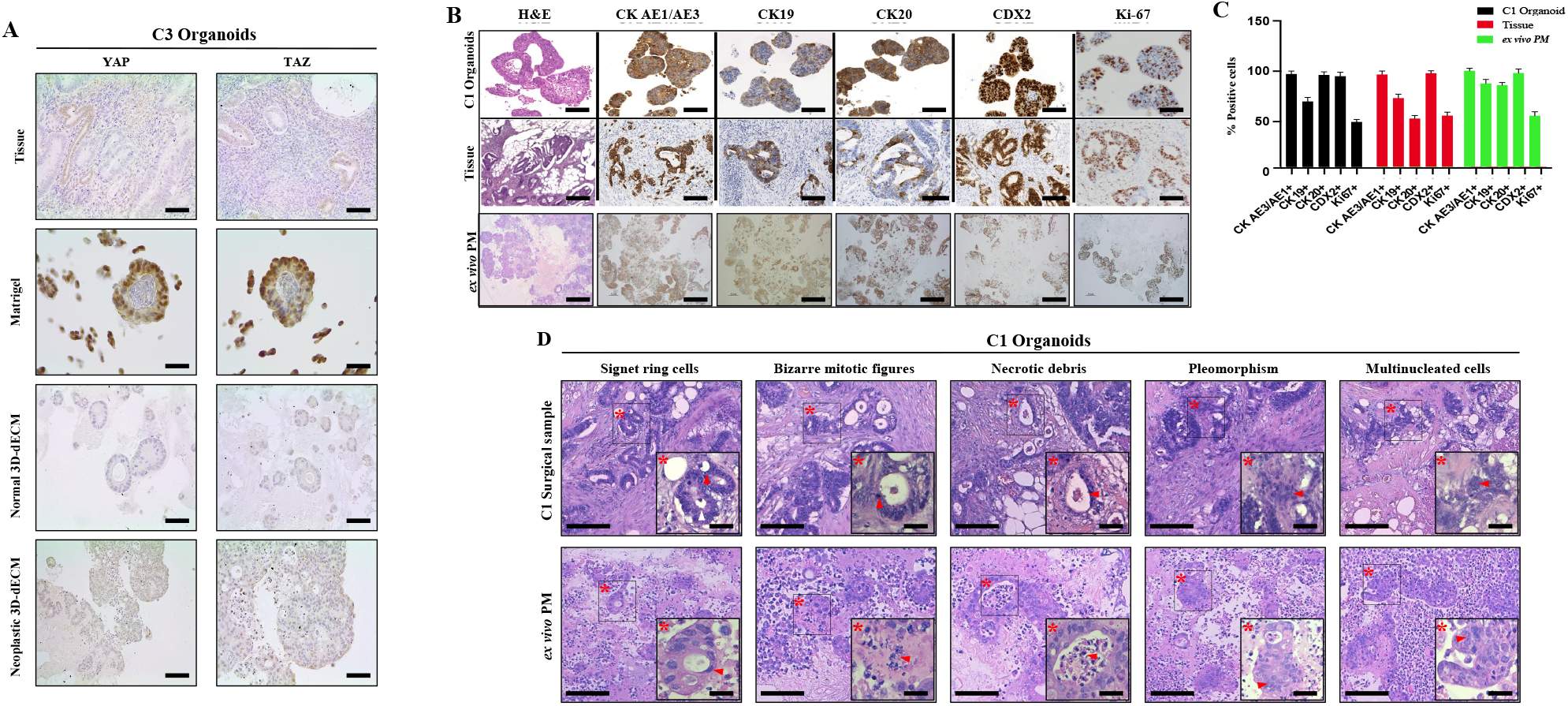
*ex vivo* engineered PM lesions are comparable to PMs found *in vivo*. **(A)** Comparative immunohistochemical images of PM-derived organoids (C3) grown on different substrates (Matrigel, Normal 3D-dECM and Neoplastic 3D-dECM) and their corresponding tumor of origin. Expression of YAP and TAZ proteins was analysed. Scale bar: 100 μM. **(B)** Comparative histological and immunohistochemical images of organoids (C1) versus their corresponding tumor of origin and the *ex vivo* engineered PM lesion. Samples were analyzed for the expression of the CRC-specific markers: CK AE1/AE3, CK19, CK20, CDX2, and Ki-67. Scale bar: 100 μM. Images in the first two lanes were previously published (20). **(C)** Quantitative counts of the percentage of CK AE/AE3, CK19, CK20, CDX2 and Ki-67 positive cells in C1 organoids Vs their corresponding tumor of origin and the *ex vivo* engineered PM lesion. Data are presented as median and SD of three fields per experiment, counted using Qpath software. One-way ANOVA did not show differences between the three groups. **(D)** Histological comparison of peritoneal metastasis and neoplastic-derived 3D-dECMs repopulated with PM-derived organoids (C1). The *ex vivo* engineered PM lesions present histological features that are typical of PMs of gastrointestinal origin. Asterisks and arrows indicated the main morphological features. Scale bar: 20 μm.

### *ex vivo* engineered PM lesions reproduce patient PM

IHC analysis of the TDO using colorectal markers showed that repopulated 3D-dECMs retain the main characteristics and the morphology of their tumor of origin (Fig. 6A and Supplementary Fig. S5A and S5C), expressing the colorectal markers in the same percentage of cells as their tissue of origin (Fig. 6B and Supplementary Fig S5E and S5F). The *ex vivo* PM lesions presented the typical histological features observed in PM patients, such as: i) signet ring cells, ii) bizarre mitotic figures, iii) necrotic debris, iv) pleomorphic cell size and shape, and v) multinucleated cells (Fig. 6C and Supplementary Fig. S5B and S5D). Signet ring cells have been reported in another *ex vivo* model system [33], highlighting that our model of PM is highly representative of *in vivo* lesions.

### Gene expression analysis of engineered PM lesions

RNA sequencing (RNA-seq) analysis of the TDO (Fig. 7) highlighted differences between 3D-dECMs and Matrigel substrates: 327 differentially expressed genes (DEGs) were identified in TDO grown on normal 3D-dECMs, and 144 DEGs were identified in TDO grown on neoplastic 3D-dECMs compared to TDO grown in Matrigel (|FC| > 1.5 and adj *p*-value <0.05).

**Fig. 7.**
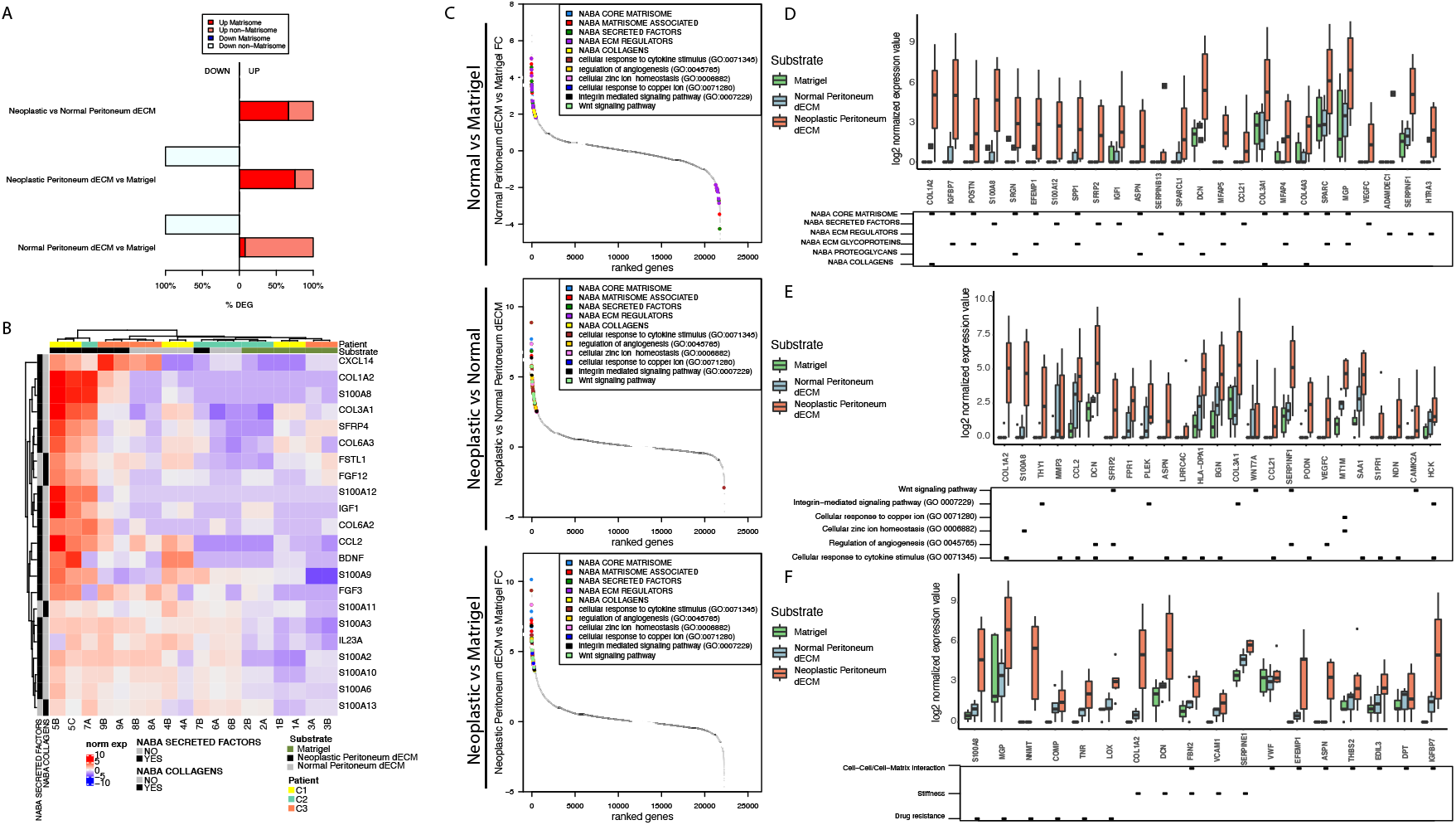
Gene expression analysis of engineered PM lesions. **(A)** Percentage of up- and down-regulated genes belonging to the Matrisome dataset, in organoids grown on 3D-dECMs compared to Matrigel or in neoplastic versus normal 3D-dECM. **(B)** Unsupervised hierarchical clustering of the organoids according to the expression of the top DEGs included in Naba Secreted Factors and in Naba Collagens categories. **(C)** Fold changes of genes belonging to the indicated gene sets among the top 100 deregulated genes. Gene ranks for relative fold change are shown on the x-axis and the logFCs on the y-axis. **(D)** Box plot showing the expression of genes selected for their involvement in the indicated processes of the Naba Matrisome datasets. Median and interquartile range are displayed as horizontal lines. Black squares in the bottom panel indicate which category the genes belong to. **(E)** Expression of genes selected for their involvement in the indicated processes of GO BP and KEGG databases. Median and interquartile range are displayed as horizontal lines. Black squares in the bottom panel indicate which category the genes belong to. **(F)** Expression of genes selected for their involvement in the indicated processes, using a selection of genes related to the following biological processes: cell-cell/cell-matrix interactions, extracellular matrix stiffness and drug resistance. Median and interquartile range are displayed as horizontal lines. Black squares in the bottom panel indicate which category the genes belong to.

The most represented biological processes include cell-cell/cell-matrix interactions, organoid behavior and interactions with the ECM, angiogenesis, metal ion homeostasis, and response to external stimuli. 3D-dECMs deregulated many genes fundamental for the 3D-architecture/organization of the ECM. Many of the up-regulated genes identified in TDO grown on neoplastic 3D-dECM were assigned to the Matrisome database (67% for neoplastic versus normal 3D-dECM, and 25% for neoplastic 3D-dECM versus Matrigel). Only 8% of the up-regulated genes identified in TDO grown on normal 3D-dECM versus TDO grown in Matrigel belonged to the Matrisome database (Fig. 7A). Unsupervised hierarchical clustering of TDO highlighted a separation between TDO grown on neoplastic 3D-dECM and TDO grown on normal 3D-dECM and in Matrigel substrates (Fig. 7B and Supplementary Fig. S6A-C). TDO grown on both normal and neoplastic 3D-dECMs presented an over-representation of DEGs involved in ECM composition, regulation and modulation (Fig. 7C). Growth on 3D-dECM also favors the expression of genes involved in the regulation of angiogenic processes, the response to cytokine stimuli, the integrin pathway, as well as copper and zinc metabolism. The same categories were found for both normal and neoplastic-derived 3D-dECMs (Supplementary Fig. S6C and S6D).

Using the Matrisome database, we identified genes related to the core composition of the ECM and to of ECM interactors/regulators through secretion of specific factors, all of which were higher in 3D-dECMs derived from neoplastic tissue (Fig. 7D and Supplementary Fig. S6E-H). 3D-dECMs presented high expression of genes involved in stem cell pathways, the cellular response to cytokines, zinc and copper metabolism, the integrin pathway, and the regulation of angiogenesis (Fig. 7E and Supplementary Fig. S6E-H). Similar results were found for genes involved in cell-cell/cell-matrix interactions (Fig. 7F). The differences observed with transcriptomic data were validated by qRT-PCR of some representative genes on C1, C2 and C3 TDO grown in Matrigel and on normal and neoplastic 3D-dECMs. The tissue of origin of the six TDO was also analyzed. The trends observed by RNA-seq were confirmed and all the genes analysed were also expressed in the tissue of origin (Supplementary Figure S6I and S6L).

### 3D-dECMs decrease the efficacy of HIPEC treatments

The TDO analysed had different IC_50_ values: C3 TDO were the most sensitive and C2 the least sensitive to both MMC and OXA (Fig. 8A, Supplementary Fig. S7C and S7E). At these values MMC treatment induced DNA damage and apoptosis in all TDO, as shown by phosphorylation of P53 (Ser 15) and H2AX (Ser139) (Fig. 8B and Supplementary Fig. S7D) and cleavage of PARP and CASPASE3(Fig. 8b and Supplementary Fig. S7D). Treatment with OXA also induced DNA damage and apoptosis in all TDO, although C2 TDO only showed PARP cleavage (Supplementary Fig. S7G).

**Fig. 8.**
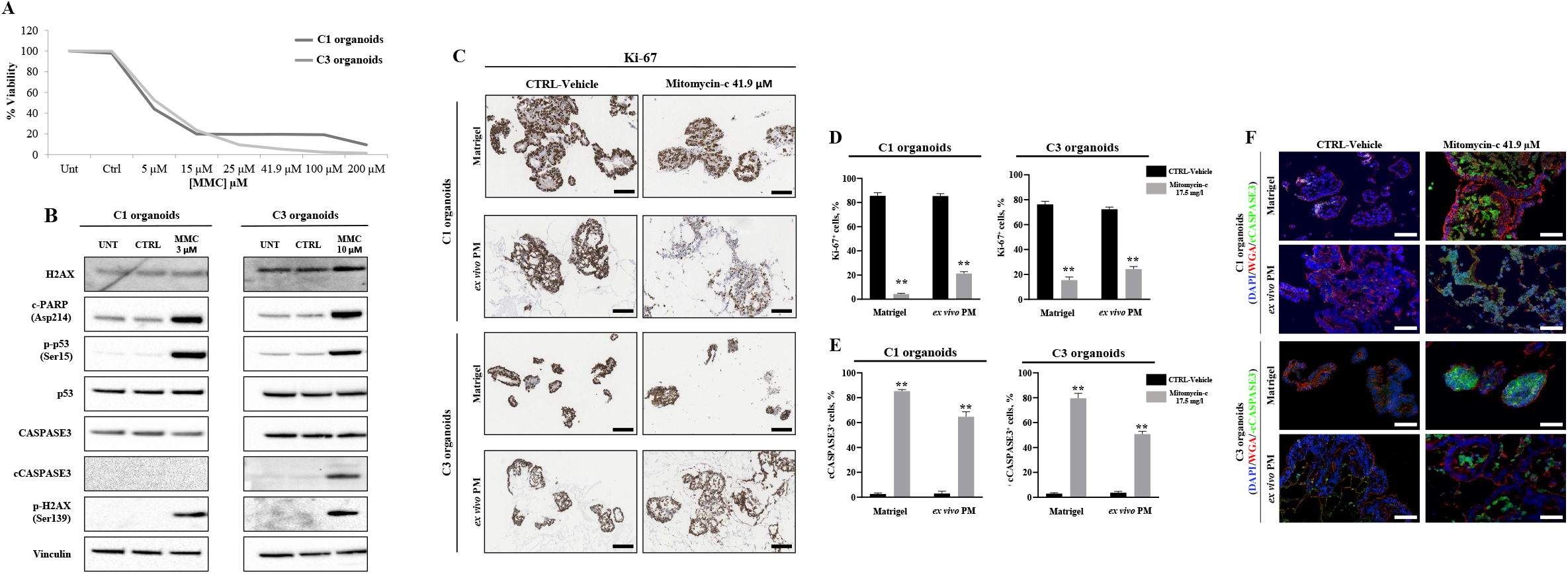
3D-dECM scaffolds decrease the efficacy of HIPEC treatments. **(A)** Dose-response curve of C1 and C3 organoids cultured in Matrigel and treated with MMC at different concentrations at 42.5 °C for 1 h. **(B)** Immunoblots of cPARP, p-p53, p53, CASPASE3, cCASPASE3, p-H2AX and H2AX in C1 and C3 organoids treated with MMC 5 and 10 μM respectively. Vinculin was used as loading control. **(C)** IHC analysis of C1 and C3 organoids cultured in Matrigel and on neoplastic-derived peritoneal 3D-dECMs, after *in vitro* HIPEC treatments, using Ki-67 immunostaining. Scale bar: 50 μm. **(D)** IF analysis of C1 and C3 organoids cultured in Matrigel and on neoplastic-derived peritoneal 3D-dECMs, after HIPEC treatments, using cCASPASE-3 antibody (green). The samples were counterstained with WGA (red) and DAPI (blue). Scale bar: 20 μm. **(E)** Proliferation rate of PM-derived organoids (C1, left panel; C3, right panel) measured as the percentage of Ki-67^+^ cells present in fields devoid of dead cells. Data are presented as median and SD of five fields per experiment (40X magnification), counted using Qpath software. One-way ANOVA (\*\**p*<0.01). **(F)** Percentage of apoptotic organoids (C1, left panel; C3, right panel) measured as the percentage of cCASPASE-3^+^ cells present in selected fields. Data are presented as median and SD of five fields per experiment (40X magnification), counted using Qpath software. One-way ANOVA (\*\**p*<0.01).

To evaluate the contribution of the ECM to treatment response, we simulated treatment with HIPEC on ex vivo engineered micrometastases. C1, C2 and C3 TDO were grown on neoplastic 3D-dECMs for 12 days, time considered sufficient for the colonization of the matrix surface [29], and treated with MMC and OXA. MMC treatment induced cellular disruption in C1 and C3 organoids (Supplementary Fig. S7A). Ki-67 expression was lower in the treated group, also showing a diffuse cytoplasmatic signal (Fig. 8C and Supplementary Fig. S7B). The number of Ki-67-positive C1 TDO grown in Matrigel or on 3D-dECMs was reduced by 95% and 72%, respectively, compared to the untreated control groups (\*\**p*<0.01; Fig. 8D). Proliferation was reduced by 80% and 66%, respectively, in C3 organoids grown in Matrigel or on 3D-dECM compared to the untreated control groups (\*\**p*<0.01; Fig. 8D). Immunofluorescence (IF) staining with cleaved CASPASE3 (cCASPASE3) confirmed that untreated C1 and C3 TDO were alive (Fig. 8D, left panel). The number of cCASPASE3-positive C1 TDO grown in Matrigel or on 3D-dECMs was 80 % and 60 % respectively, compared to the untreated control groups (**p<0.01; Fig. 8D), and 80 % and 50 % respectively in C3 TDO (**p<0.01; Fig. 8F). MMC treatment had no effect on C2 organoids (Supplementary Fig. S7C-E).

Regarding OXA treatment, the number of CASPASE3-positive TDO grown in Matrigel or on 3D-dECMs was respectively 50 % and 70 % for C1, 60 % and 85 % for C2 and 45 % and 65 % for C3, compared with untreated control groups (** p < 0.01; Supplementary Fig. S7F-H).

These results demonstrate that HIPEC induces apoptosis in TDO and that presence of 3D-dECM hinders its efficacy (Fig. 8D and F).

## Discussion

Here we describe an engineered model that combines key features of TDO within the microenvironment, enabling the recapitulation of the PM niche under physiological conditions.

Transcriptome analysis showed that organoids grown on peritoneal matrices express genes involved in pathways that favor implantation into the matrix and remodelling of the ECM. The 3D-dECM models reproduce the complex tissue architecture and cell-matrix interactions of the native environment of PM. 3D-dECMs derived from the neoplastic peritoneum allow the development of a tissue microenvironment that preserves the stem cell pool of TDO, enhancing their proliferation and favoring a repopulation pattern similar to the one that is usually observed *in vivo*. Our findings are in line with previous works showing that cancer-derived ECM sustains the proliferation and invasion of CRC cell lines [34].

Scaffolds deriving from neoplastic peritoneum showed greater stiffness than those derived from normal peritoneum. Increased stiffness and crosslinking of the perilesional ECM was identified as an environmental change predisposing to CRC invasion [21]. The observed stiffening of the neoplastic ECM can be partially attributed to the more compact fine structure of the matrix and to the linearization of the fibers in bundles, as observed in Nebuloni et al. [21]. The increase in the amount of GAG in the perilesional ECM may also be related to the increase in stiffness. Some authors have highlighted the relationship between GAG and matrix stiffness: negatively charged GAGs provide a repulsive force that opposes compression and shear in the ECM [35]; moreover, GAGs ability to retain high quantities of water and hydrated cations confers resistance to compressive forces [35]. The stiffness of healthy tissues increases with the age of the patient [36]. The relative stiffening in the neoplastic tissues appears to be related to the aggressiveness of the tumor, which, in our study, was greater in the younger patient, confirming that tumor stiffness favors its metastatic spread [37]. The high amount of GAGs observed in neoplastic tissues could be the result of metastatic transformation. In fact, chondoritin-sulfate is the main binding site for the isoform v of CD44, which is a key player in the metastatic dissemination [38]. Transcriptome analysis also confirmed higher expression of GAGs in the repopulated neoplastic 3D-dEMCs. Further experiments will be performed with more donors to confirm the observed trends.

Increased stiffness of the dECM does not activate YAP/TAZ signaling, since TDO grown on 3D-dECMs did not express YAP/TAZ proteins. Both proteins are expressed in TDO grown in Matrigel, and in the tumor tissue from which C1 TDO were derived, but their localization (TAZ cytoplasmatic and YAP nuclear) does not allow their activity. In agreement with these results, pathways regulated by YAP/TAZ signaling were not found activated in the transcriptomic analysis. Our observations support the proposed role of YAP as a tumor suppressor in metastatic CRC, where it can inhibit the Wnt pathway by reprogramming *LGR5*-positive cells and, *in vivo*, by reducing Wnt activity [39]. Indeed, Wnt pathway activity was one of the most represented GO categories higher in TDO grown on 3D-dECMs than in Matrigel. Analysis of YAP/TAZ expression in tumor tissue from which TDO were derived corroborates this hypothesis, since YAP/TAZ were absent in patients with a more aggressive and indifferentiated tumor phenotype (C2 and C3). The presence of YAP/TAZ in TDO grown in Matrigel is probably due to the different stiffness of Matrigel compared to 3D-dECM and to the absence of factors previously released by the same tumor cells, which are instead present in the 3D-dECM, and which could regulate YAP/TAZ signaling activity. This result highlights how commercially available substrates fail to fully recapitulate the *in vivo* context. Factors other than YAP/TAZ signaling, present in the stromal compartment, could influence the mechanobiology of the peritoneum. For example, cancer-activated fibroblasts (CAFs) are influenced by the tumor cells and can activate a complex signaling network capable of remodeling and/or removing the ECM, through the activation of the TGFβ pathway [40]. The interactions between the ECM and stromal and tumor cells could also regulate the secretion of specific molecules that can induce fibrosis via cytokine and growth factors-mediated pathways [41]. Indeed, the cytokine signaling-mediated pathway was one of the most represented GO categories observed in the transcriptomic analysis. These data indicate the development of a complex cell-cell and cell-ECM communication network and suggest a prominent role of the stromal compartment on the mechanical characteristics of the peritoneal ECM. Future experiments will be needed to elucidate how stromal cells can affect mechanobiology of the peritoneal ECM, through co-repopulation experiments between tumor cells, ECM and stromal cells.

Expression of genes critical for tissue architecture and stiffness, ECM remodelling, fibril generation, epithelial cell differentiation, resistance to compression and regulation of angiogenesis [42–46] was higher in 3D-dECMs generated from neoplastic tissue than in 3D-dECMs obtained from normal tissue or Matrigel, confirming the ability of our model to reproduce the PM microenvironment.

Morphological and topographical experiments showed that the neoplastic scaffold undergoes complex structural modifications that enhance TDO adhesion and proliferation. In support of this concept, both normal and tumor 3D-dECMs showed an upregulation of genes involved in zinc and copper metabolism and homeostasis, which was higher in the neoplastic-derived peritoneal matrix. These metal ions are involved in the regulation and activation of the metalloproteinase enzymes, which are the main modulators of the ECM through a proteolytic activity [47] and could play a role in the activation of ECM remodeling pathways underlying the metastatic niche. All these observations make our model more attractive than the conventional culture methods with collagen-based scaffolds, which do not mimic real tissue conditions.

Transcriptomic analyses showed that repopulated 3D-dECMs presented features typical of the PM disease [48, 49] and expressed genes involved in ECM remodelling, such as NABA Matrisome and stem cell-related genes, ECM regulators and genes involved in the response to cytokine and pro-inflammatory stimuli, integrin interactions and collagen/proteoglycans modifications. These pathways were less represented in normal 3D-dECMs and absent in Matrigel samples, confirming a previous study showing that growth on the ECM of normal colon organoids transfected with mutant *APC* induces features typically associated with CRC progression [12]. The deregulation of genes belonging to pathways involved in the metastatic process, linked to metastatic spread and the development of the metastatic niche [5, 50], is in agreement with the fact that our organoids derive from metastatic lesions, where the cells activate a series of pathways to better adapt to the niche. HIPEC simulation experiments highlighted the potential role of the neoplastic ECM in the development of drug resistance. Transcriptomic data indicated the activation of mechanisms correlated with drug resistance along with the modification of the mechanical properties of the ECM. In fact, growth on scaffolds increased the expression of anti-apoptotic and pro-survival genes, as well as genes involved in resistance to platin-based drugs. High expression of stiffness-related genes was also observed. Also integrins, which are involved in ECM remodeling and can function as mechanotransducers, contributing to cancer metastasis, stemness and drug resistance [51] were one of the most representative GO categories in the transcriptome analyses, supporting the activation of drug resistance mechanism when TDO are grown on 3D-dECMs. Dose–response curves to MMC and OXA showed that the TDO had different drug sensitivity and that the doses of MMC and OXA administered clinically were insufficient to eliminate all cancer cells. Response to treatment was even less for TDO grown on 3D-dECMs, which showed greater resistance to both drugs. In support of these findings, genes involved in drug resistance, especially to platinum-based compounds, were upregulated in TDO grown on 3D-dECMs. The results observed with the model better reproduce the results in clinics, as approximately 60% of patients treated with HIPEC recur in one year. All these findings highlight that the engineered model we propose could be a drug screening tool that more faithfully recapitulates the tumor microenvironment and response to treatment for tailored therapies than the classical monoculture 2D models, or even 3D-cultures, which are still being used [52].

However, the model presents some limitations, as it still does not reach a resolution level that allows it to mimic the PM niche in all its constituents. In fact, other components of the microenvironment play a fundamental role in the spread of PM [53, 54], such as immune surveillance and the vascular system [55, 56]. Further optimization of our model will, therefore, imply the reconstruction of a specialized physiological microenvironment by incorporating vascular networks, the immune system, as well as organ-specific microbes. Moreover, the replacement of patient-derived 3D-dECM with a synthetic support with the same biochemical and physical characteristics of the components of the decellularized matrix will improve the reproducibility and allow personalized drug screenings to be performed on TDO.

Our 3D model could be used as a pre-clinical platform to study the role of tumor ECM in the development of the PM niche. The model represents a physiological tool that can aid the identification of key players in the metastatic development, and may allow the selection, in a biologically relevant setting, of new therapeutic strategies, also providing a new tool to boost the bench-to-bed-side process to improve patient care. Finally, the approach described here might be used to generate other types of *ex vivo* metastatic niches.

## Acknowledgements

The authors wish to thank the Imaging Technological Development Unit (TDU) of the FIRC Institute of Molecular Oncology, IFOM, Milan, Italy and the DNA Sequencing Facility, in particular Sara Volorio, Mirko Ribone and Claudia Valli, of Cogentech Ltd, Benefit Corporation with a Sole Shareholder, Milan, Italy. We also wish to thank Oscar Illescas Pomposo from Fondazione IRCCS Istituto Nazionale dei Tumori for helping us in TDO mainteinance and development. The authors express their gratitude to Wessen Maruwge, an English author’s editor in Milan, for revising the manuscript.

## Author contributions

L.V., S.B. designed and performed all the experiments and edited the manuscript. L.V., M.G1., M. G2 and M.A.P., conceived and design the study. M.C., E.L., A.O., and A.P., performed AFM analysis and edited the manuscript. A.O., E.C., and D. P., aided all the imaging acquisition. F. Z., and F. I., performed all the bioinformatics analyses and critically discussed the data. S.Z., S. P. M., and S. F., performed RNA-seq and NGS experiments. M.G1., K.S., D.B., and M.D., performed surgical intervention and collected human samples. M.F., and G.G., performed IHC and HC analyses. L.C., and M.M., selected the cases and interpreted IHC and HC analyses. A.B., aided in generating organoids. M.V., aided in the transcirptomics data interpretation. L.V., M.G1., M.G2., A.P., and M.A.P., wrote the manuscript. L.V., S.Z. M.G2., and M.A.P., critically read the manuscript

