## supplementary material for "Decellularized Normal and Tumor Extracellular Matrix as Scaffold for Cancer Organoid Cultures of Colorectal Peritoneal Metastases"

### Supplementary Table S1

| Factor | Description | Vendor | Working Concentration |
|---|---|---|---|
| Gentamicin | Antibiotic | ThermoFisher Scientific | 50 ng/ml |
| HEPES | Buffer | ThermoFisher Scientific | 10 mM |
| L-Glutamine (GlutaMAX) | Cell culture supplement | ThermoFisher Scientific | 2 mM |
| B27 | Cell culture supplement | ThermoFisher Scientific | 1:50 |
| Gastrin-1, recombinant human | Recombinant protein | Sigma Aldrich | 10 nM |
| N-acetylcysteine | Colonic niche factor | Wako | 1 mM |
| EGF, recombinant human | Recombinant protein | ThermoFisher Scientific | 50 ng/ml |
| Noggin, recombinant human | Recombinant protein | Preprotech | 100 ng/ml |
| R-spondin-1, recombinant human | Recombinant protein | Preprotech | 100 ng/ml |
| Wnt3A, recombinant human | Recombinant protein | Preprotech | 50 ng/ml |
| Prostaglandin E2 | Colonic niche factor | Tocris | 100 nM |
| A83-01 | p38 inhibitor | Tocris | 500 nM |
| SB202190 | ROCK inhibitor | Sigma Aldrich | 10 µM |

**Supplementary Table S1:** The complete list of growth factors, media supplements and concentrations used for PM-derived organoid cultures.

### Supplementary Table S2

| Organoid culture | Medium composition |
|---|---|
| **C1** | DMEM-F12; B27; Glutamax; N-acetylcysteine; prostaglandin-E2; gastrin-I |
| **C2** | DMEM-F12; B27; Glutamax |
| **C3** | DMEM-F12; B27; Glutamax; N-acetylcysteine; prostaglandin-E2; gastrin-I; A83-01 |
| **C4** | DMEM-F12; B27; Glutamax; N-acetylcysteine; prostaglandin-E2; gastrin-I; EGF; A83-01; Noggin |
| **C5** | DMEM-F12; B27; Glutamax; N-acetylcysteine; prostaglandin-E2; gastrin-I; A83-01; SB202190; Noggin |
| **C6** | DMEM-F12; B27; Glutamax; N-acetylcysteine; prostaglandin-E2; gastrin-I; A83-01; Noggin |

**Supplementary Table S2**: The specific media composition for each PM-derived organoid culture.

### Supplementary Table S3

| Antigen (human) | Host | Clone | Vendor | Dilution | Antigen retrieval solution |
|---|---|---|---|---|---|
| **Ki-67** | Mouse | MIB-1 | Dako | 1:400 | 5 mM EDTA (pH 8), 10 min, 96 °C |
| **CK19** | Mouse | A53-B/A2 | Dako | 1:1000 | 5 mM EDTA (pH 8), 15 min, 96°C |
| **CK20** | Mouse | Ks20.8 | Dako | 1:500 | 5 mM EDTA (pH 8), 30 min, 96°C |
| **CK AE1/AE3** | Mouse | AE1+AE3 | Dako | 1:100 | 10 mM Citrate (pH 6), 15 min, 96°C |
| **CDX2** | Mouse | CDX2_88 | Dako | 1:50 | 5 mM EDTA (pH 8), 30 min, 96°C |
| **LGR5** | Mouse | OTI2A2 | Origene | 1:250 | 10 mM Citrate (pH 6), 15 min, 96 °C |
| **Vimentin** | Mouse | V9 | Dako | 1:400 | 10 mM Citrate (pH 6), 15 min, 96°C |
| **Collagen-IV** | Rabbit | CIV-22 | Abcam | 1:200 | 5 mM EDTA (pH 8), 10 min, 96 °C |
| **cCASPASE3** | Rabbit | #9661 | Cell Signaling | 1:250 | 10 mM Citrate (pH 6), 15 min, 96 °C |
| **TAZ** | Rabbit | WWTR1 | Sigma | 1:100 | 10 mM Citrate (pH 6), 15 min, 96 °C |
| **YAP** | Mouse | 63.7 | Santa Cruz | 1:100 | 10 mM Citrate (pH 6), 15 min, 96 °C |

**Supplementary Table S3:** The primary antibodies and the experimental conditions used for IHC and IF analyses.

### Supplementary Table S4

| Antigen (human) | Vendor | Host | Code | MW | Dilution |
|---|---|---|---|---|---|
| **Vinculin** | Sigma | Mouse | V9131 | 116 kDa | 1:10000 in 5% MILK |
| **Cleaved PARP (Asp214)** | Cell Signaling | Rabbit | 9541S | 89 kDa | 1:1000 in 5% MILK |
| **p-p53 (Ser15)** | Cell Signaling | Rabbit | 9284S | 53 kDa | 1:1000 in 5% MILK |
| **p53** | Santa Cruz | Mouse | sc-126 | 53 kDa | 1:1000 in 5% MILK |
| **Caspase-3** | Cell Signaling | Rabbit | 9662 | 35, 19, 17 kDa | 1:1000 in 5% BSA |
| **Cleaved Caspase-3 (Asp175)** | Cell Sigaling | Rabbit | 9661S/L | 17, 19 kDa | 1:1000 in 5% MILK |
| **p-Histone H3(Ser10 Thr11)** | Abcam | Rabbit | Ab 32107 | 17 kDa | 1:2500 in 5% BSA |
| **p-H2AX (Ser139)** | Upstate, Millipore | Mouse | 05-636 | 17 kDa | 1:1000 in 5% MILK |
| **H2AX** | Abcam | Rabbit | Ab11175 | 15 kDa | 1:5000 in 5% MILK |

**Supplementary Table S4:** The primary antibodies and the experimental conditions used for WB analyses.

### Supplementary Table S5

| ID | Diagnosis | Grade | Mutations | Microsatellite | Chemotherapy | PM Organoid line |
|---|---|---|---|---|---|---|
| S07-7576 | Moderately differentiated infiltrating adenocarcinoma | pT3G2 | G12S KRAS;TP53 | MSS | None | C1 |
| S11-2361 | Poorly differentiated mucinous multiple intestinal adenocarcinoma | pT4G3 | V600E BRAF; TP53 | MSS | None | C2 |
| S16-8598 | Intestinal mucinous adenocarcinoma | pT4G3N2 | G12S KRAS; TP53; FGFR1 amplification | MSS | Yes | C3 |
| S17-3963 | Moderately differentiated adenocarcinoma | T4aG3N2aM1b | G12S KRAS | MSS | None | C4 |
| S17-3610 | Intestinal adenocarcinoma with mucinous component | pT4aG2N0Mx | G12S KRAS | MSS | None | C5 |
| S18-8607 | Colloidal / gelatinous mucinous adenocarcinoma associated with hairy adenocarcinoma of high and low grade | T4aN2bG3Mx | G12S KRAS; TP53 | MSS | Yes | C6 |

**Supplementary Table S5:** The main pathological characteristics of the patients from whom the tumor specimens were obtained.

### Supplementary Figure S1

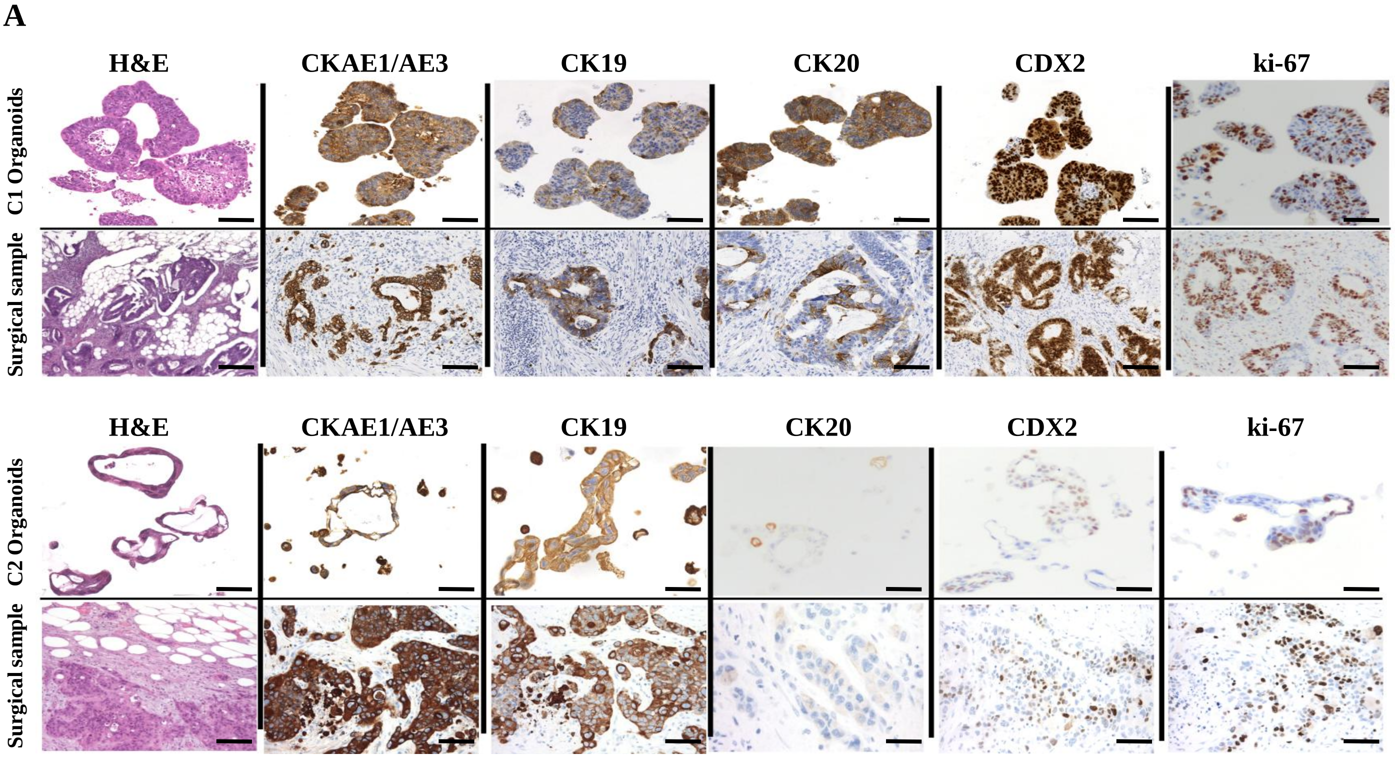


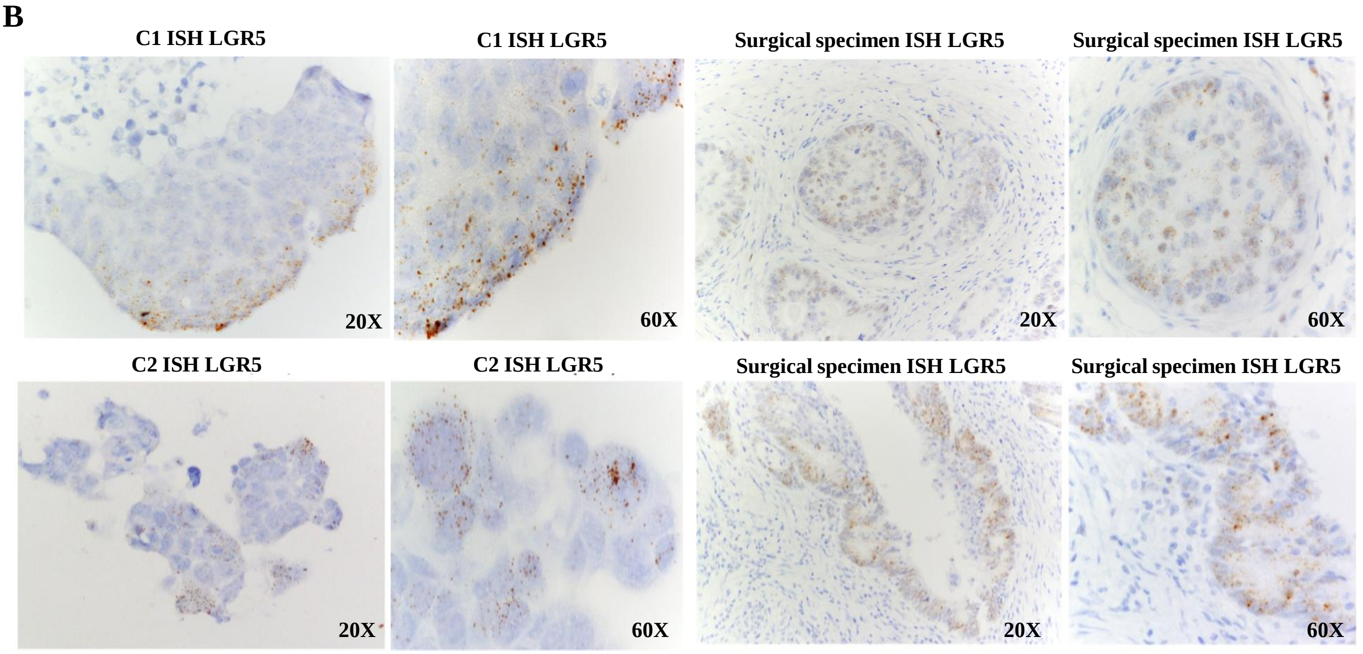


**Supplementary Fig. S1. (A)** IHC analysis of C1 and C2 organoids and their corresponding surgical samples, using H&E staining and CK AE1/AE3, CK19, CK20, CDX2 and Ki-67 immunostaining (20X magnification). The images were previously published by Bozzi et al. [11]. **(B)** In situ hybridization (ISH) of C1 and C2 organoids and corresponding surgical samples, using LGR5 immunostaining (20X magnification). The images were previously published [11].

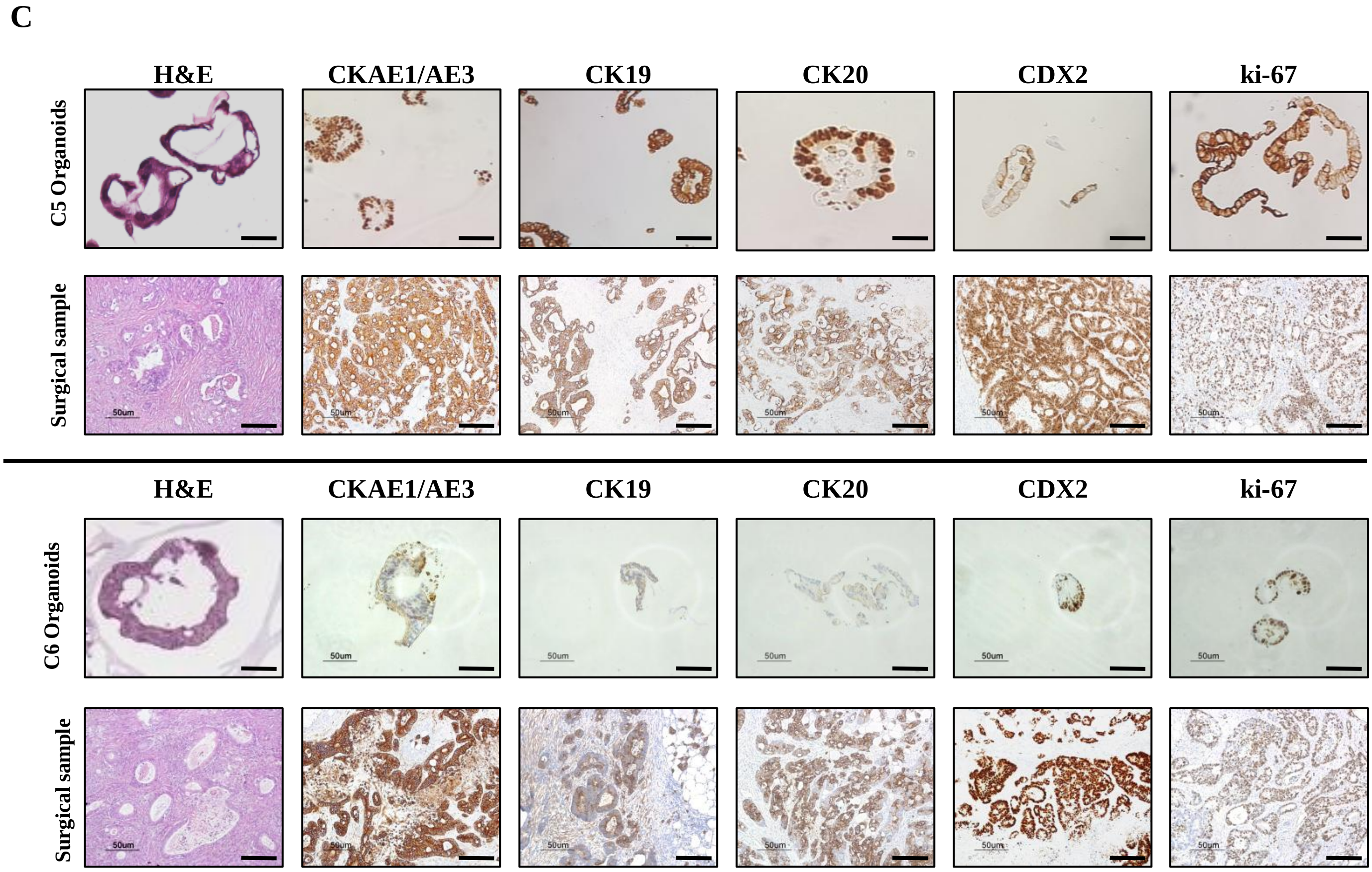


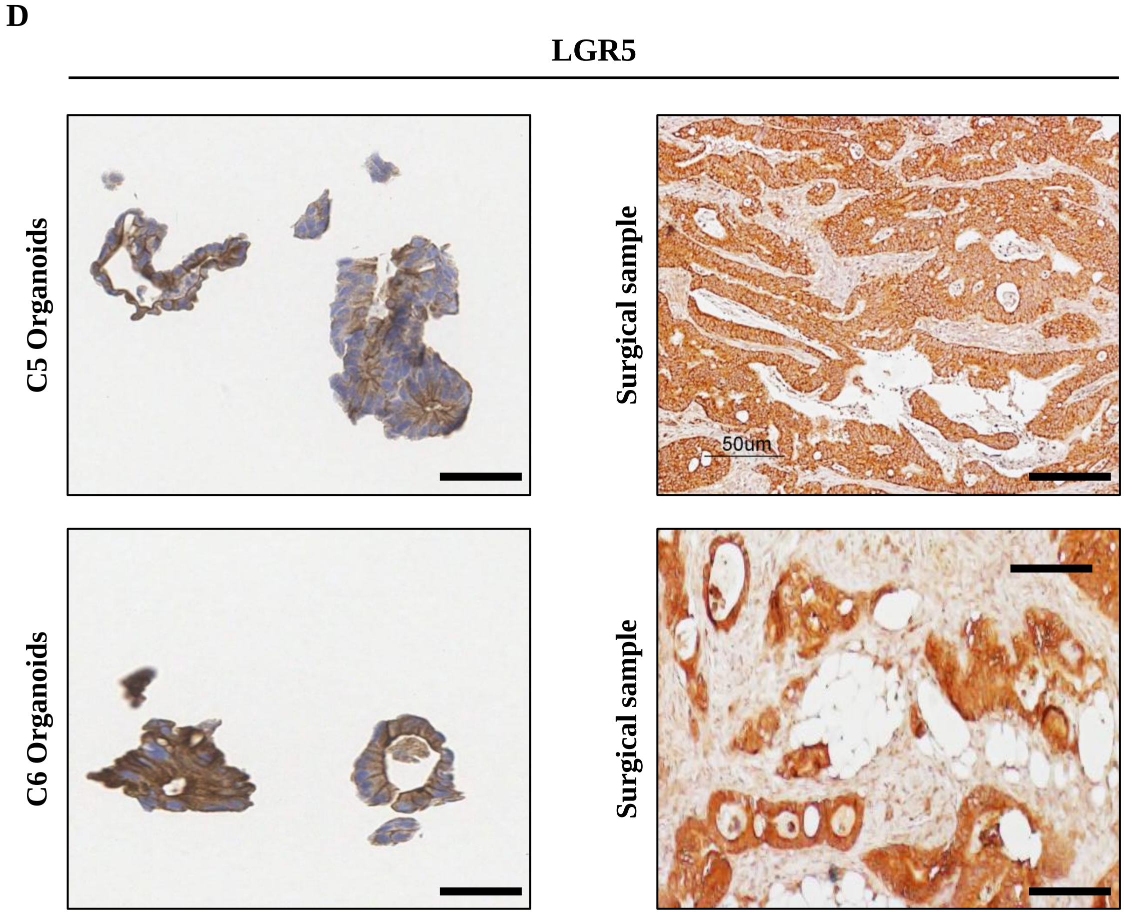


**Supplementary Fig. S1. (C)** H&E staining and CK AE1/AE3, CK19, CK20, CDX2, and Ki-67 immunostaining of C5 and C6 organoid cultures and their corresponding surgical samples. Scale bar: 50 µm. **(D)** IHC analysis of C5 and C6 organoid cultures and their tumor of origin, using LGR5 immunostaining. Scale bar: 50 µm.

**E**

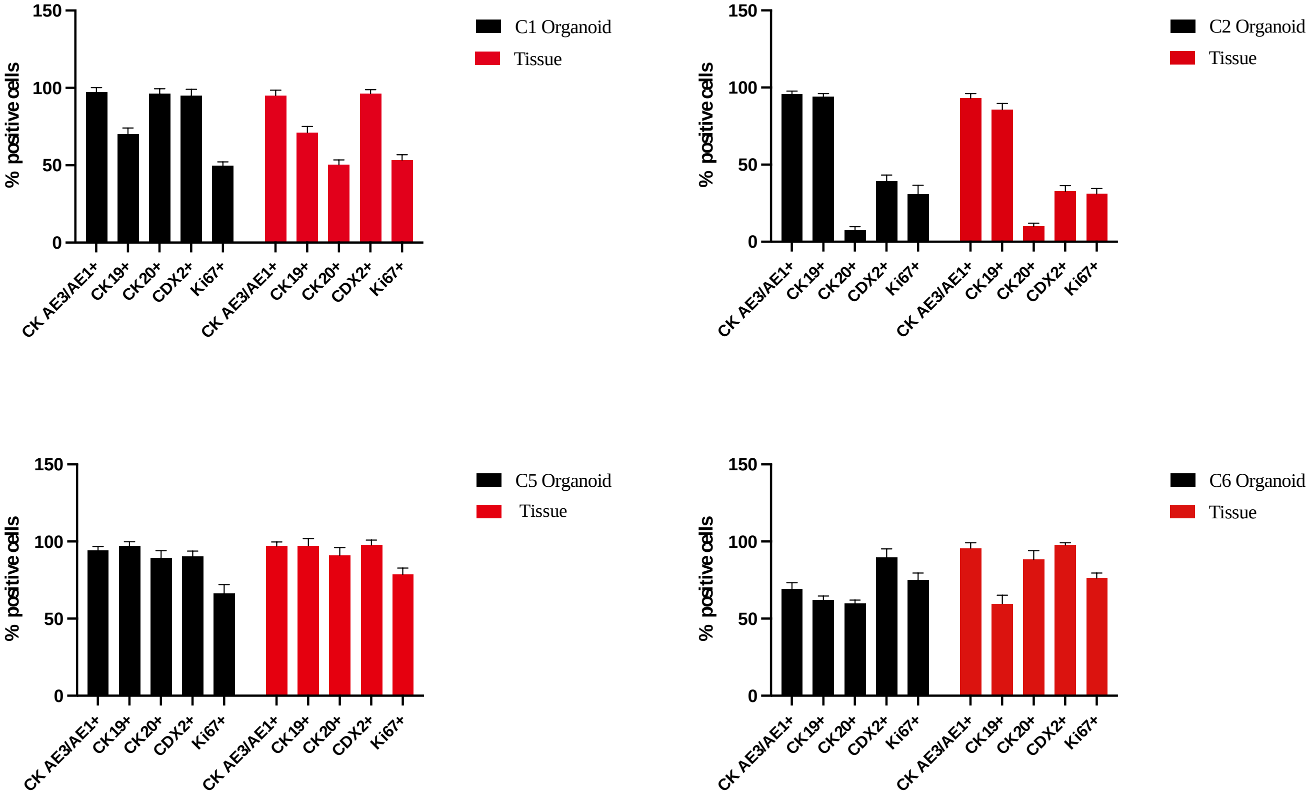


**F**

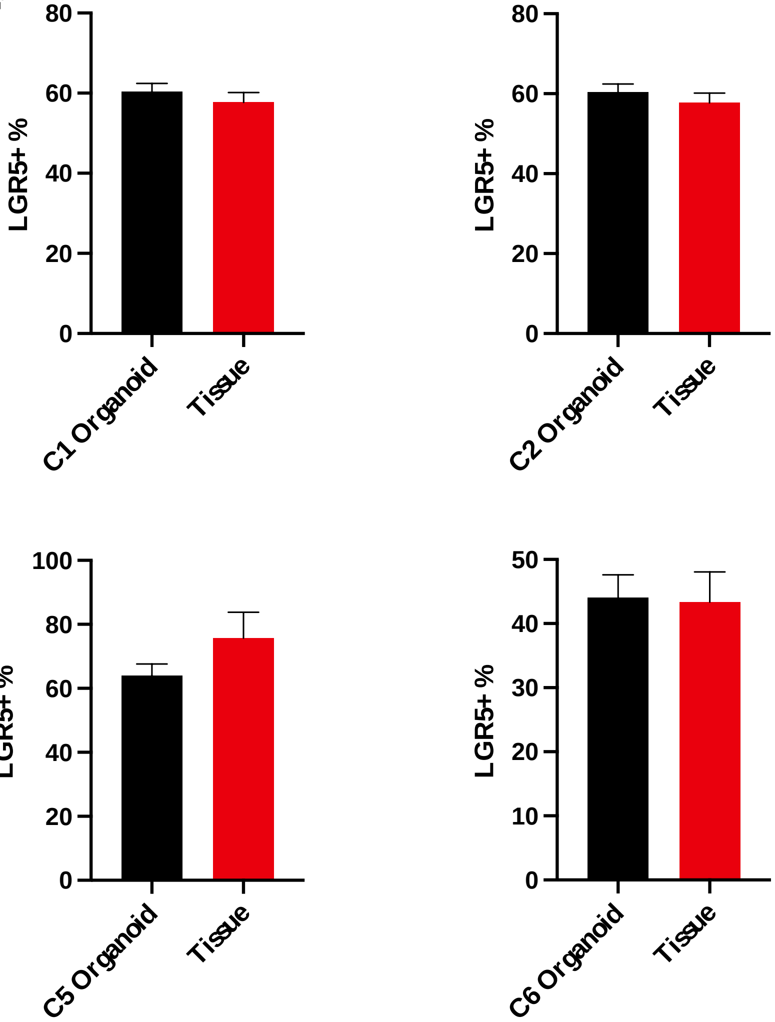


**Supplementary Fig. S1. (E)** Quantitative counts of the percentage of CK AE/AE3, CK19, CK20, CDX2 and Ki-67 positive cells in C1, C2, C5 and C6 PM-derived organoids Vs their corresponding tumor of origin. Three fields per experiments were counted using Qpath software. Data are presented as median and SD. One-way ANOVA did not show differences between the two groups. **(F)** Quantitative counts of the percentage of LGR5 positive cells in C1, C2, C5 and C6 PM-derived organoids Vs their corresponding tumor of origin. Three fields per experiments were counted using Qpath software. Data are presented as median and SD. One-way ANOVA did not show differences between the two groups.

**G**

| Genes | C1 Organoid | Tissue of origin |
|---|---|---|
| MSH3 (A62P) | | |
| BRCA2 (G267Q) | | |
| BRCA2 (S2414S) | | |
| POLE (L1903L) | | |
| PMS2 (S260S) | | |
| BRCA2 (K1132K) | | |
| RAD51C (D253H) | | |
| POLE (A252V) | | |
| BRCA2 (V2466A) | | |
| ATM (D1853N) | | |
| BRCA1 (E1038G) | | |
| BRCA2 (V2171V) | | |
| EGFR (N158N) | | |
| ERBB2 (A356D) | | |
| BRCA1 (S694S) | | |
| BRCA1 (K1183R) | | |
| BRCA1 (L771L) | | |
| BRCA1 (D693N) | | |
| BRCA1 (P871L) | | |
| BRCA1 (S1436S) | | |
| BRIP1 (Y1137Y) | | |
| EGFR (T629T) | | |
| BRCA1 (S1613G) | | |
| KRAS (R161R) | | |
| ATM (N1983S) | | |
| BRCA1 (S1040N) | | |
| ERBB2 (P1140A) | | |
| POLD1 (K738N) | | |
| MSH3 (E949R) | | |
| BRCA2 (L1521L) | | |
| EGFR (A613A) | | |
| MSH6 (G39Q) | | |
| BRIP1 (Q879Q) | | |
| TP53 (R213R) | | |
| BRIP1 (S919P) | | |
| CDH1 (A692A) | | |
| EGFR (T903T) | | |
| KRAS (Q61H) | | |
| POLE (A1510A) | | |
| POLE (T1052T) | | |
| POLE (S2084S) | | |
| BARD1 (H506H) | | |
| APC (S1411S) | | |
| TP53 (P72R) | | |
| EGFR (Q787Q) | | |
| TP53 (R175H) | | |
| BARD1 (T351T) | | |
| PMS2 (K541E) | | |

**% of Similarity: 97.92 %**

| Genes | C2 Organoid | Tissue of origin |
|---|---|---|
| MSH2 (L556L) | | |
| PALB2 (T1100T) | | |
| BMPR1A (P2T) | | |
| PALB2 (E559R) | | |
| APC (E260) | | |
| MLH1 (I219V) | | |
| APC (T1556fs) | | |
| PMS2 (V280V) | | |
| PALB2 (G998Q) | | |
| MSH6 (D180D) | | |
| MSH6 (P92P) | | |
| POLD1 (T495T) | | |
| PIK3CA (I391M) | | |
| CTNNA1 (L180L) | | |
| MSH6 (R62R) | | |
| BRCA2 (V2466A) | | |
| KRAS (R162R) | | |
| BARD1 (A40V) | | |
| MSH3 (I78V) | | |
| BRCA2 (V1269V) | | |
| PALB2 (E672Q) | | |
| FANCM (I1460V) | | |
| ATM (S1270C) | | |
| BRCA2 (I3412V) | | |
| BRAF (G643G) | | |
| BRCA2 (V2171V) | | |
| MSH3 (Q949R) | | |
| FANCM (V878L) | | |
| ATM (N1983S) | | |
| EGFR (T629T) | | |
| BRAF (V600E) | | |
| ATM (D1853N) | | |
| MSH3 (A1045T) | | |
| POLE (P1548T) | | |
| BRCA2 (L1521L) | | |
| CTNNA1 (S740S) | | |
| EGFR (T903T) | | |
| FANCM (S175F) | | |
| BRIP1(E879E) | | |
| NBN (D399D) | | |
| BARD1 (P24S) | | |
| CDH1 (A692A) | | |
| BARD1 (H506H) | | |
| TP53 (R273H) | | |
| PMS2 (K541E) | | |
| EGFR (Q787Q) | | |
| EGFR (R521K) | | |
| FANCM (P1812A) | | |
| TP53 (P72R) | | |

**% of Similarity: 91.84 %**

**Supplementary Fig. S1. (G)** List of the gene mutations acquired in the TDOs compared to the tumor of origin (red boxes). The percentage of similarity is also reported. Passage numbers of the organoid lines were: C1: P11; C2: P13; C3: P10; C4: P14; C6: P10.

**G**

| Genes | C3 Organoid | Tissue of origin |
|---|---|---|
| PMS2 (K541E) | | |
| PMS2 (P470S) | | |
| PMS2 (S260S) | | |
| TP53 (R273C) | | |
| TP53 (P72R) | | |
| CDKN2A (A148T) | | |
| PALB2 (V932M) | | |
| KRAS (R161R) | | |
| BRCA2 (N372H) | | |
| BRCA2 (K1132K) | | |
| BRCA2 (L1521L) | | |
| BRCA2 (V2171V) | | |
| BRCA2 (S2414S) | | |
| BRCA2 (V2466A) | | |
| RAD51D (R24R) | | |
| ERBB2 (I625V) | | |
| ERBB2 (P1104A) | | |
| BRCA1 (S1613G) | | |
| BRCA1 (S1436S) | | |
| BRCA1 (K1183R) | | |
| BRCA1 (E1038G) | | |
| BRCA1 (P871L) | | |
| BRCA1 (L771L) | | |
| BRCA1 (S694S) | | |
| BRCA1 (D693N) | | |
| FANCM (S175F) | | |
| FANCM (V878L) | | |
| FANCM (I1460V) | | |
| FANCM (I1742V) | | |
| FANCM (P1812A) | | |
| MSH6 (R62R) | | |
| MSH6 (P92P) | | |
| MSH6 (D180D) | | |
| EGFR (N158N) | | |
| EGFR (T903T) | | |
| BRIP1 (Y1137Y) | | |
| BRIP1 (S919P) | | |
| BRIP1 (E879E) | | |
| CDH1 (A692A) | | |
| MSH3 (A55Adel) | | |
| MSH3 (A61Pdup | | |
| MSH3 (P67P69del) | | |
| MSH3 (I79V) | | |
| MSH3 (Q949R) | | |
| MSH3 (A1045T) | | |
| BMPR1A (P2T) | | |
| NBN (P672P) | | |
| NBN (D399D) | | |
| NBN (E185Q) | | |
| NBN (L34L) | | |
| ATM (N1983S) | | |
| APC (Q480) | | |
| APC (D774D) | | |
| APC (I1417fs) | | |
| APC (V1822D) | | |
| POLE (S2084S) | | |
| POLE (A1510A) | | |
| POLE (T1052T) | | |
| CTNNA1 (A179V) | | |
| PIK3CA (Q546R) | | |

**% of Similarity: 100 %**

| Genes | C4 Organoid | Tissue of origin |
|---|---|---|
| KRAS (R161R) | | |
| KRAS (G12S) | | |
| BRCA2 (K1132L) | | |
| BRCA2 (L1521L) | | |
| BRCA2 (T1915M) | | |
| BRCA2 (V2171V) | | |
| BRCA2 (S2414) | | |
| BRCA2 (V2466A) | | |
| MSH6 (R62R) | | |
| MSH6 (P92P) | | |
| MSH6 (D180D) | | |
| MSH6 (R1034Q) | | |
| SMAD4 (D351N) | | |
| POLD1 (T495T) | | |
| EGFR (T903T) | | |
| EGFR (D994D) | | |
| BRIP1 (Y1137Y) | | |
| BRIP1 (S919P) | | |
| BRIP1 (E879E) | | |
| CDH1 (T379M) | | |
| CDH1 (A692A) | | |
| MSH3 (A55A60del) | | |
| MSH3 (P67P69del) | | |
| MSH3 (I79V) | | |
| MSH3 (Q949R) | | |
| BMPR1A (P2T) | | |
| ATM (N1983S) | | |
| APC (Y486Y) | | |
| APC (A545A) | | |
| APC (S835fs) | | |
| APC (L1488) | | |
| APC (T1493T) | | |
| APC (G1678G) | | |
| APC (S1756S) | | |
| APC (V1822D) | | |
| APC (P1960P) | | |
| POLE (S2084S) | | |
| BRAF (G643G) | | |
| BARD1 (V507M) | | |

**% of Similarity: 87.18 %**

**Supplementary Fig. S1. (G)** List of the gene mutations acquired in the TDOs compared to the tumor of origin (red boxes). The percentage of similarity is also reported. Passage numbers of the organoid lines were: C1: P11; C2: P13; C3: P10; C4: P14; C6: P10.

**G**

| Genes | C6 organoid | Tissue of origin |
|---|---|---|
| PMS2 (P470S) | | |
| PMS2 (S260S) | | |
| TP53 (G245V) | | |
| TP53 (P72A) | | |
| CDKN2A (A148T) | | |
| KRAS (R161R) | | |
| KRAS (G12A) | | |
| BRCA2 (N372H) | | |
| BRCA2 (K1132K) | | |
| BRCA2 (L1521L) | | |
| BRCA2 (T1915M) | | |
| BRCA2 (V2171V) | | |
| BRCA2 (S2414S) | | |
| BRCA2 (V2466A) | | |
| RAD51D (R145C) | | |
| MLH1 (I219V) | | |
| ERBB2 (P1140A) | | |
| BRCA1 (S1613G) | | |
| BRCA1 (S1436S) | | |
| BRCA1 (K1183R) | | |
| BRCA1 (E1038G) | | |
| BRCA1 (P871L) | | |
| BRCA1 (L771L) | | |
| BRCA1 (S694S) | | |
| MUTYH (S515F) | | |
| MUTYH (Q338H) | | |
| MSH6 (R62R) | | |
| MSH6 (P92P) | | |
| MSH6 (D180D) | | |
| MSH6 (Y214Y) | | |
| SMAD4 (D315H) | | |
| EGFR (N158N) | | |
| EGFR (R521K) | | |
| EGFR (Q787Q) | | |
| EGFR (R836R) | | |
| EGFR (T903T) | | |
| BRIP1 (Y1137Y) | | |
| BRIP1 (S919P) | | |
| BRIP1 (E879E) | | |
| CDH1 (A692A) | | |
| MSH3 (A55A60del) | | |
| MSH3 (P67P69del) | | |
| MSH3 (I79V) | | |
| MSH3 (Q949R) | | |
| MSH3 (A1045T) | | |
| BMPR1A (P2T) | | |
| ATM (D1853N) | | |
| ATM (N1983S) | | |
| APC (Y486Y) | | |
| APC (A545A) | | |
| APC (L1129S) | | |
| APC (Q1406fs) | | |
| APC (T1493T) | | |
| APC (G1678G) | | |
| APC (S1756S) | | |
| APC (V1822D) | | |
| APC (P1960P) | | |
| POLE (S2084S) | | |
| POLE (A1510A) | | |
| POLE (N1396S) | | |
| POLE (T1052T) | | |
| CTNNA1 (S740S) | | |
| BRAF (G643G) | | |
| BARD1 (H506H) | | |
| BARD1 (T351T) | | |

**% of Similarity: 93.85 %**

**Supplementary Fig. S1. (G)** List of the gene mutations acquired in the TDOs compared to the tumor of origin (red boxes). The percentage of similarity is also reported. Passage numbers of the organoid lines were: C1: P11; C2: P13; C3: P10; C4: P14; C6: P10.

### Supplementary Figure S2

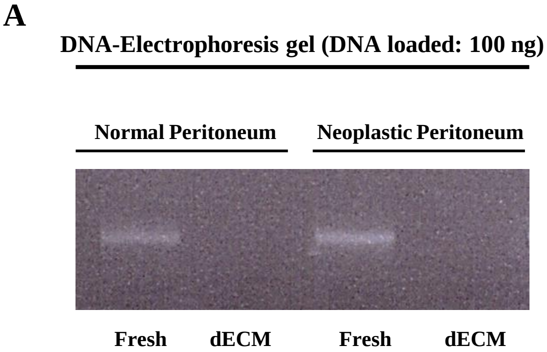


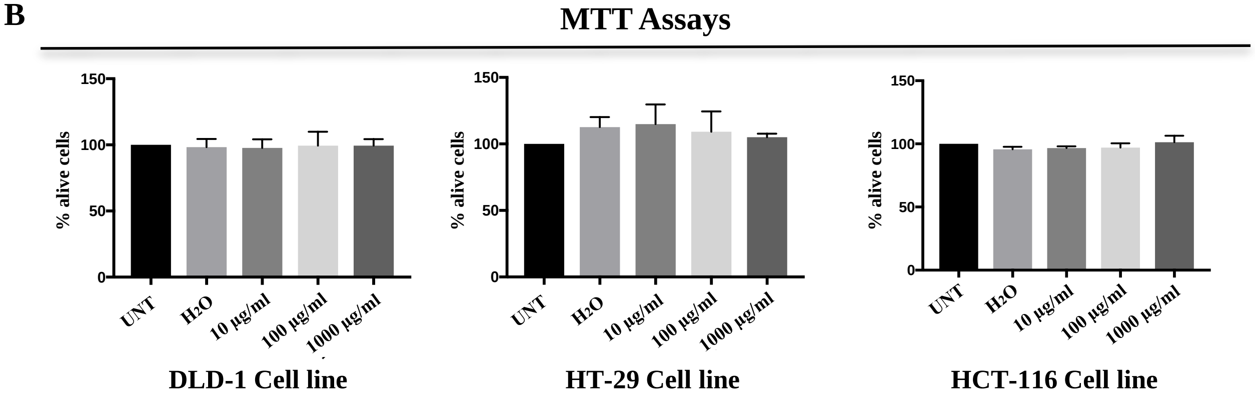


**Supplementary Fig. S2. (A)** To confirm the efficacy of the decellularization treatment, the loss of DNA content was assessed. DNA was extracted from 20 mg of non-decellularized or decellularized ECM and quantified with the Nanodrop instrument. Ten µl of DNA from PM or dECM were loaded onto a 1% agarose gel and the images were acquired using Gel Doc (Biorad). The results showed the absence of DNA in dECM samples, indicating the success of the decellularization procedure. The experiments were performed in triplicate. **(B)** Assessment of whether the components and procedures of the decellularization protocol can influence cell viability. CRC-derived DLD-1, HT-29 and HCT-116 cell lines were obtained from the American Type Culture Collection (ATCC, Rockville, MD, USA) and were authenticated by the Cell Culture Facility at the FIRC Institute of Molecular Oncology (IFOM, Milan, Italy) with the StemElite ID system (Promega). Cells were routinely tested for mycoplasma and cultured following the recommended ATCC's protocols. The 3D-dECMs were placed at − 80 °C overnight, lyophilized using freeze-drier cycles following standard procedures, and then added to the culture media of the three cell lines. During the experiments, three different concentrations of lyophilized 3D-dECMs were tested and cell viability was evaluated after 72 hours of growth by MTT (3-(4, 5-dimethylthiazol-2-yl)-2, 5-diphenyltetrazolium bromide) assay (Sigma Aldrich), following the manufacturer's instructions. The results showed no differences in cell viability between cells cultured with or without lyophilized ECM, highlighting that the decellularization procedure is safe for cell growth. The experiments were performed in duplicate.

### Supplementary Figure S3

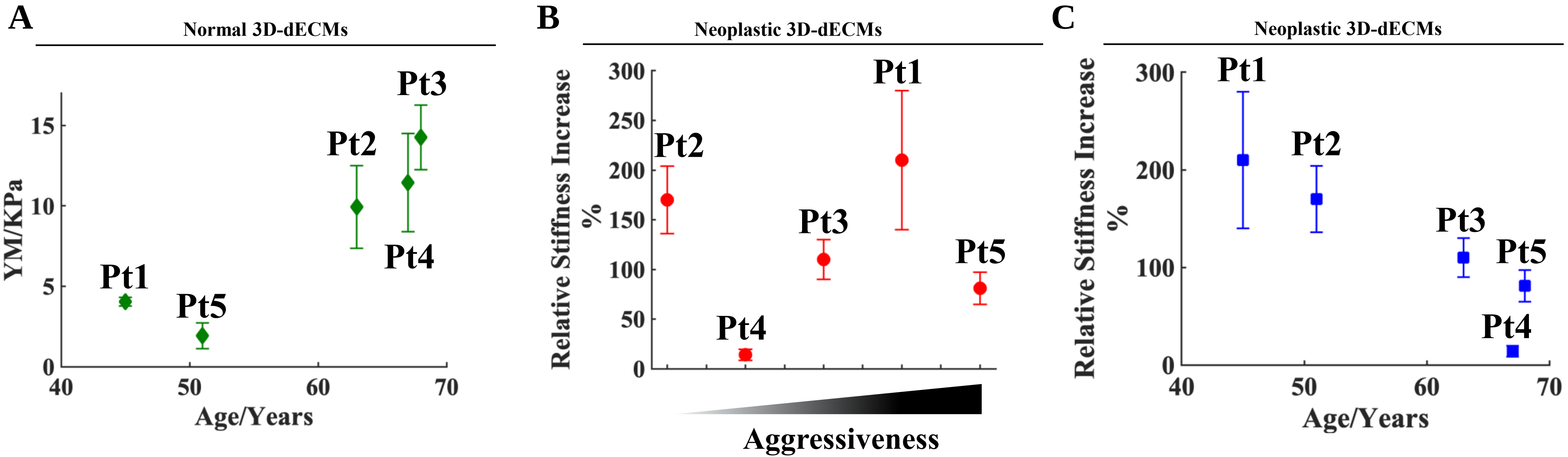


**Supplementary Fig. S3. (A)** YM values of normal-derived 3D-dECMs increases with the age of the patient. **(B)** The relative stiffening of neoplastic-derived 3D-dECMs (rel. increase of the YM) associated with tumor aggressiveness.

Tumor aggressiveness was estimated on the basis of patient's clinical and molecular characteristics and follow up (e.g., stage, grade, mutational profile, response to standard therapies and/or presence of recurrence). **(C)** The relative stiffening of neoplastic-derived 3D-dECMs associated with the age of the patient.

### Supplementary Figure S4

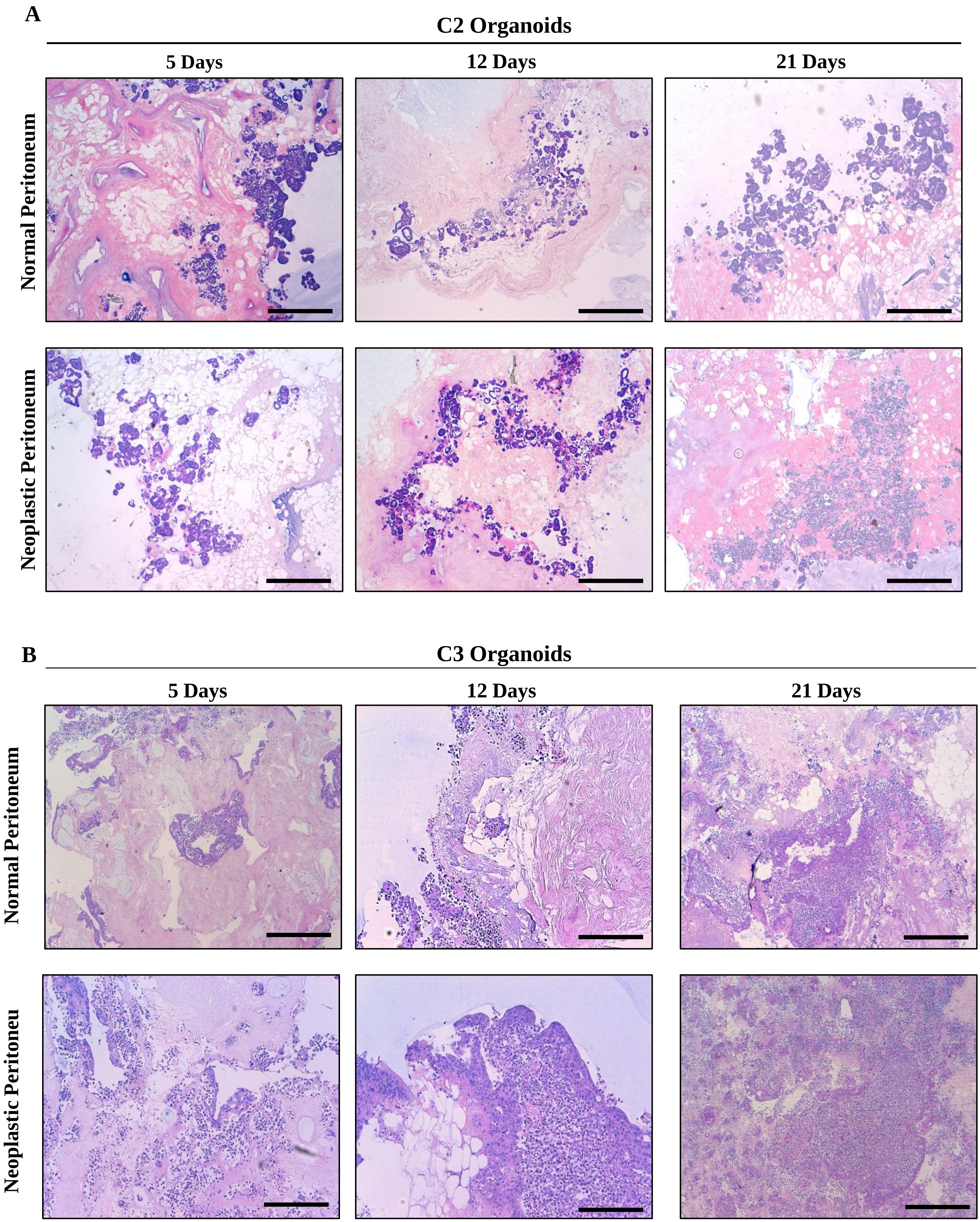


**Supplementary Fig. S4. (A)** H&E staining of decellularized matrices derived from normal (top) or neoplastic (bottom) peritoneum repopulated with PM-derived organoids (C2) at three different time points as indicated. Scale bar: 100 µm. The repopulation experiments were performed in triplicate. **(B)** H&E staining of decellularized matrices derived from normal (top) or neoplastic (bottom) peritoneum repopulated with PM-derived organoids (C3) on day 12. Scale bar: 100 µm. The repopulation experiments were performed in triplicate.

C

#### C2 Organoid

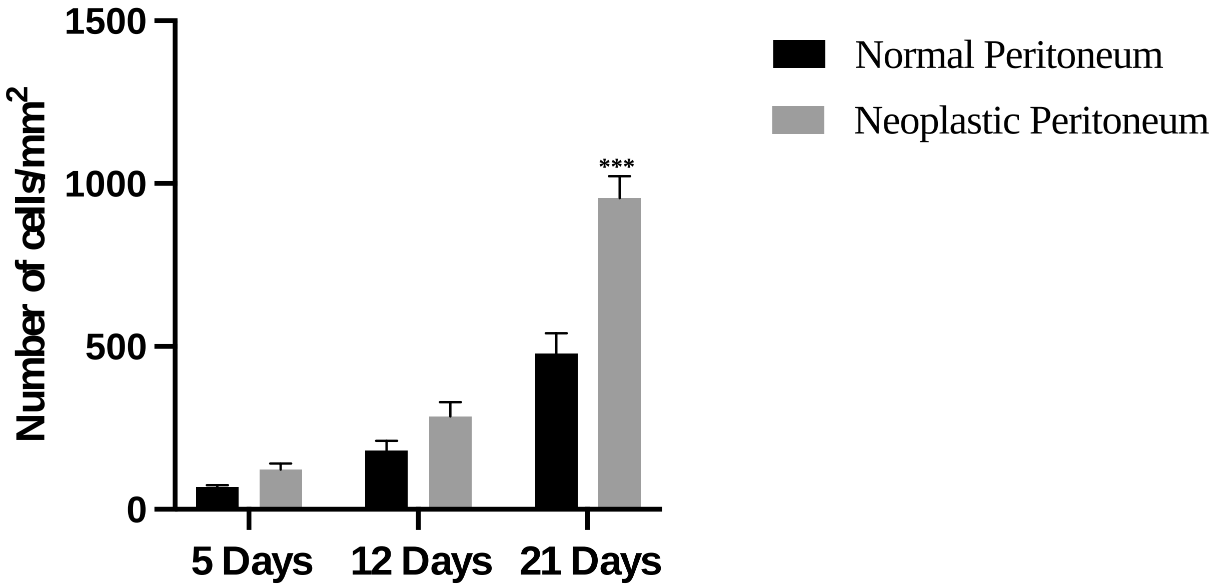


D

#### C3 Organoid

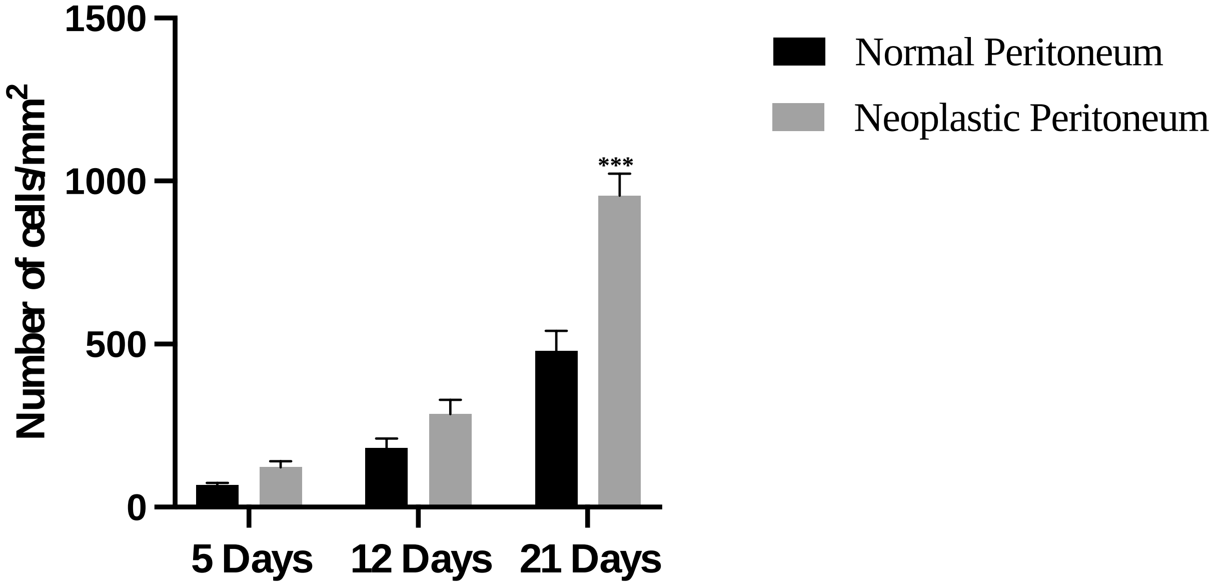


**Supplementary Fig. S4. (C)** Number of C2 PM-organoid derived cells grown onto normal and neoplastic-derived 3D-dECM per $mm^2$ after 5, 12 and 21 days. Three fields per experiments were counted using Qpath software. Data are presented as median and SD. One-way ANOVA (***$p$<0.001). **(D)** Number of C3 PM-organoid derived cells grown onto normal and neoplastic-derived 3D-dECM per $mm^2$ after 5, 12 and 21 days. Three fields per experiments were counted using Qpath software. Data are presented as median and SD for surgical specimens of three patients. One-way ANOVA (***$p$<0.001). The repopulation experiments were performed in triplicate.

E

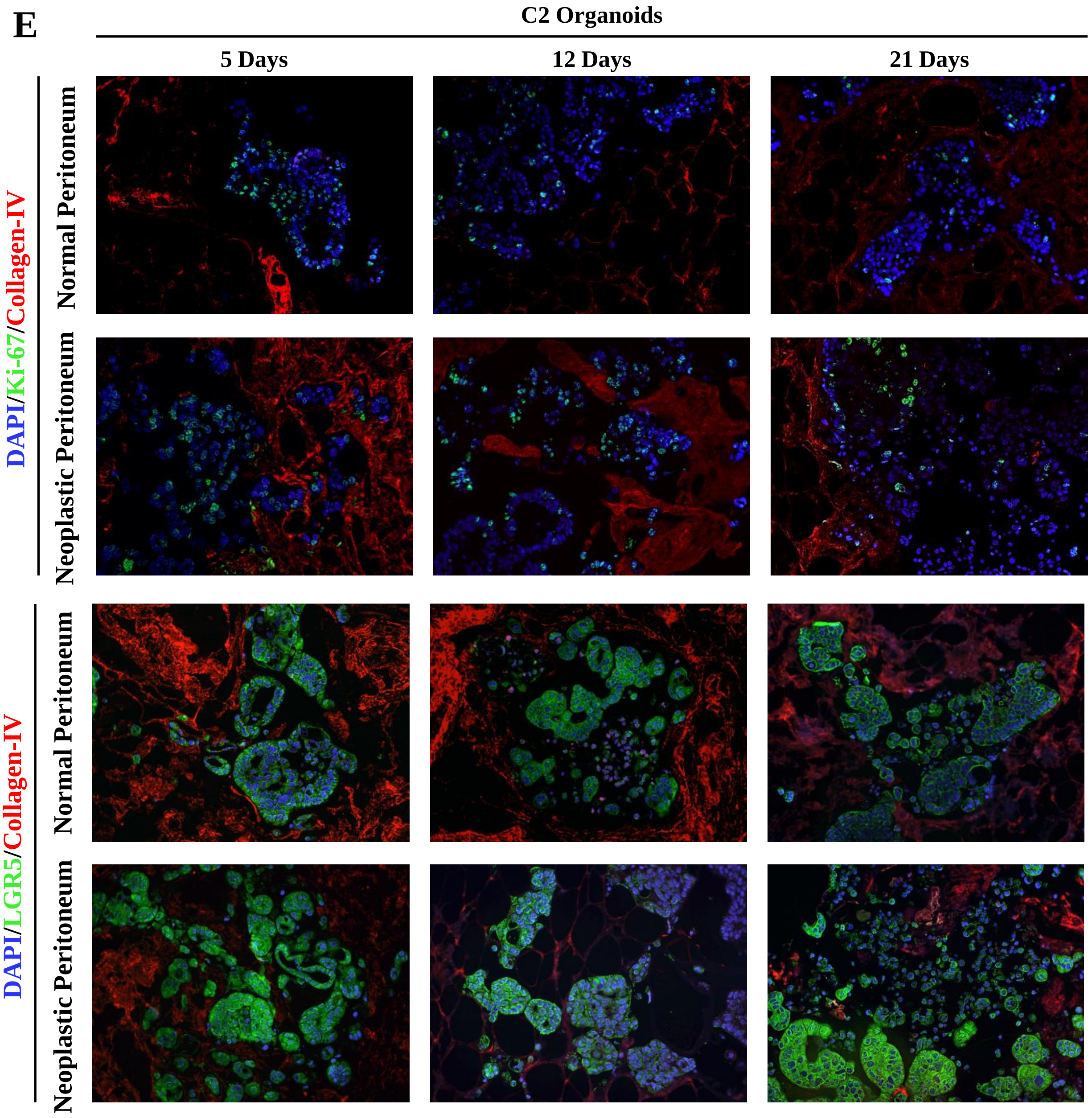


F

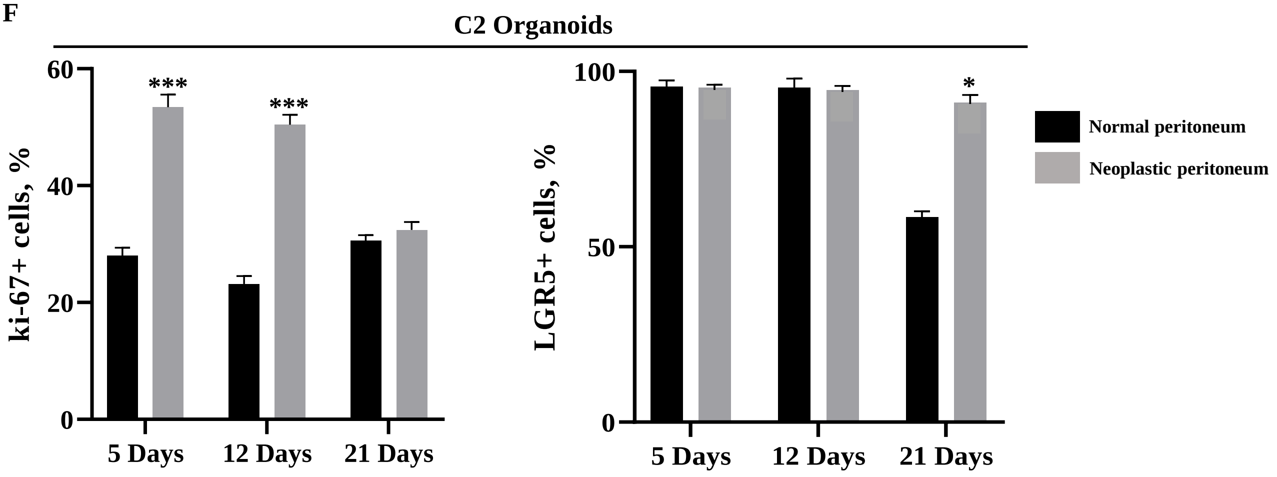


**Supplementary Fig. S4. (E) Top panel:** IF analysis of 3D-dECMs derived from normal (top) and neoplastic (bottom) peritoneum repopulated with PM-derived organoids (C2) at different time points as indicated, using Ki-67 (green) and collagen IV (red) antibodies. Scale bar: 50 µm. The experiments were performed in triplicate. **Bottom panel:** IF analysis of 3D-dECMs derived from normal (top) and neoplastic (bottom) peritoneum repopulated with PM-derived organoids (C2) at different time points as indicated, using LGR5 (green) and collagen IV (red) antibodies. The samples were counterstained with DAPI (blue) Scale bar: 50 µm. The experiments were performed in triplicate. **(F) Left panel:** proliferation rate of PM-derived organoids, measured as the percentage of Ki-67$^{+}$ cells present in fields devoid of dead cells. Five fields per experiment (40X magnification) were counted. Data are presented as median and SD. One-way ANOVA (***$p$<0.001). **Right panel:** amount of stem cells in PM-derived organoids, measured as the percentage of LGR5$^{+}$ cells present in fields devoid of dead cells. Five fields per experiment (40X magnification) were counted. Data are presented as median and SD. One-way ANOVA (*$p$<0.05). The experiments were performed in triplicate.

G

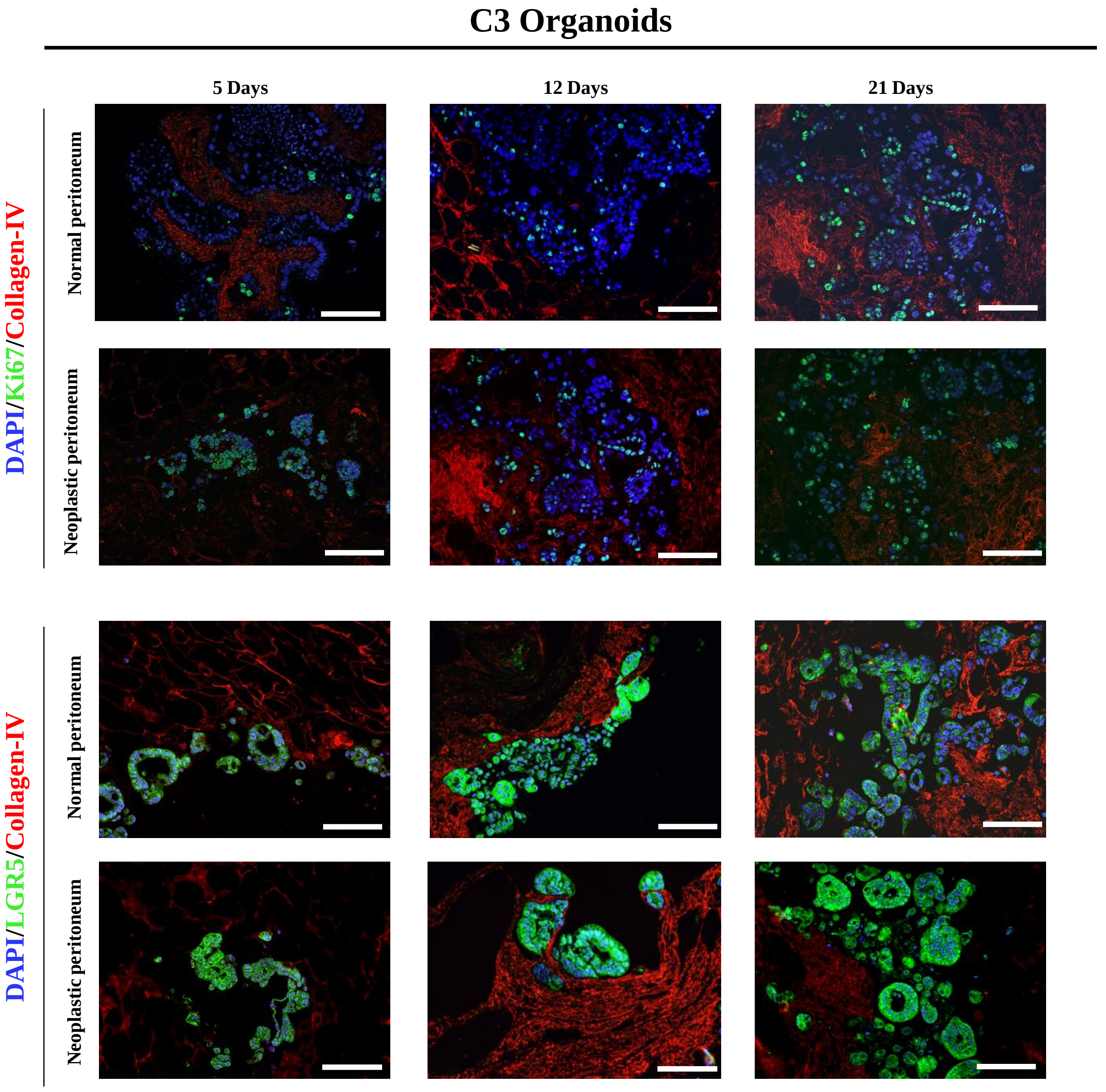


H

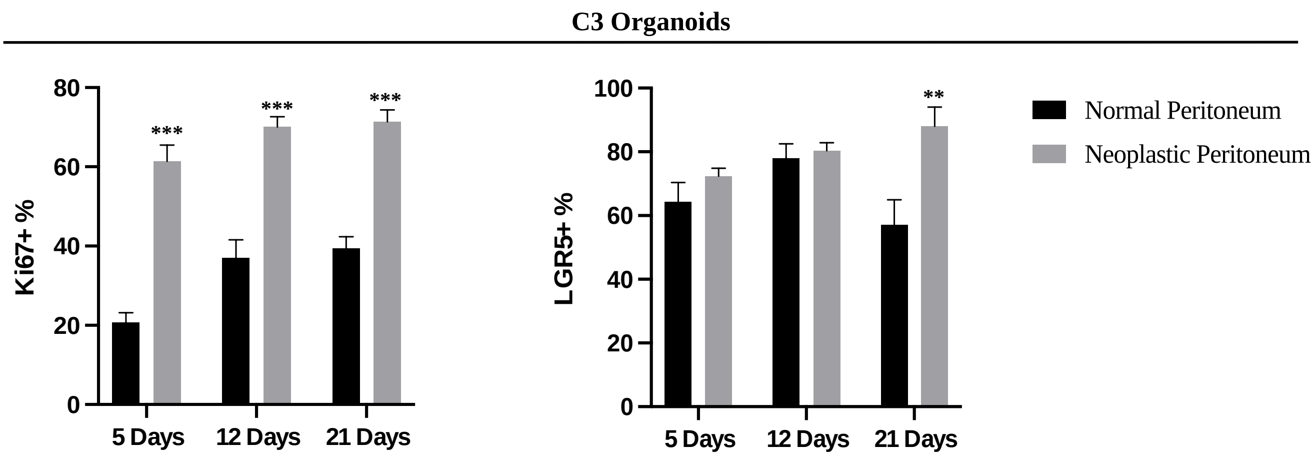


**Supplementary Fig. S4. (G) Top panel:** IF analysis of 3D decellularized matrices derived from normal (top) and neoplastic (bottom) peritoneum repopulated with PM-derived organoids (C3), using Ki-67 (green) and collagen IV (red) (left panel). **Bottom panel:** LGR5 (green) and collagen IV (red) (right panel) antibodies. The samples were counterstained with DAPI (blue). Scale bar: 50 µm. The experiments were performed in triplicate. **(H) Left panel:** proliferation rate of PM-derived organoids measured as the percentage of Ki-67$^{+}$ cells present in fields devoid of dead cells. Five fields per experiment (40X magnification) were counted. Data are presented as median and SD. Student's *t-test* (***p*<0.01). **Right panel:** percentage of stem cells in PM-derived organoids, measured as the percentage of LGR5$^{+}$ cells present in fields devoid of dead cells. Five fields per experiment (40X magnification) were counted. Data are presented as median and SD. Student's *t-test* (**p*<0.05). The experiments were performed in triplicate.

### Supplementary Figure S5

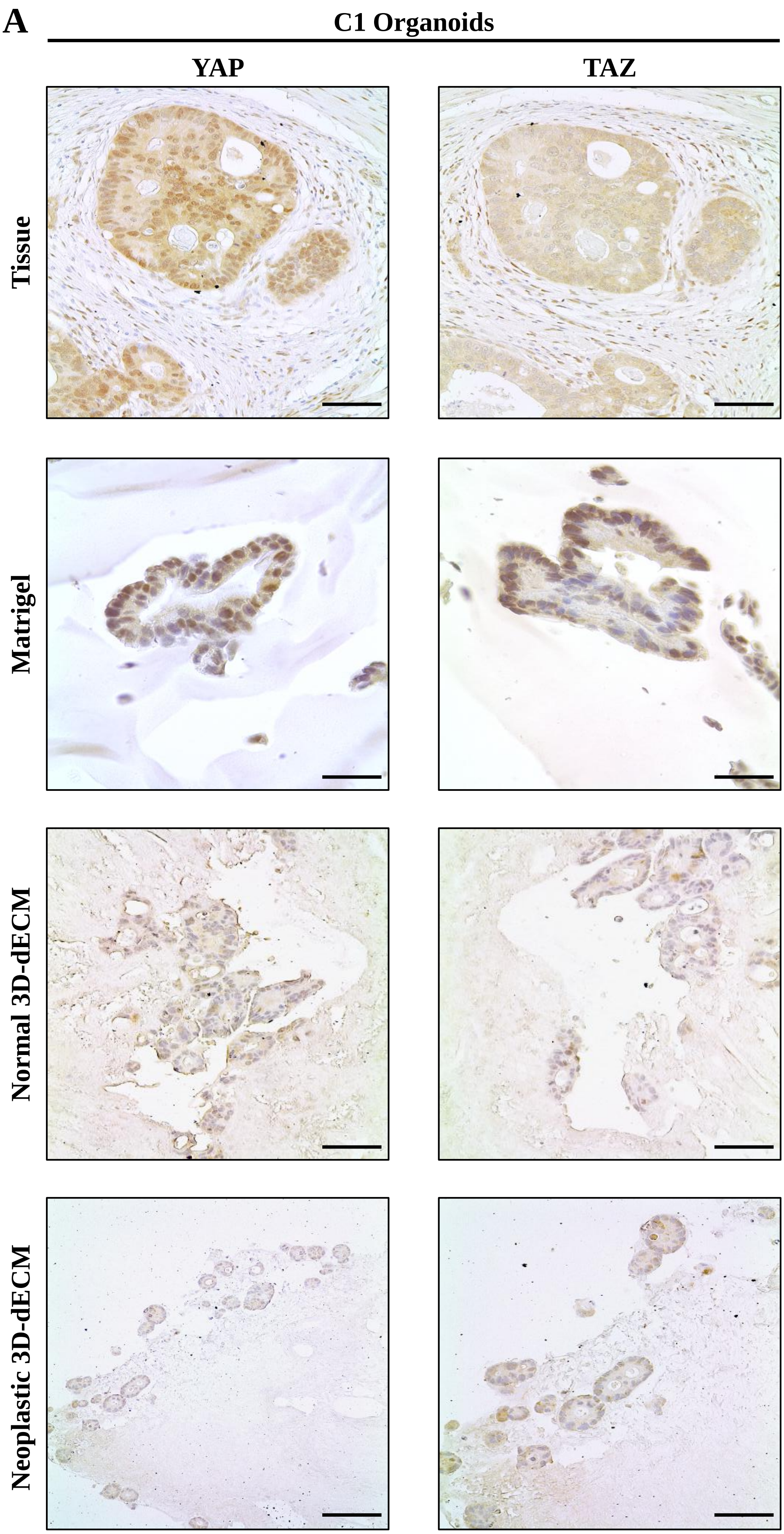


**Supplementary Fig. S5. (A)** Comparative immunohistochemical images of PM-derived organoids (C1) grown on different substrates (Matrigel, Normal 3D-dECM and Neoplastic 3D-dECM) and their corresponding tumor of origin. Expression of YAP and TAZ proteins was analysed. Scale bar: 100 µM.

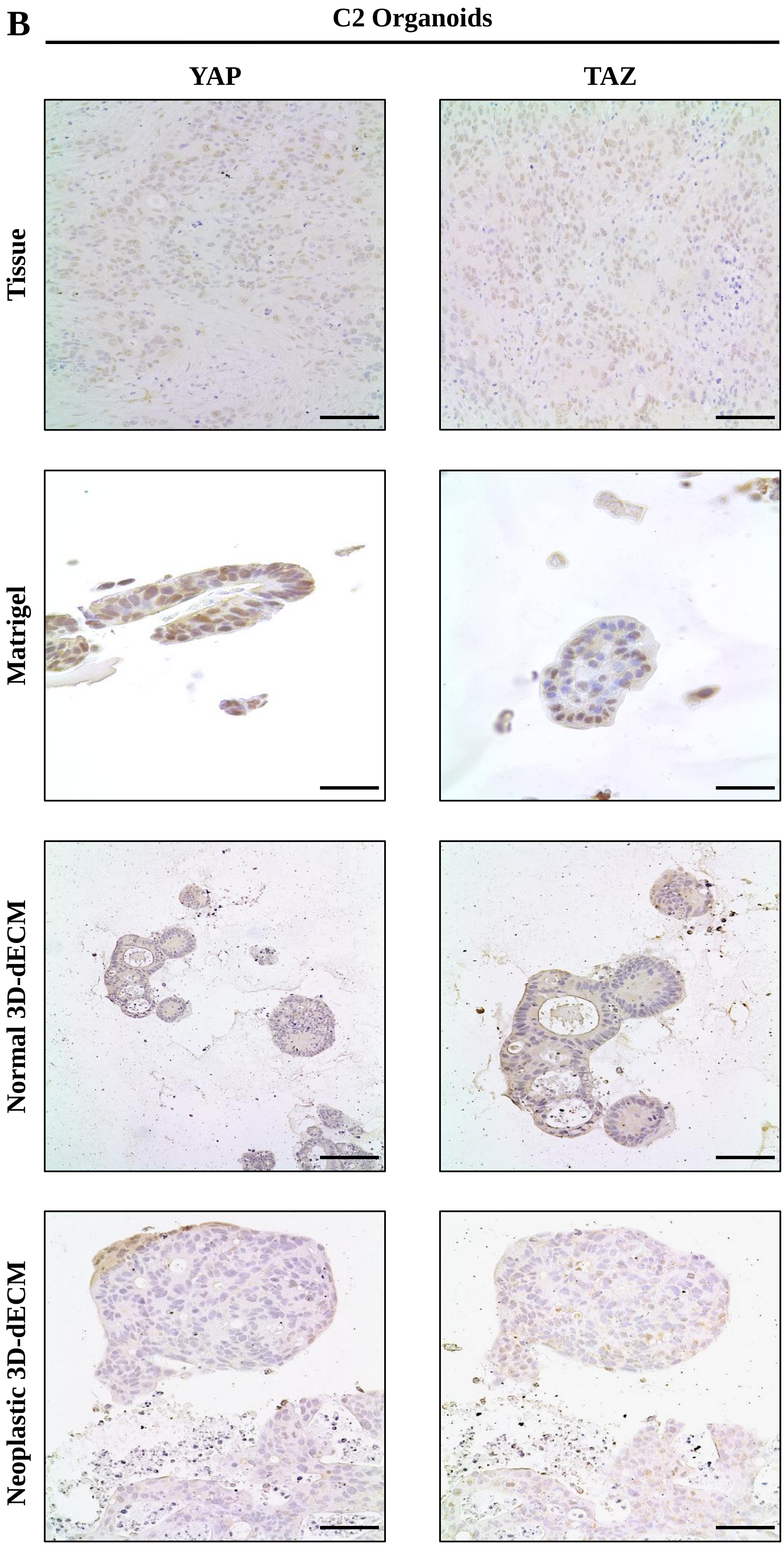


**Supplementary Fig. S5. (B)** Comparative immunohistochemical images of PM-derived organoids (C2) grown on different substrates (Matrigel, Normal 3D-dECM and Neoplastic 3D-dECM) and their corresponding tumor of origin. Expression of YAP and TAZ proteins was analyzed. Scale bar: 100 μM.

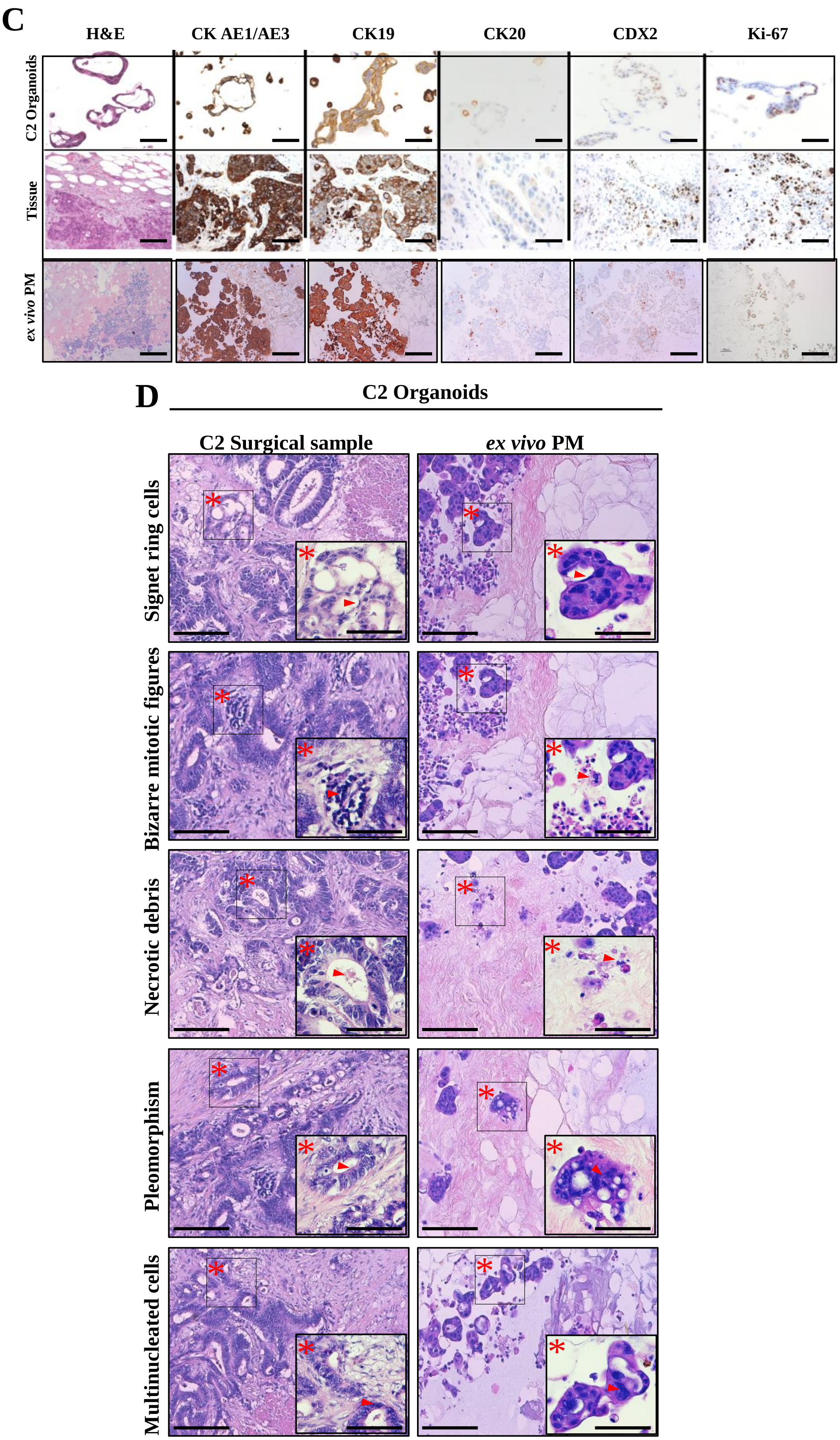


**Supplementary Fig. S5. (C)** Comparative histological and immunohistochemical analysis of C2 PM-derived organoids Vs their corresponding tumor of origin and the *ex vivo* engineered PM lesion. Samples were analyzed for the expression of the CRC-specific markers as indicated. Scale bar: 50 µM. Images in the first two lanes were previously published [11]. **(D)** Histological analysis of PM and neoplastic-derived 3D-dECM repopulated with C2 PM-derived organoids. The *ex vivo* engineered PM lesions present histological features that are typical of PMs of gastrointestinal origin. Asterisks and arrows indicated the main morphological features. Scale bar: 20 µm.

**E**

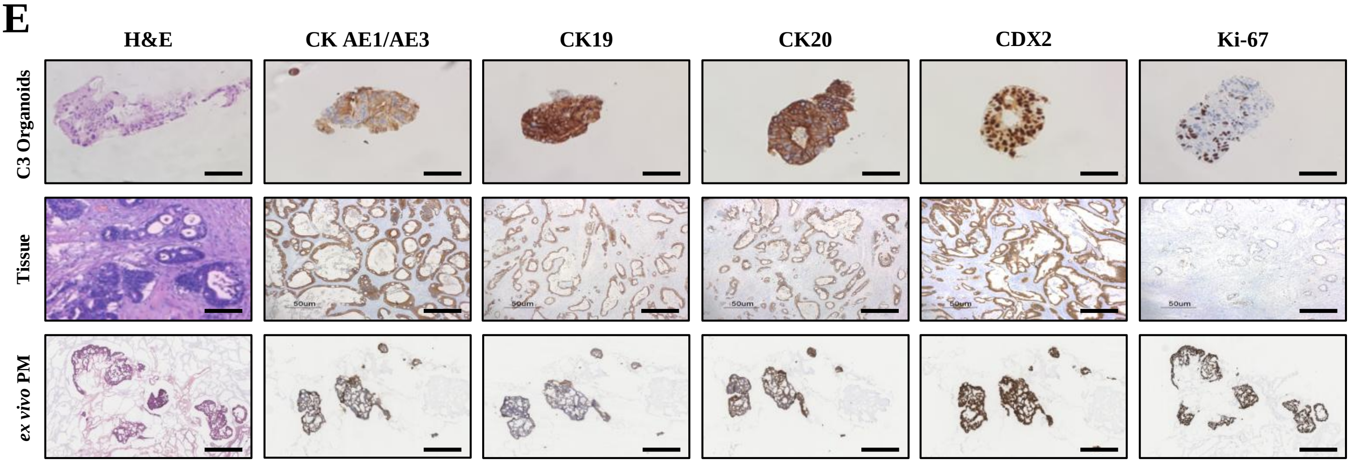


**F**

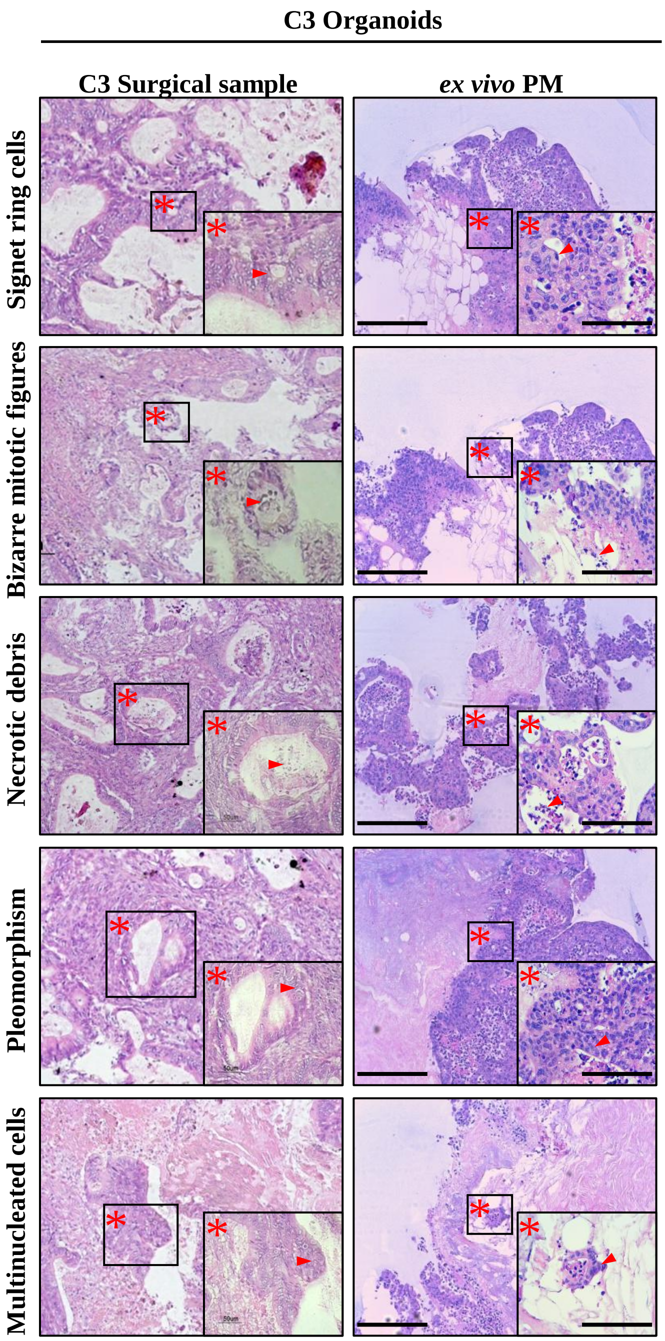


**Supplementary Fig. S5. (E)** Comparative histological and immunohistochemical analysis of C3 PM-derived organoids Vs their corresponding tumor of origin and the *ex vivo* engineered PM lesion. Samples were analyzed for the expression of the CRC-specific markers as indicated. Scale bar: 50 μm. **(F)** Histological analysis of peritoneal metastasis and neoplastic-derived 3D-dECMS repopulated with C3 PM-derived organoids. The *ex vivo* engineered PM lesions present histological features that are typical of PMs of gastrointestinal origin. Asterisks and arrows indicated the main morphological features. Scale bar: 20 μm.

**G**

**C2 Organoids**

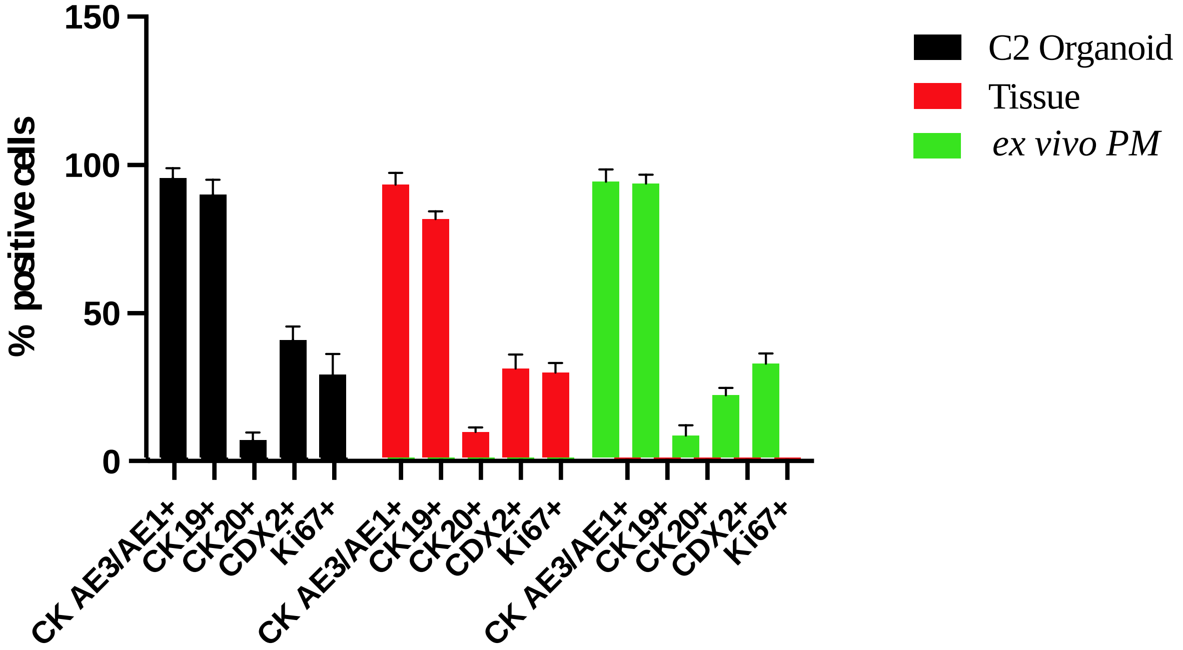


**H**

**C3 Organoids**

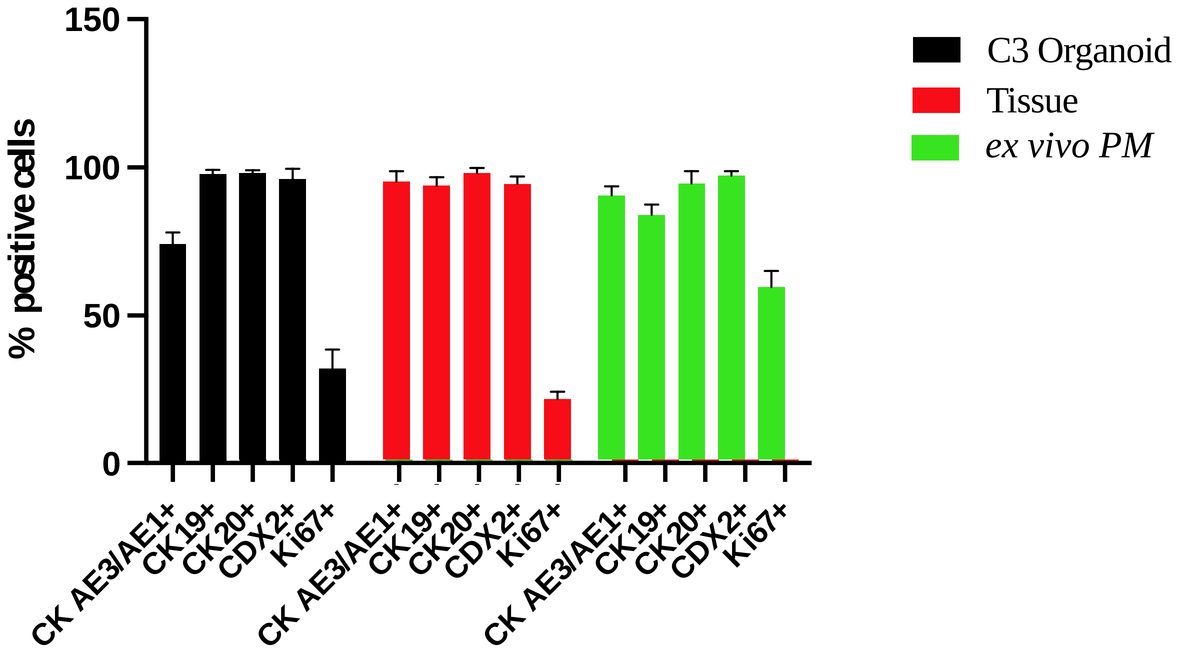


**Supplementary Fig. S5. (G)** Quantitative counts of the percentage of CK AE/AE3, CK19, CK20, CDX2 and Ki-67 positive cells in C2 PM-derived organoids Vs their corresponding tumor of origin and the *ex vivo* engineered PM lesion Three fields per experiments were counted using Qpath software. Data are presented as median and SD. One-way ANOVA did not show differences between the two groups. **(H)** Quantitative counts of the percentage of CK AE/AE3, CK19, CK20, CDX2 and Ki-67 positive cells in C3 PM-derived organoids Vs their corresponding tumor of origin and the *ex vivo* PM engineered lesion. Three fields per experiments were counted using Qpath software. Data are presented as median and SD. One-way ANOVA did not show differences between the two groups.

### Supplementary Figure S6

**A**

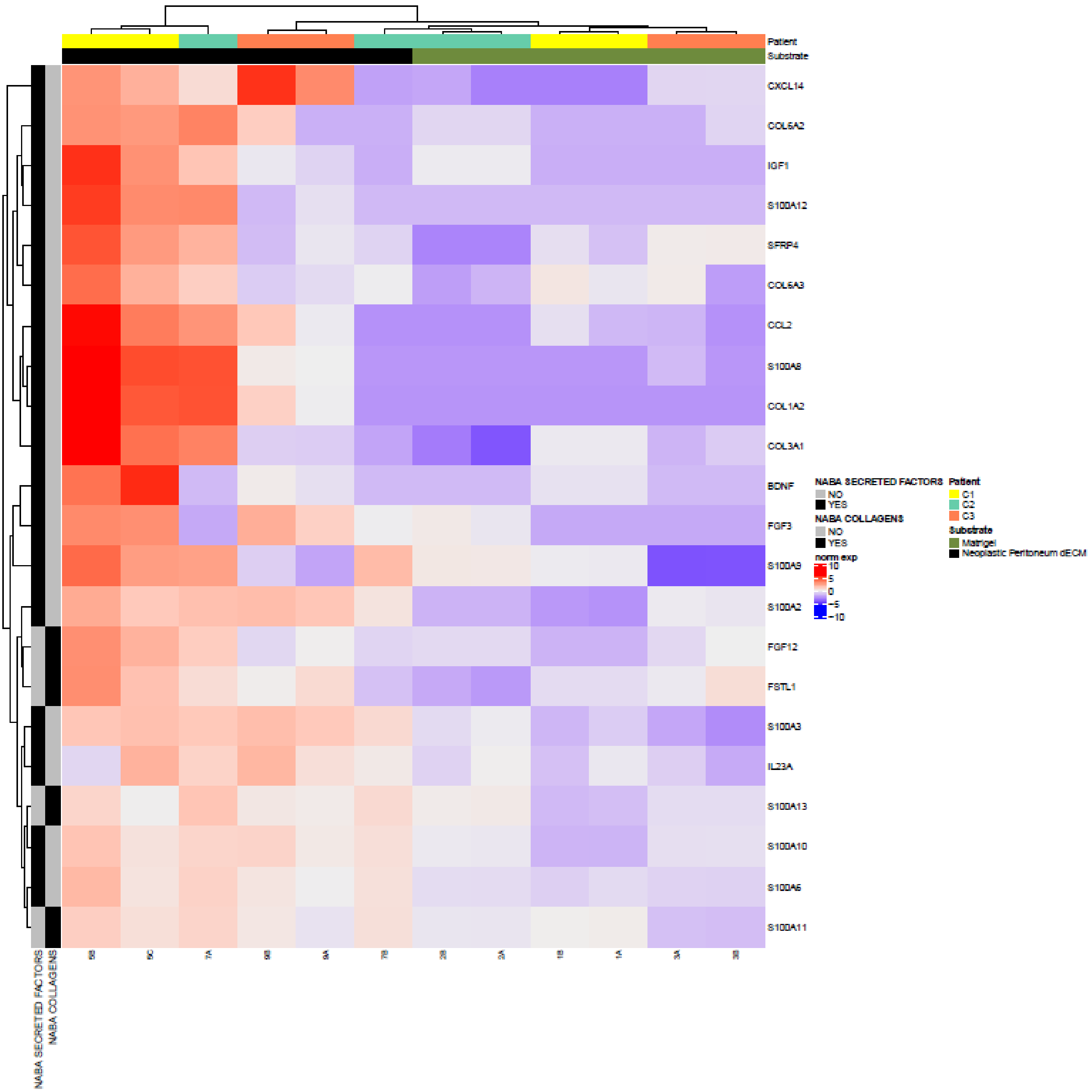


**Supplementary Fig. S6. (A)** Unsupervised hierarchical clustering of organoids based on the expression of the top DEGs between organoids grown on neoplastic 3D-dECM and in Matrigel, and present in Naba Secreted Factors or Naba Collagen categories.

**B**

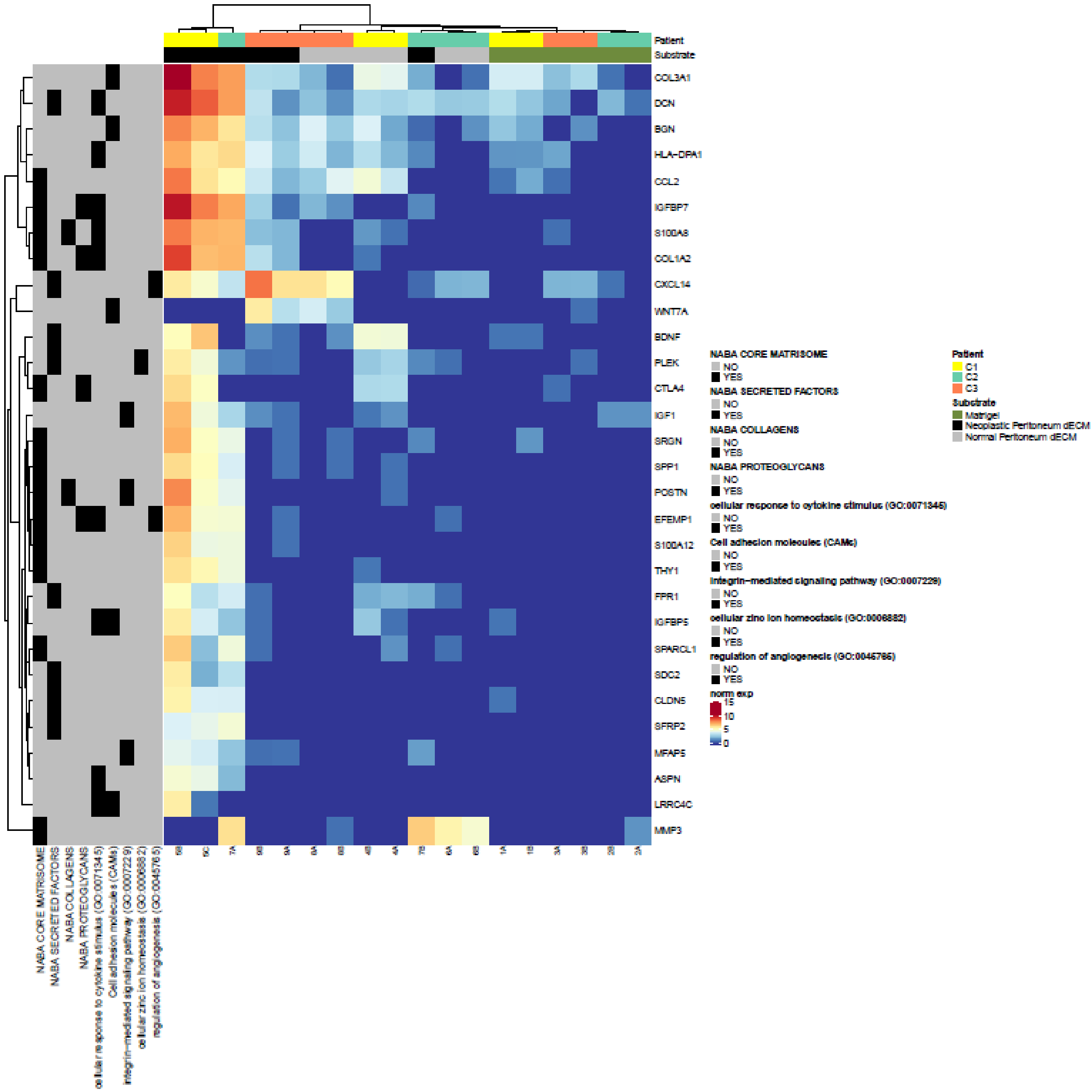


**Supplementary Fig. S6. (B)** Unsupervised hierarchical clustering of organoids based on the expression of the top DEGs between organoids grown on neoplastic 3D-dECM, normal 3D-dECM and in Matrigel, and present in Naba Core Matrisome, Naba Secreted Factors, Naba Collagen categories, Naba Proteoglycan categories, cellular response to cytokine stimulus, cell adhesion molecules, integrin mediated signaling pathway, cellular zinc ion homeostasis, and regulation of angiogenesis categories.

**C**

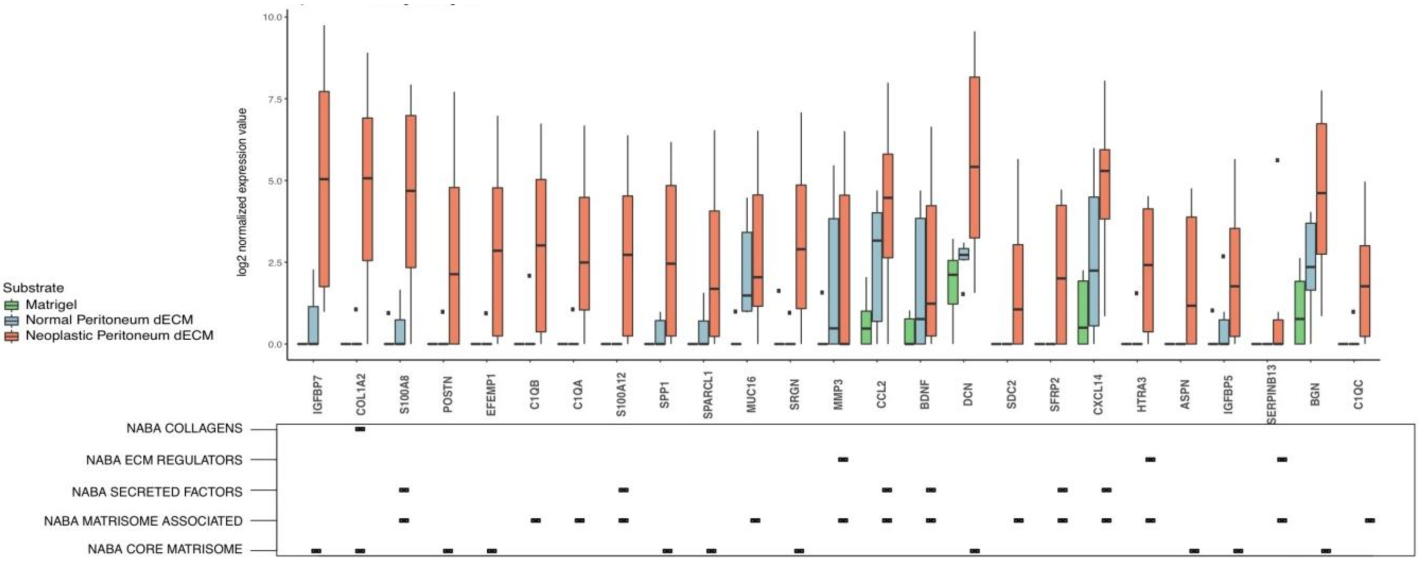


**D**

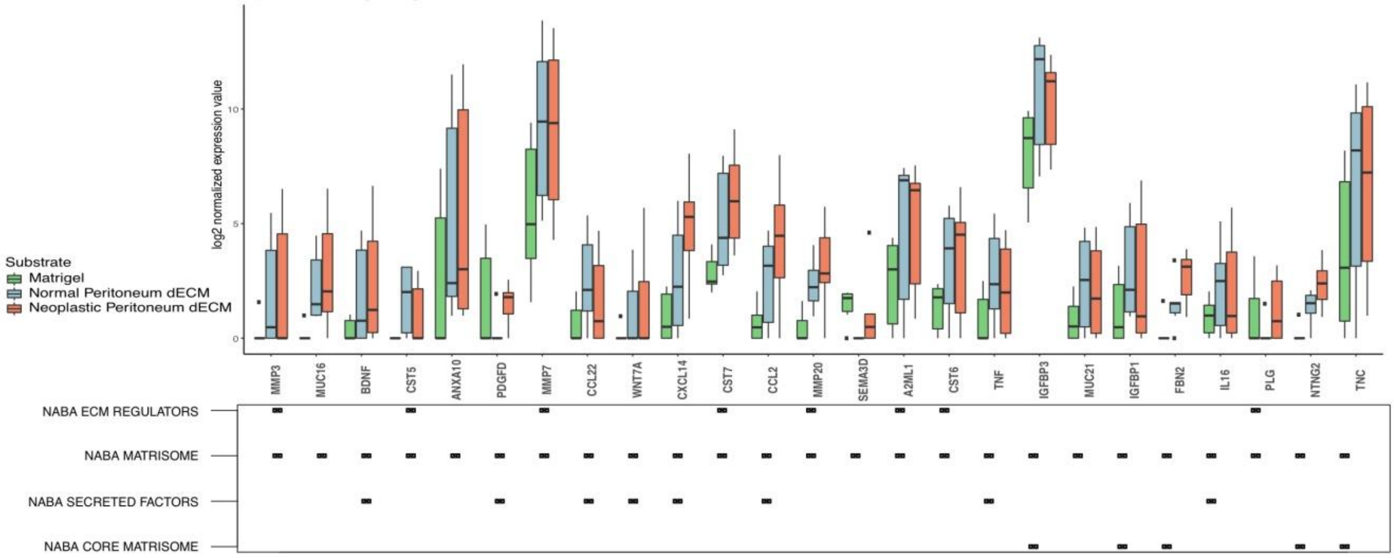


**Supplementary Fig. S6. (C)** Box plot showing the expression in organoids grown on neoplastic 3D-dECM and in Matrigel, of DEGs selected based on their involvement in the indicated processes of the Naba Matrisome geneset. DEGs expression in organoids grown on normal 3D-dECM is also shown. Median and interquartile range are displayed as horizontal lines. Black squares in the bottom panel indicate which category the genes belong to. **(D)** Box plot showing the expression, in organoids grown on normal 3D-dECM and in Matrigel, of DEGs selected based on their involvement in the indicated processes of the Naba Matrisome geneset. DEGs expression in organoids grown on neoplastic 3D-dECM is also shown. Median and interquartile range are displayed as horizontal lines. Black squares in the bottom panel indicate which category the genes belong to.

**E**

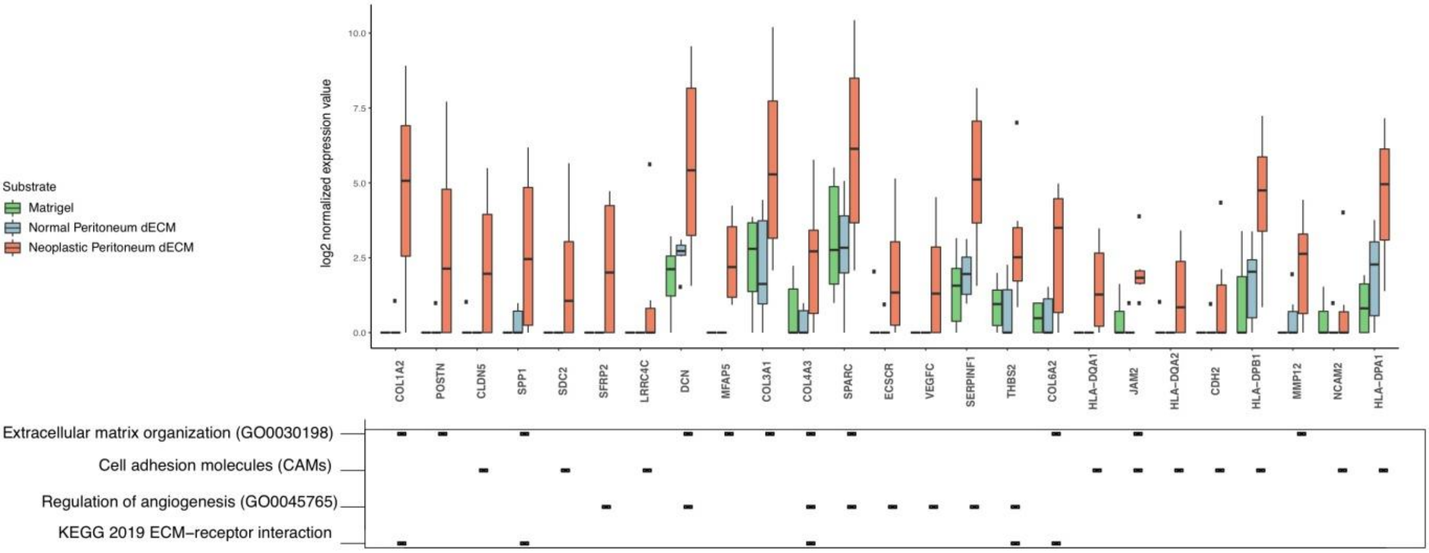


**F**

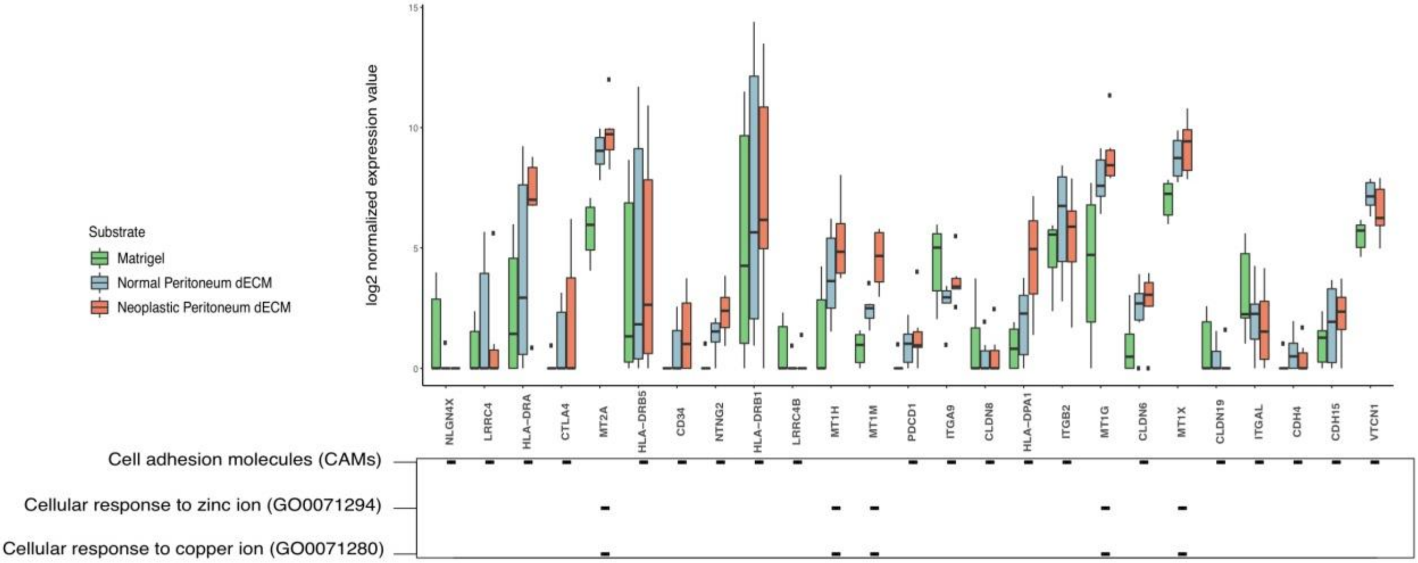


**Supplementary Fig. S6. (E)** Box plot showing the expression in organoids grown on normal 3D-dECMs and in Matrigel, of DEGs selected based on their involvement in the indicated processes of GO Biological Process, KEGG and Reactome databases. DEGs expression in organoids grown on neoplastic 3D-dECM is also shown. Median and interquartile range are displayed as horizontal lines. Black squares in the bottom panel indicate which category the genes belong to. **(F)** Box plot showing the expression levels of genes selected based on their involvement in the indicated processes. The expression levels of the selected genes were compared in organoids grown in Matrigel, on neoplastic 3D-dECM and on normal 3D-dECM. Median and interquartile range are displayed as horizontal lines. Black squares in the bottom panel indicate which category the genes belong to.

**G**

| | Collection | ID | Term | Adjusted *p-value* | Leading Edge genes (%) | # Genes |
|---|---|---|---|---|---|---|
| Neoplastic 3D-dECM Vs Matrigel | GO BP | GO:0071345 | Cellular response to cytokine stimulus | 0.003365 | 27.1 | 413 |
| | GO BP | GO:0045765 | Regulation of angiogenesis | 0.003365 | 19.3 | 171 |
| | GO BP | GO:0006882 | Cellular zinc ion homeostasis | 0.003365 | 46.7 | 30 |
| | GO BP | GO:0071280 | Cellular response to copper ion | 0.003365 | 55.0 | 20 |
| | GO BP | GO:0007229 | Integrin-mediated signaling pathway | 0.011230 | 19.6 | 56 |
| | KEGG | | Wnt signaling pathway | 0.022987 | 20.5 | 151 |
| Normal 3D-dECM Vs Matrigel | GO BP | GO:0071280 | Cellular response to copper ion | 0.006015 | 55.0 | 20 |
| | GO BP | GO:0071294 | Cellular response to zinc ion | 0.006015 | 64.7 | 17 |
| | KEGG | | Cell adhesion molecules (CAMs) | 0.006015 | 32.8 | 128 |
| Neoplastic 3D-dECM Vs Normal 3D-dECM | KEGG | | ECM-receptor interaction | 0.004499 | 25.0 | 80 |
| | GO BP | GO:0045765 | Regulation of angiogenesis | 0.009176 | 15.2 | 171 |
| | KEGG | | Cell adhesion molecules (CAMs) | 0.020511 | 22.7 | 132 |
| | Reactome | R-HSA-5660526 | Response to metal ions Homo sapiens | 0.040965 | 81.8 | 11 |
| | GO BP | GO:0030198 | Extracellular matrix organization | 0.002740 | 25.3 | 221 |

**H**

| | Collection | Term | Adjusted *p-value* | Leading Edge genes (%) | # Genes |
|---|---|---|---|---|---|
| Neoplastic 3D-dECM Vs Matrigel | NABA MATRISOME | Naba Matrisome Associated | 0.003365 | 38.2 | 663 |
| | NABA MATRISOME | Naba Core Matrisome | 0.003365 | 36.1 | 266 |
| | NABA MATRISOME | Naba ECM Regulators | 0.003365 | 30.6 | 222 |
| | NABA MATRISOME | Naba Secreted Factors | 0.003365 | 45.1 | 288 |
| | NABA MATRISOME | Naba Collagens | 0.004307 | 31.2 | 44 |
| Normal 3D-dECM Vs Matrigel | NABA MATRISOME | Naba ECM Regulators | 0.0060150 | 39.2 | 217 |
| | NABA MATRISOME | Naba Core Matrisome | 0.006015 | 22.7 | 256 |
| | NABA MATRISOME | Naba Secreted Factors | 0.006015 | 34.7 | 274 |
| | NABA MATRISOME | Naba Matrisome | 0.006024 | 32.9 | 897 |
| Neoplastic 3D-dECM Vs Normal 3D-dECM | NABA MATRISOME | Naba Core Matrisome | 0.002740 | 26.3 | 266 |
| | NABA MATRISOME | Naba Secreted Factors | 0.002740 | 39.2 | 291 |
| | NABA MATRISOME | Naba ECM Glycoproteins | 0.002740 | 21.9 | 192 |
| | NABA MATRISOME | Naba Collagens | 0.002740 | 40.9 | 44 |
| | NABA MATRISOME | Naba Proteoglycans | 0.002740 | 53.3 | 30 |
| | NABA MATRISOME | Naba ECM Regulators | 0.024908 | 28.2 | 16 |

**Supplementary Fig. S6. (G)** Selected categories enriched in DEGs between organoids grown on neoplastic 3D-dECM, normal 3D-dECM and in Matrigel involved in relevant biological processes according to GO Biological Process, KEGG and Reactome databases. **(H)** Selected Matrisome categories enriched in DEGs between organoids grown on neoplastic 3D-dECM, normal 3D-dECM and in Matrigel involved in relevant biological processes according to Naba Maatrisome dataset.

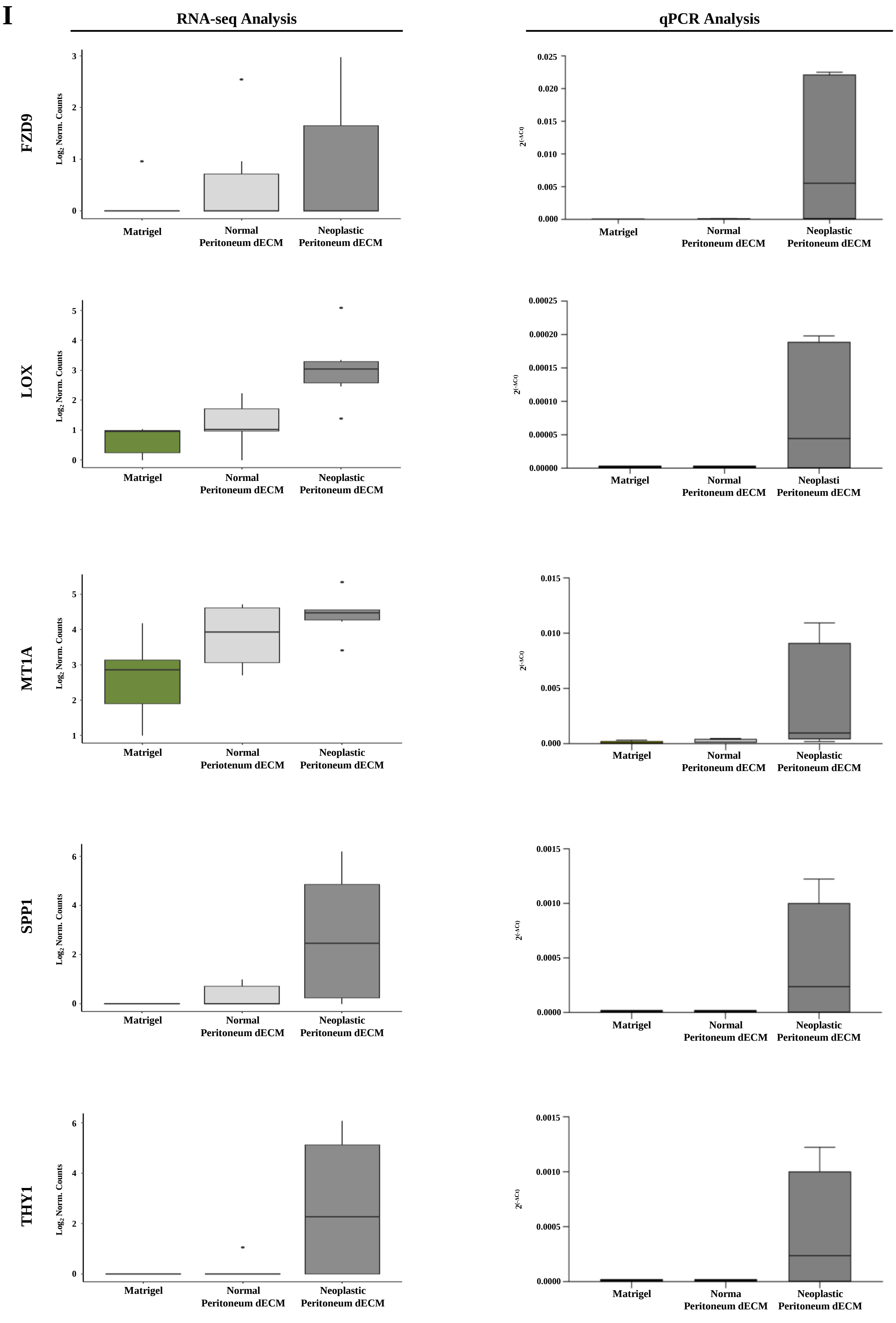
I
RNA-seq Analysis
qPCR Analysis
FZD9
LOX
MT1A
SPP1
THY1
Log2 Norm. Counts
2(-ΔCt)
Matrigel
Normal
Peritoneum dECM
Neoplastic
Peritoneum dECM
Periotenum dECM
Neoplasti
Norma

**L**

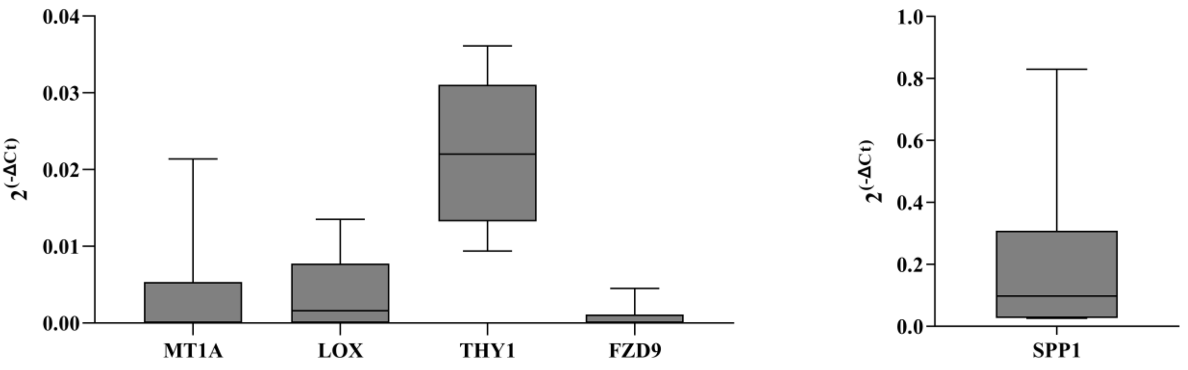


**Supplementary Fig. S6. (I)** Expression levels of MT1A, LOX, THY1, FZD9 and SPP1 in C1, C2 and C3 grown into Matrigel and on normal and neoplastic-derived 3D-dECMs (left panel). Values are calculated as $log_2$ of the total normal counts. The horizontal line inside the box indicates the median and the whiskers indicate the extreme measured values.

Expression levels of MT1A, LOX, THY1, FZD9 and SPP1 genes on organoid cultures grown in Matrigel and on normal or neoplastic peritoneal 3D-dECMs (right panel). Values are calculated as $2(^{-\Delta Ct})$ and normalized to GAPDH. The horizontal line inside the box indicates the median and the whiskers indicate the extreme measured values. **(L)** Expression levels of MT1A, LOX, THY1, FZD9 and SPP1 genes on FFPE tissues from PM-derived organoids. Values are calculated as $2(^{-\Delta Ct})$ and normalized to GAPDH. The horizontal line inside the box indicates the median and the whiskers indicate the extreme measured values.

### Supplementary Figure S7

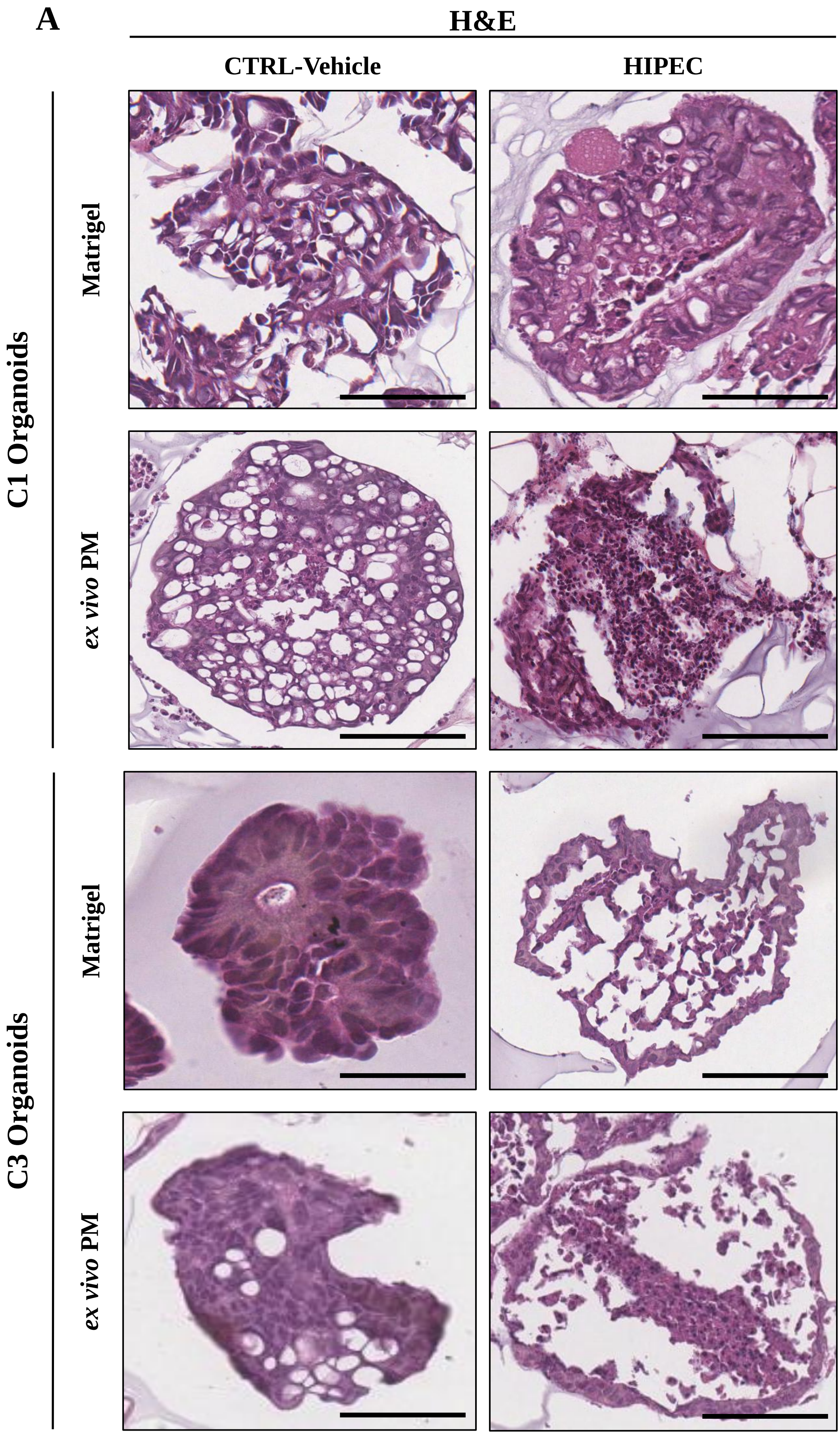


**Supplementary Fig. S7. (A)** H&E staining of C1 and C3 organoids cultured in Matrigel and on neoplastic-derived peritoneal 3D-dECMs after *in vitro* HIPEC treatments. Scale bar: 50 µm.

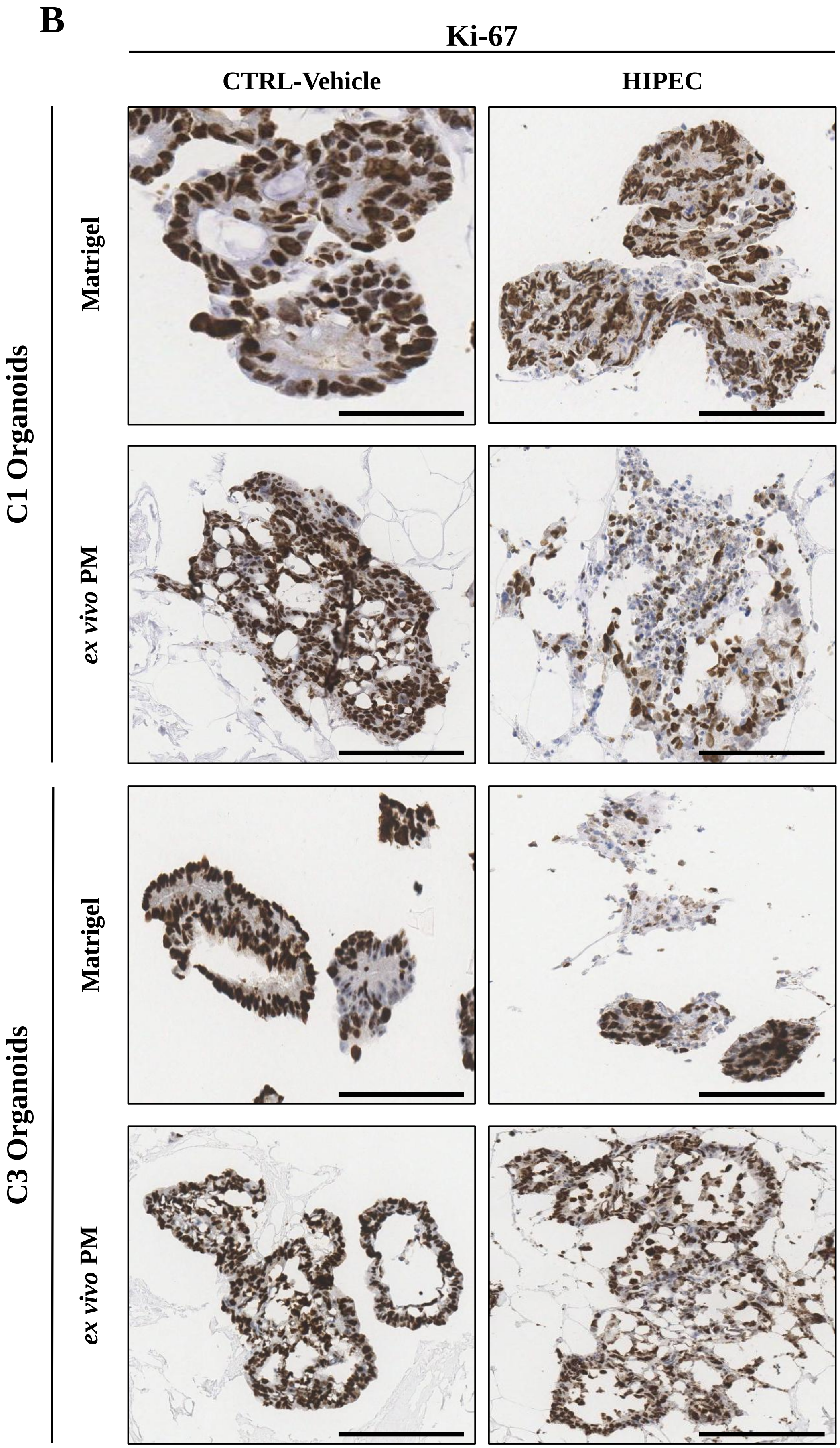


**Supplementary Fig. S7. (B)** Ki-67 immunostaining of C1 and C3 organoids cultured in Matrigel and on neoplastic-derived peritoneal 3D-dECMs after *in vitro* HIPEC treatments. Scale bar: 50 µm.

C

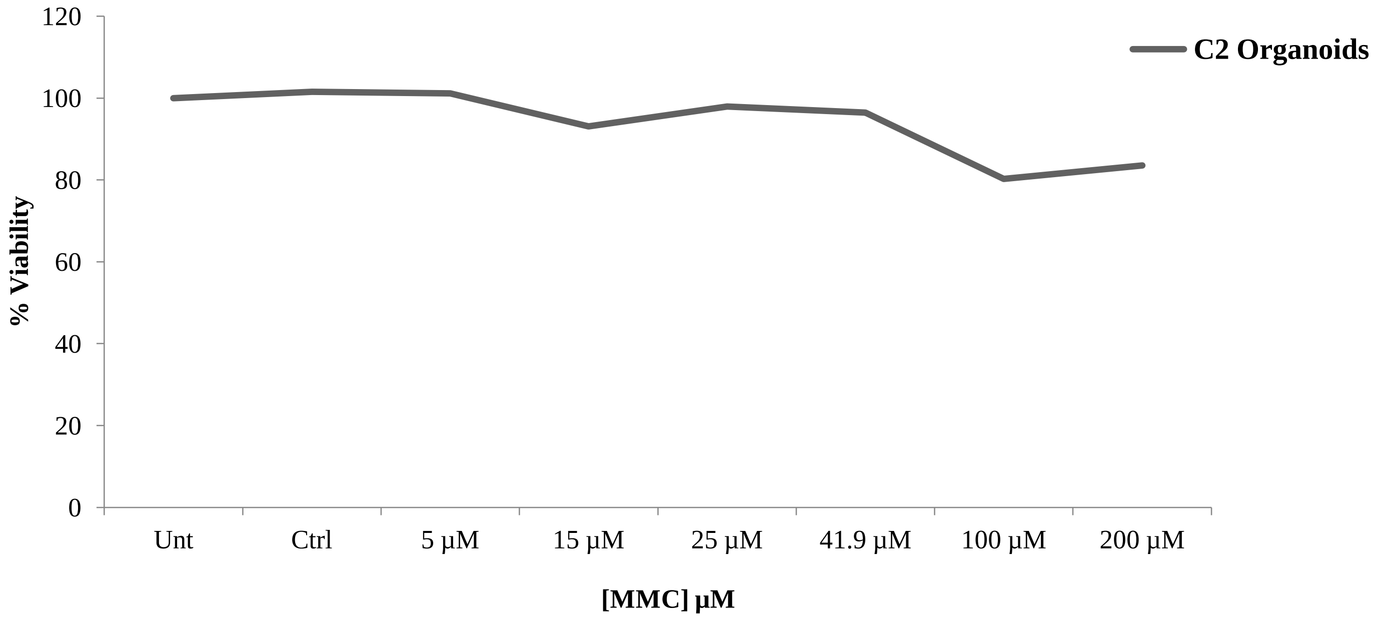


D

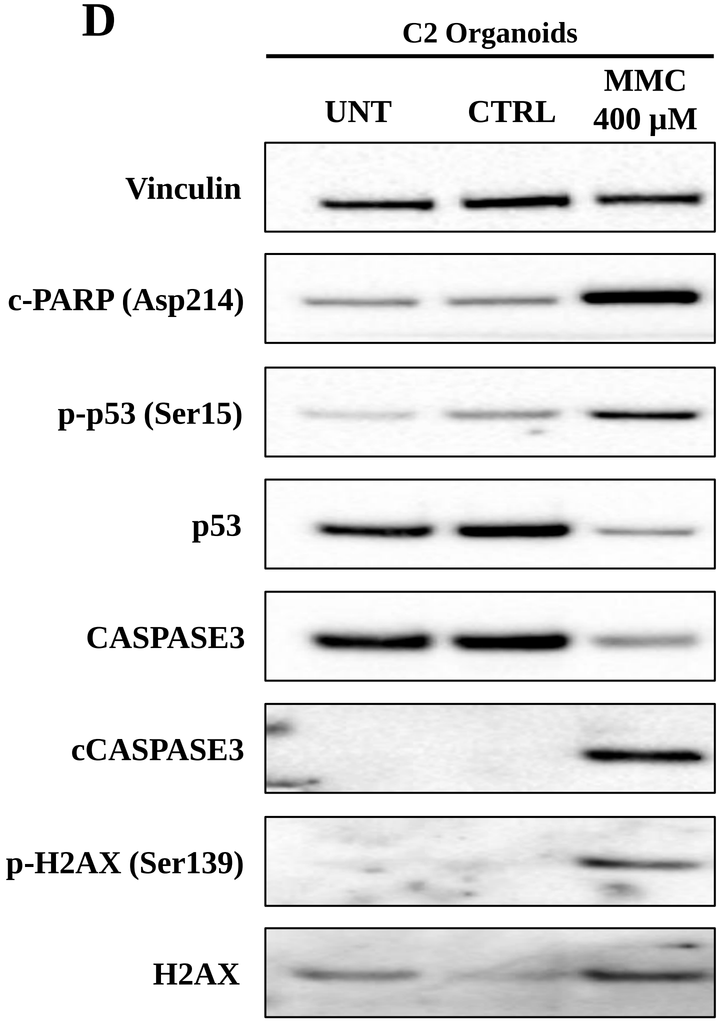


**Supplementary Fig. S7. (C)** Dose response curve of C2 PM-derived organoids cultured in Matrigel and treated with MMC at different concentrations at 42.5 °C for 1 h. **(D)** Immunoblots of cPARP, p-p53, p53, CASPASE3, cCASPASE3, p-H2AX and H2AX in C2 PM-derived organoids treated with MMC 400 µM. Vinculin was used as loading control.

**E**

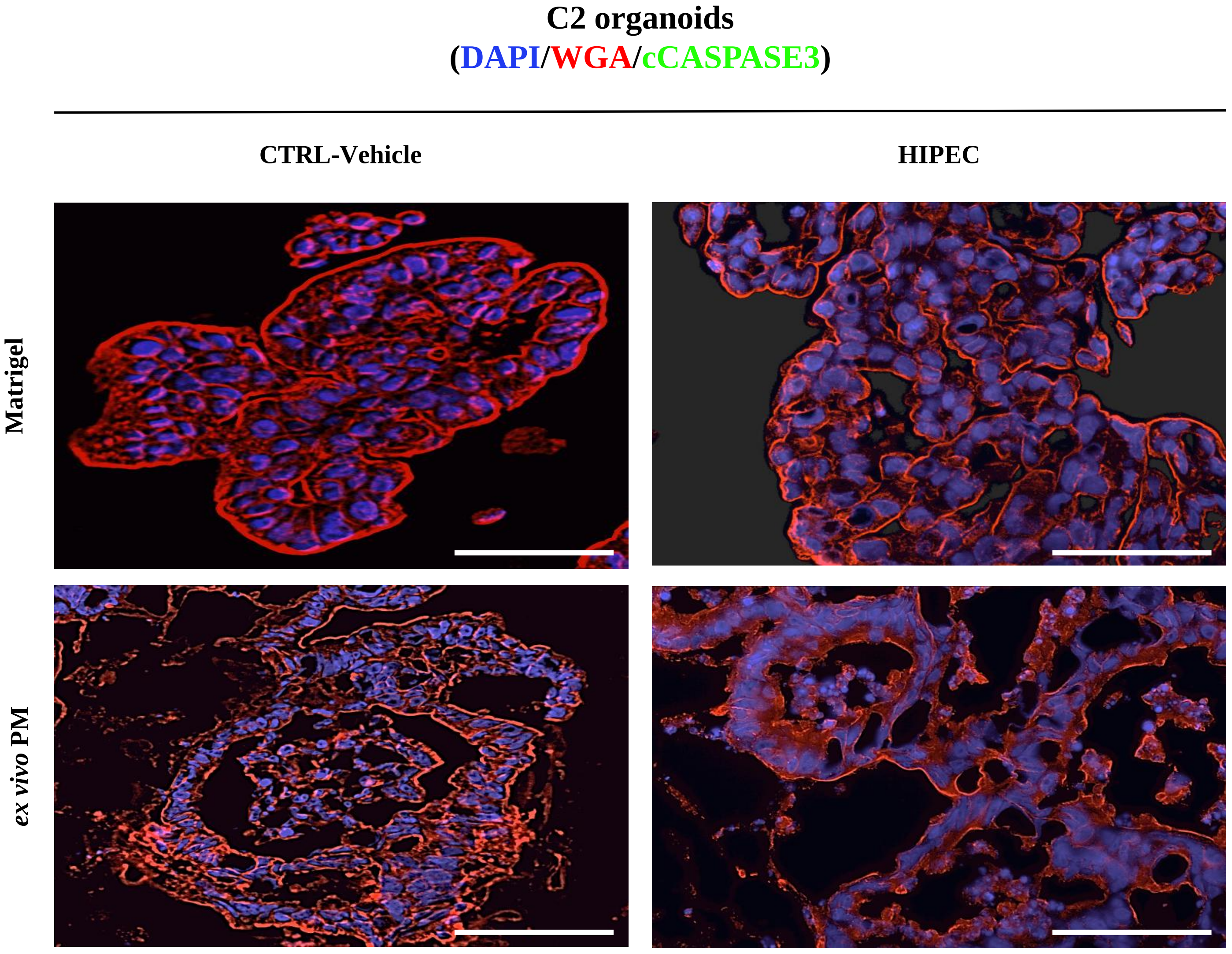


**Supplementary Fig. S7 (E)** IF analysis of C2 PM-derived organoids cultured in Matrigel and on neoplastic-derived peritoneal 3D-dECMs after *in vitro* HIPEC treatments, using cCASPASE3 (green) antibody. The samples were counterstained with WGA (red) and DAPI (blue). Scale bar: 50 µm.

F

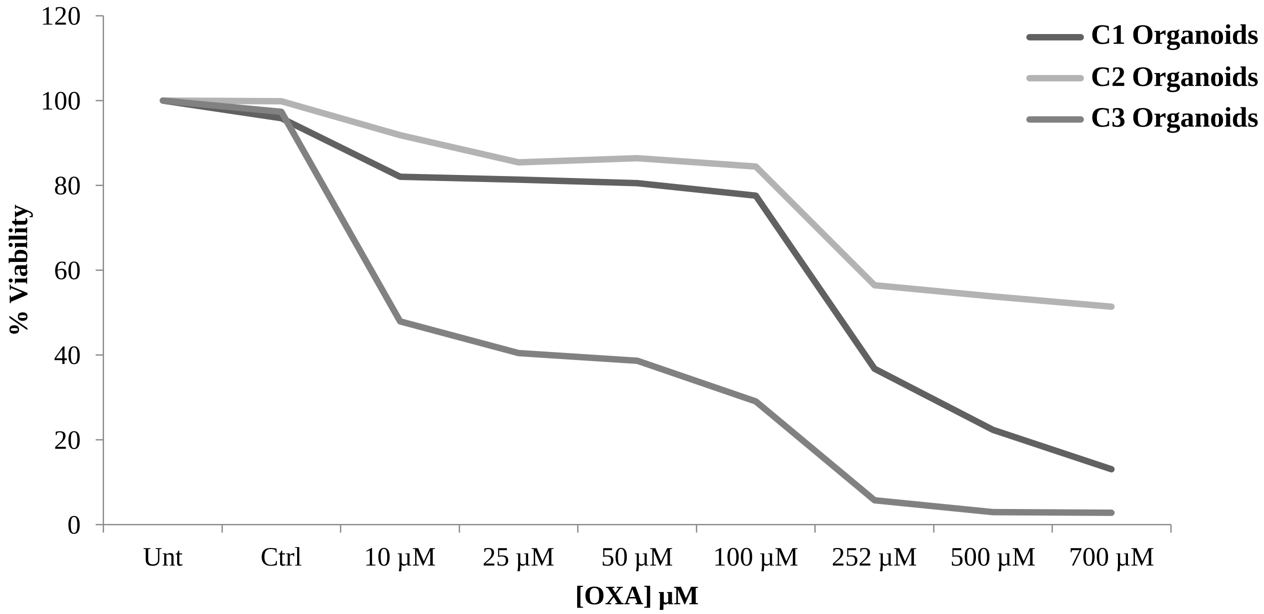


G

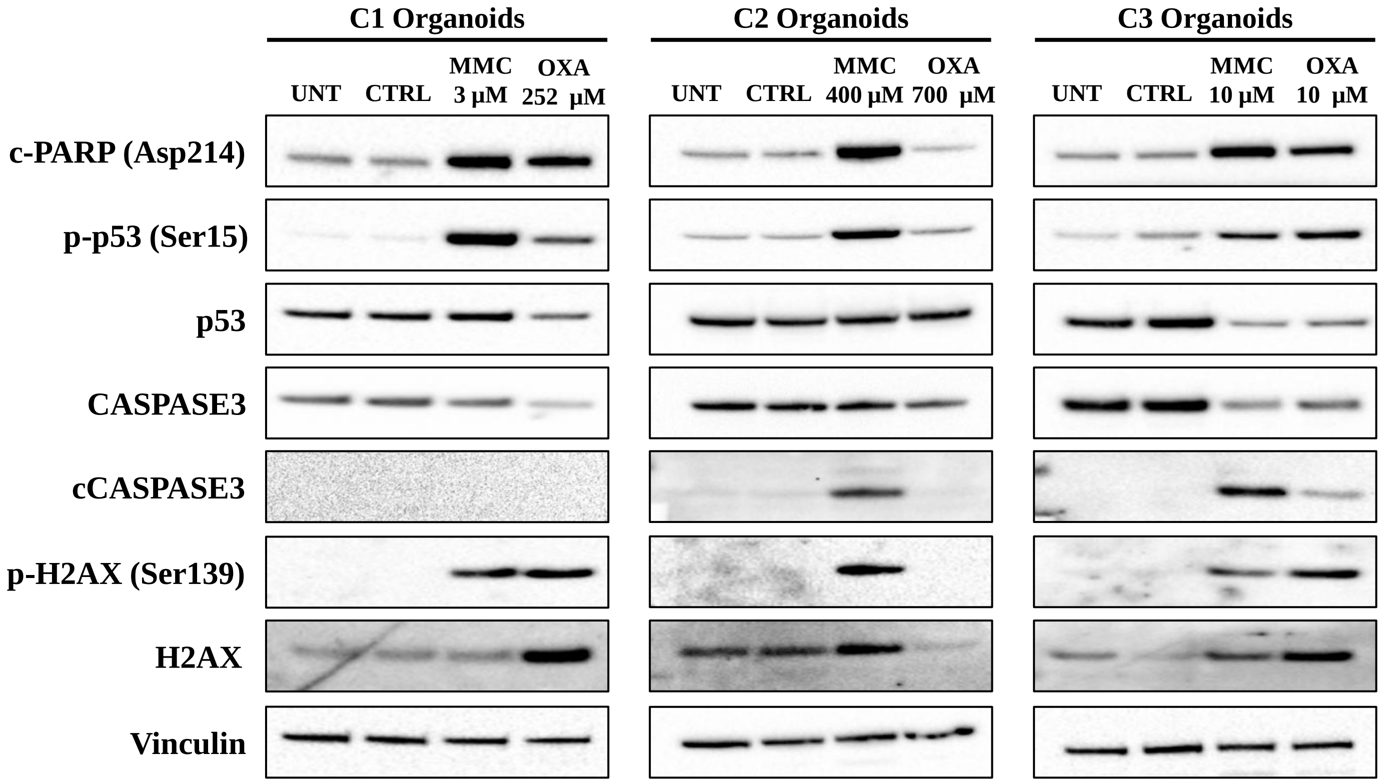


**Supplementary Fig. S7.** (**F**) Dose response curve of C1, C2 and C3 PM-derived organoids cultured in Matrigel and treated with OXA at different concentrations at 42.5 °C for 90 min. **(G)** Immunoblots of cPARP, p-p53, p53, CASPASE3, cCASPASE3, p-H2AX and H2AX in C1, C2 and C3 PM-derived organoids treated with MMC and OXA at the respective $IC_{50}$ concentrations. Vinculin was used as loading control. Ki-67 immunostaining of C2 PM-derived organoids cultured in Matrigel and on neoplastic-derived peritoneal 3D-dECMs after *in vitro* HIPEC treatments.

**H**

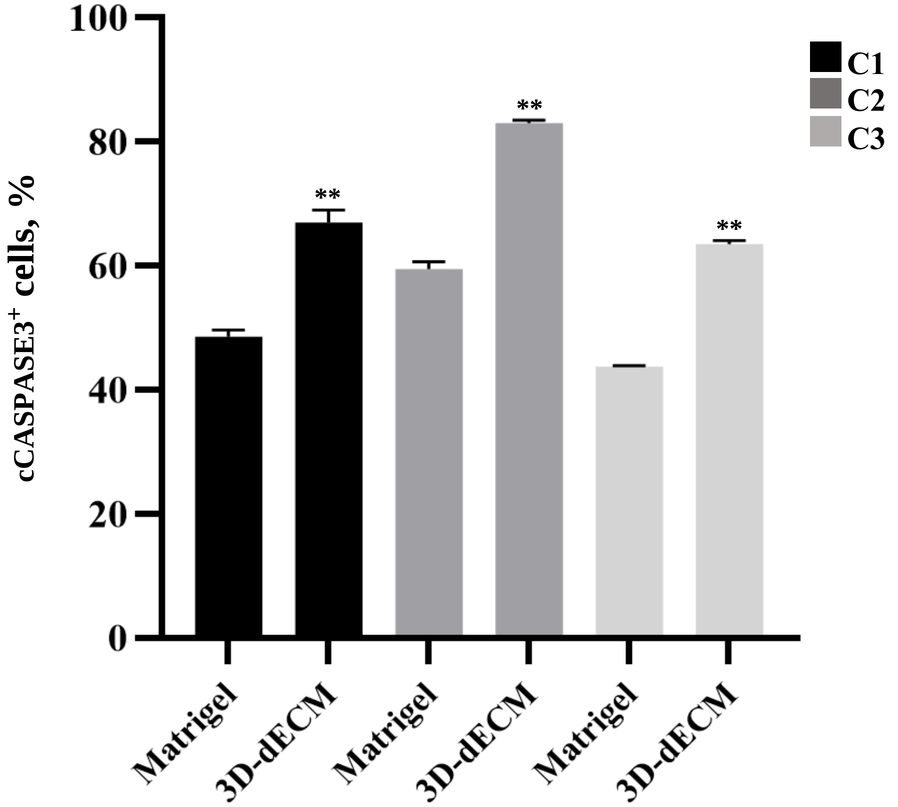


**Supplementary Fig. S7.** (**H**) Percentage of apoptotic TDO treated with OXA, measured as the percentage of cCASPASE-$3^{+}$ cells present in selected fields. Five fields per experiment (40X magnification) were counted. Data are presented as median and SD . One-way ANOVA (**$p$<0.01).
